# Comparative analyses of SAR-CoV2 genomes from different geographical locations and other coronavirus family genomes reveals unique features potentially consequential to host-virus interaction and pathogenesis

**DOI:** 10.1101/2020.03.21.001586

**Authors:** Rahila Sardar, Deepshikha Satish, Shweta Birla, Dinesh Gupta

## Abstract

The ongoing pandemic of the coronavirus disease 2019 (COVID*-*19) is an infectious disease caused by severe acute respiratory syndrome coronavirus 2 (SARS-CoV2). We have performed an integrated sequence-based analysis of SARS-CoV2 genomes from different geographical locations in order to identify its unique features absent in SARS-CoV and other related coronavirus family genomes, conferring unique infection, facilitation of transmission, virulence and immunogenic features to the virus. The phylogeny of the genomes yields some interesting results. Systematic gene level mutational analysis of the genomes has enabled us to identify several unique features of the SARS-CoV2 genome, which includes a unique mutation in the spike surface glycoprotein (A930V (24351C>T)) in the Indian SARS-CoV2, absent in other strains studied here. We have also predicted the impact of the mutations in the spike glycoprotein function and stability, using computational approach. To gain further insights into host responses to viral infection, we predict that antiviral host-miRNAs may be controlling the viral pathogenesis. Our analysis reveals nine host miRNAs which can potentially target SARS-CoV2 genes. Interestingly, the nine miRNAs do not have targets in SARS and MERS genomes. Also, hsa-miR-27b is the only unique miRNA which has a target gene in the Indian SARS-CoV2 genome. We also predicted immune epitopes in the genomes

## Introduction

The first case of COVID-19 patient was reported in December 2019 at Wuhan (China) and then it has spread worldwide to become a pandemic, with maximum death cases in Italy, though initiallythe maximum mortality was reported from China (*1*). According to a WHO report, as on 18^th^ March 2020 there were confirmed 209, 839 COVID-19 cases and 8778 cases of deaths, that includes cases which were locally transmitted or imported(*2*). There are published reports which suggests that SARS-CoV2 shares highest similarity with bat SARS-CoV. Scientists across the globe are trying to elucidate the genome characteristics using phylogenetic, structural and mutational analysis. Recent paper identified specific mutations in receptor binding domain (RBD) domain of spike protein which is most variable part in coronavirus genome(*3*).

There are more than 400 SARS-CoV2 assembled genomes available at NCBI database. Sequence analysis of the genomes can give us plethora of information which can of use for drug development and vaccine development research attempts. In the current work we collected SARS-CoV2 genomes from different geographical origins mainly from India, Italy, USA, Nepal and Wuhan to identify notable genomic features of SARS-CoV2 by integrated analysis. These analyses include identification of notable mutational signatures, host antiviral-miRNA identification and epitope prediction. As a host defense mechanism, a repertoire of host miRNAs also target invading viruses. We followed the parameters used in various anti-viral miRNA databases to predict host anti-viral miRNAs against SARS-CoV2. Our analysis shows unique host-miRNAs targeting SARS-CoV2 virus genes.

## Methodology

### Genome sequences retrieval

Genome sequences of SARS-CoV2 genomes from India, Italy, USA, Wuhan, Nepal, SARS-COV and MERS; with genome NCBI-IDs MT050493.1, MT066156.1, MN985325.1, NC_045512.2, MT072688.1, NC_004718.3 and KC164505.2, respectively, were retrieved from NCBI genome database. SARS-CoV2 genomes from India, Italy, USA, Nepal along with SARS-CoV and MERS were used as query genomes to compare with Wuhan SARS-CoV2 genome. Genes and protein sequences of SARS-CoV2 were retrieved from ViPR database(*4*).

### Coronavirus subtyping and Mutation Analysis

All assembled query genomes in FASTA format were analyzed using Genome Detective Coronavirus Typing Tool (version 1.1.3)(*5*) which allows quick identification and characterization of novel coronavirus genomes. The tool allows submission of up to 2000 sequences and performs analysis within few minutes. This tool has been validated in classification of novel SARS-CoV2 among other coronavirus species (Figure 1 (a)). The tool identifies mutations for each gene of query genomes i.e. SARS-CoV, SARS-CoV2 (India), SARS-CoV2 (Italy), SARS-CoV2 (USA) with respect to Wuhan (China) SAR-CoV2. MSA was performed using online CLUSTAL-OMEGA software.

**Figure 1:**
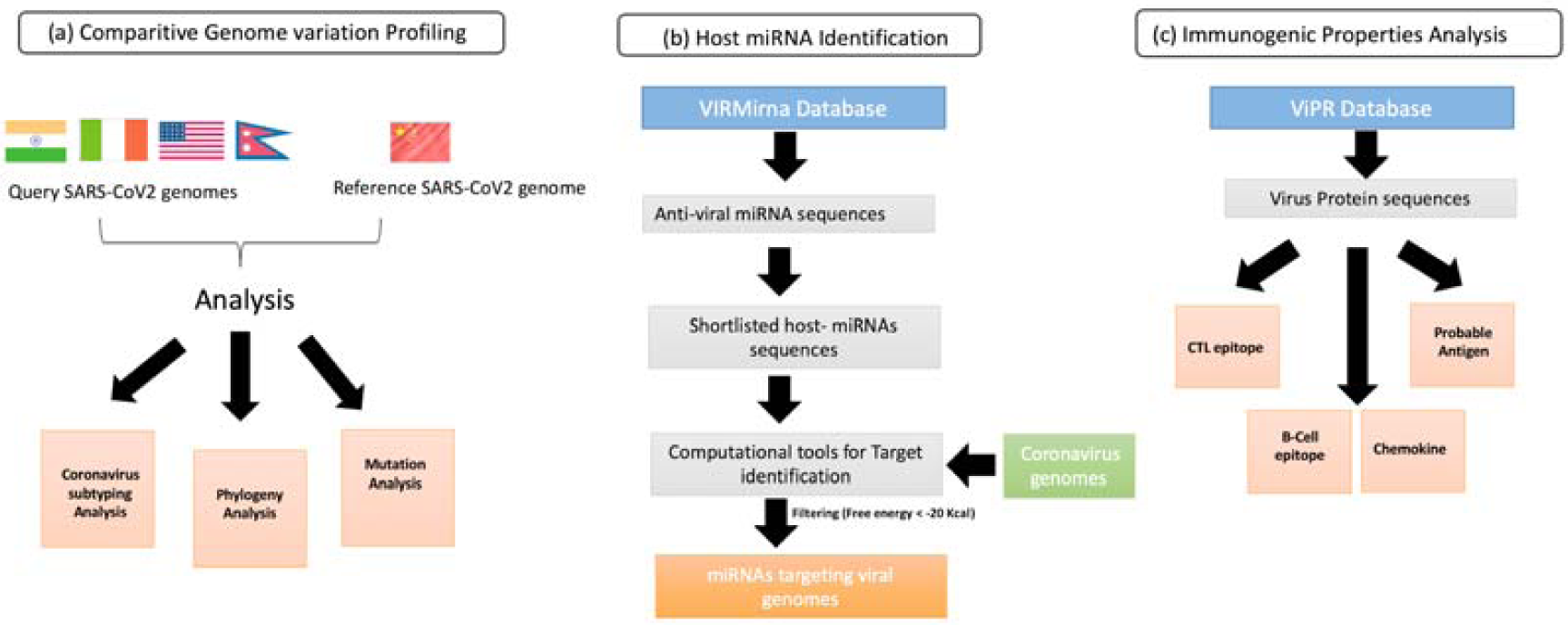
Pipeline used to identify (a) Genome variations in SARS-CoV2 (b) Host anti-viral miRNA (c) Immunogenic property analysis.

### Phylogenetic and Gene Ontology (GO) analysis

To understand the variation in genomes from various geographical areas used in the study, we performed a phylogenetic analysis. Neighbor joining method with bootstrap value of 1000 replicates was used for the construction of consensus tree using MEGA software(*6*) (10.1.7 version). CELLO2GO (*7*)server was used to infer biological function for each protein of SARS-CoV2 genome with their localization prediction. The mutations reported in literature(*3*)were catalogued and evaluated for pathogenicity. We used MutPred(*8*)server to identify disease associated amino acid substitution from neutral substitution, with a p-value of >=0.05. In order to assess the impact of SNPs on protein stability, we used two machine learning based prediction methods. The first method, I-MUTANT server(*9*) was used to predict stability of the protein sequences at pH 7.0 and temperature 25°C. The second prediction method is MuPro(*10*) server, the predictions with the former method helps in getting a consensus prediction.

### Prediction of anti-viral host miRNAs

To predict host miRNAs targeting the virus, we collected a list of experimentally verified antiviral miRNAs with their targets from VIRmiRNA database(*11*). Only these host miRNAs were processed for downstream analysis. (Figure 1(b)) To identify potential host microRNA target sites in the virus genome sequences, we have used miRanda (3.3 a version)(*12, 13*) software, with an energy threshold of −20 kcal/mol. We also used psRNATarget server to compare the predicted targets by the two methods(*14*).

### Immunogenic properties analysis

All the genes and protein sequences for SARS-CoV2 were retrieved from ViPR database. To identify CTL and B-cell epitopes we have used CTLpred(*15*), ABCpred(*16*) servers with default parameters. CHEMOpred(*17*)and Vaxijen server (*18*) were used to predict chemokines and protective probable antigen, respectively (Figure 1(c)).

## Results

### Phylogenetic analysis

Assembled SARS-CoV2 genomes sequences in FASTA format from India, USA, China, Italy and Nepal used for coronavirus typing tool analysis. Using the tool, we were able to locate query SARS-CoV genomes with known SARS-CoV2 to obtain a cladogram for evolutionary analysis as shown in Figure 2. As expected and depicted clearly from the figure, all the SARS-CoV2 genomes are under SARS-CoV2 clade.

**Figure 2:**
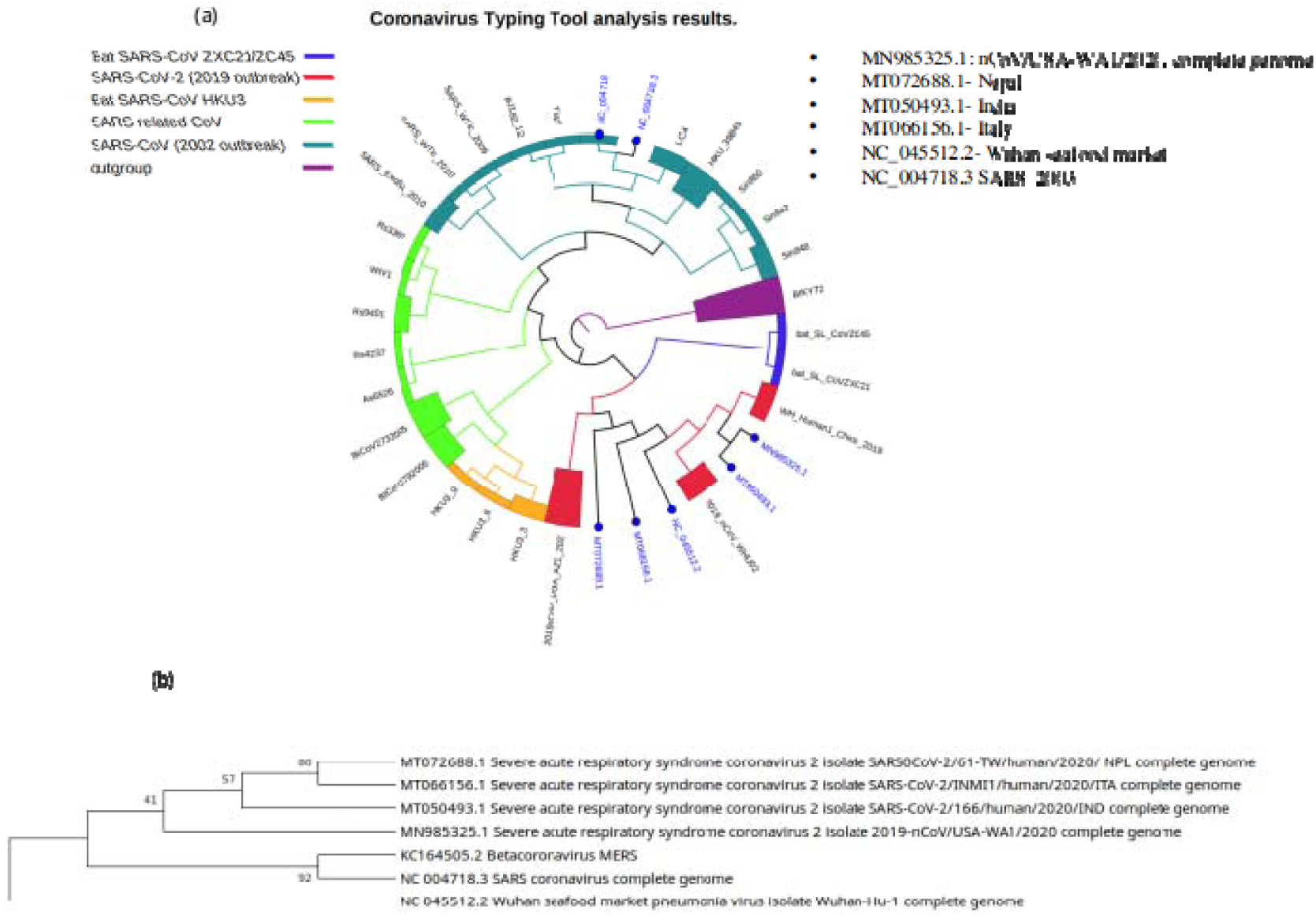
Phylogenetic analysis of SARS-CoV2 genomes from different geographical locations: (a) Genome Detective Coronavirus Subtyping Analysis (b) NJ bootstrap tree for reference SARS-CoV2 genomes

### Genome variation Analysis

From the variation analysis using Genome Detective Coronavirus Subtyping tool, it is found that SARS-CoV2 shares 79.5772 % nucleotide and 83.3672 % amino acid identity with SARS-CoV genome.

The 17% variation in the genomes corresponds to mutations in each protein, including the spike surface glycoprotein, NSP proteins, envelope proteins and the other SARs-CoV proteins (Table S1). We have observed that there are 27, 27, 24 and 17 deletions in orf1ab, orf1a, spike surface glycoprotein and ORF8 (Table1).

**Table 1:** Table showing variation in genes between SARS-CoV2 and SARS-CoV. Variations are shown as I/D/M/F, where I=Indels, D= deletions, M= Misaligned, F= Frameshift

| Variation profiling parameters | SARS-CoV2 vs SARS-CoV (I/D/M/F) |
| --- | --- |
| NT Identity (%) | 79.5772 |
| AA Identity (%) | 83.3672 |
| Number of stop codons | 14 |

| Genes | I/D/M/F |
| --- | --- |
| 1_orf1ab | 4/27/0/0 |
| 2_orf1a | 4/27/0/0 |
| 3_S | 6/24/0/0 |
| 4_ORF3a | 0/1/0/0 |
| 5_E | 1/0/0/0 |
| 6_M | 0/1/0/0 |
| 7_ORF6 | 0/0/0/0 |
| 8_ORF7a | 1/0/0/0 |
| 9_ORF7b | 1/0/0/0 |
| 10_ORF8 | 5/17/1/1 |
| N | 3/0/0/0 |
| ORF9b | 1/0/0/0 |
| ORF10 | 0/0/0/0 |
| ORF14 | 0/0/0/0 |

### Spike surface glycoprotein Mutation and functional Analysis

Several mutations are revealed when SAR-CoV2 and SARS-CoV spike glycoproteins are compared. Six frameshift mutations and 1 insertion in the genome that corresponds to S13_Q14insSDLD (21601_21602insAGTGACCTTGAC) (Table S1) was also revealed. We also observed that there are several mutations located in the regions associated with high immune response (Table S2). These mutations might have significant impact on the antigenic and immunogenic changes, unique to the viruses studied here.

From the sequence alignment we observed that there are mutations in the RBD domain which includes L455Y, F486L, Q493N, S494D and N501T, recently reported by Kristian G. Andersen et. al(*3*). A better understanding of the impact of these mutations on protein function and stability is revealed by MutPred, I-Mutant and MuPro analysis.

From SNPs analysis we observed that all the mutations might bring about decrease in stability without changing their properties i.e. hydrophobicity to hydrophilicity or vice versa. L455Y mutation predicted to altered Ordered interface, Disordered interface Stability, transmembrane protein and gain of GPI-anchor amidation at N450 position (Table 2).

**Table 2:** Predicted effect of reported mutations on protein function and stability using various prediction methods.

| Non-Synonymous SNPs | Type of aa change | Change in hydrophobicity /hydrophilicity due to SNPs | MutPred Hypothesis | MutPred Score | Affected PROSITE and ELM Motifs | I-Mutant | MUpro |
| --- | --- | --- | --- | --- | --- | --- | --- |
| L455Y | aliphatic-aromatic | no | Altered Ordered interface<br>Altered Disordered interface<br>Altered Stability,<br>Altered Transmembrane protein ;<br>Gain of GPI-anchor amidation at N450 | 0.514 | ELME000120,<br>ELME000182 | Decrease | G = -2.3415564 |
| F486L | aromatic-aliphatic | no | NA | 0.36 | NA | Decrease | G = -0.68400097<br>(DECREASE stability) |
| Q493N | polar-polar | no | NA | 0.46 | NA | Decrease | G = -0.54834709<br>(DECREASE stability) |
| N501T | polar-polar | no | NA | 0.36 | NA | Decrease | G =<br>-1.5187308<br>(DECREAS<br>E stability) |
| S494D | polar-polar | no | NA | 0.44 | NA | Decrease | G =<br>-0.2069765<br>(DECREAS<br>E stability) |

It is known, and also confirmed by Gene ontology analysis that the protein is involved in pathogenesis, membrane organization, reproduction, symbiosis, encompassing mutualism through parasitism, and locomotion.

To understand the variations that are occurring with the geographical area we have analyzed Indian, Italian, USA, Nepal SARS-CoV2 genomes (query genomes) compared with Wuhan SARS-CoV2 genome. It was observed that all the genomes shares ∼99% similarity with Wuhan (SARS-CoV2) genome. However, interestingly we also observed that each genome has unique mutations-except the genome from Nepal which shares 100% similarity with the Wuhan genome.

We found that Indian SARS-CoV2 have mutations in orf1ab, nsp2, nsp3, helicase, ORF8 protein and spike surface glycoprotein (Table 3). Spike glycoprotein multiple sequence alignment (S. Figure1) with SARS-CoV2(India), SARS-CoV2 (China, Wuhan) and SARS-Cov(2003) clearly shows the replacement of A to V (24351C>T) at 930 position which are absent in other query genomes namely Italy, USA and Nepal along with SARS-CoV (Table 3). I296V, P1261L and T214I mutations in NSP2, NSP3 and helicase are specific to India SARS-CoV2.

**Table 3:**
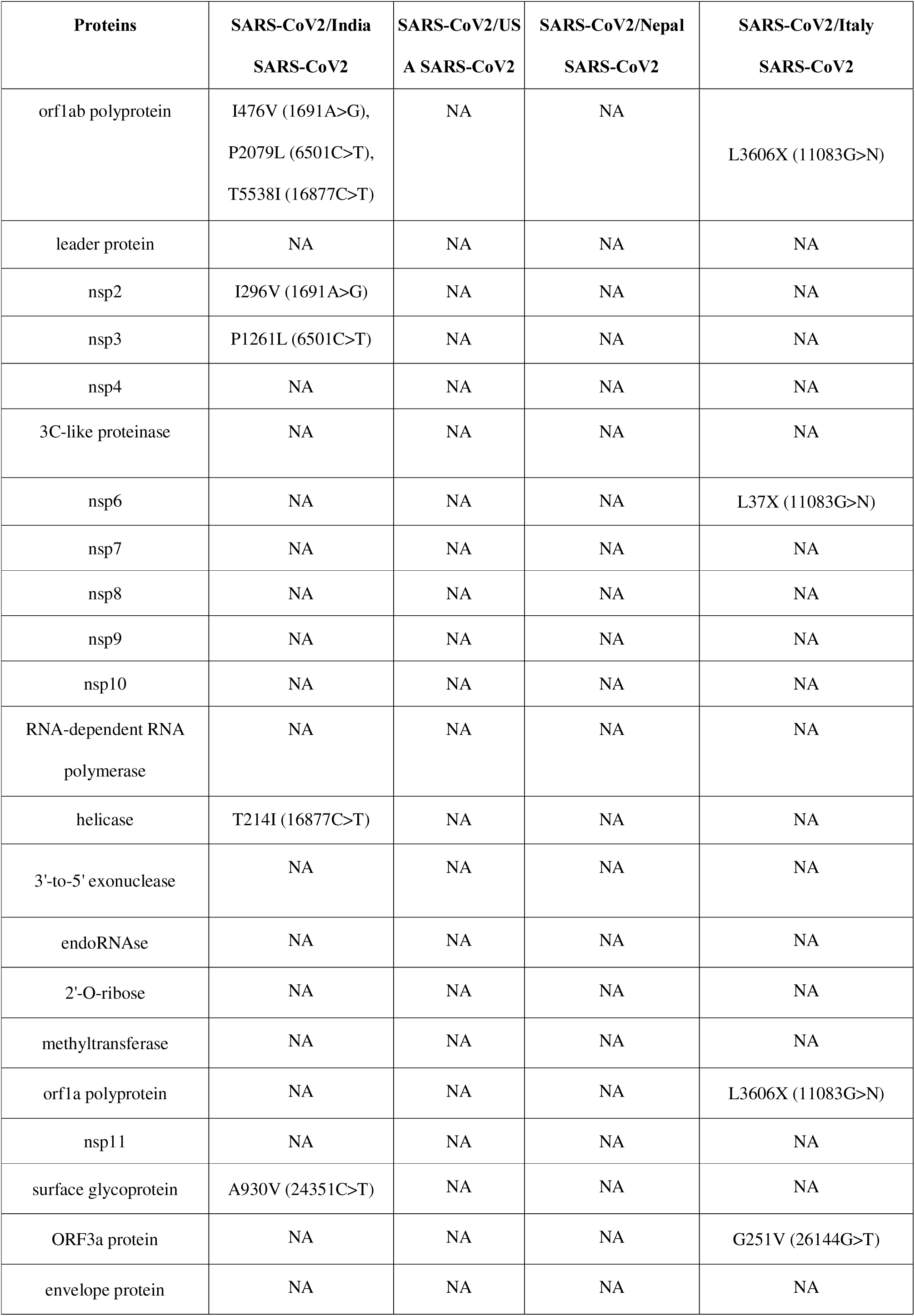
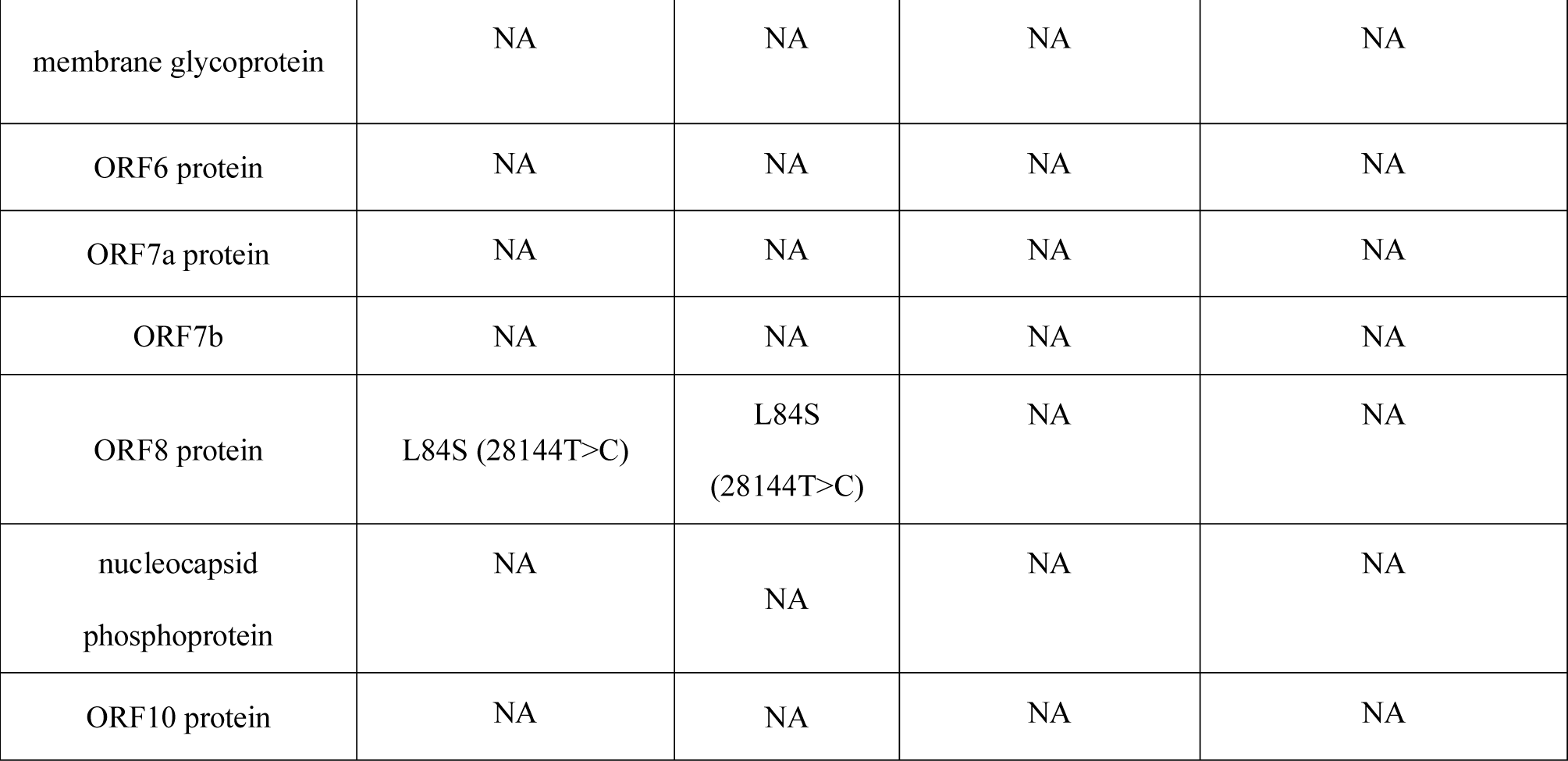
Comparative of SAR-CoV2 genomes variations

While comparing SARS-CoV2 sequence from Italy with that of Wuhan SARS-CoV2, we observed 3 deletions at 3606^th^ and 37^th^ position which correspond to orf1ab and NSP6 protein (Table 3). There is only one mutation, L84S (28144T>C) in ORF8 protein was detected in USA SARS-CoV2 depicting exact similarity with the Wuhan SARS-CoV2.

We have identified 9 miRNAs predicted to specifically target SARS-CoV2 genes. The miRNA targets are absent in SARS and MERS genomes. The miRNAs are hsa-let-7a, hsa-miR101, hsa-miR125a-5p, hsa-miR126, hsa-miR222, hsa-miR23b, hsa-miR378, hsa-miR380-5 and hsa-miR98 which target the SARS-CoV2 genome. These were reported to be targeting Hepatitis C, Herpes simplex virus 1, Hepatitis B, Influenza A, Enterovirus 71 and Vesicular stomatitis virus. The miRNA viral targets predicted by miRanda includes *IFN-B, ATP5B, ERBB2, PB2, PA, NS1, NP, VP1* genes.

PsRNATarget analysis based on the complementary matching between the sRNA sequence and target mRNA sequence with predefined scoring schema identified 6 miRNAs out of the 9 identified miRNAs to target SARS-CoV2 genes. The 6 miRNAsare predicted to act on the viral genomes by cleaving their target sites (Table 4).

**Table 4:** Targets of the six antiviral host miRNAs in SARS-CoV2

| miRNA_Acc. | Inhibition method | Virus Targets gene | Disease associated | Host target gene |
| --- | --- | --- | --- | --- |
| hsa-let-7a | Cleavage | NSP | Hepatitis C virus | IFN beta |
| hsa-miR101 | Cleavage | NSP | Herpes simplex virus 1 | ATP5B |
| hsa-miR126 | Translation | Nucleocapsid | Hepatitis C virus | IFN beta |
| hsa-miR23b | Cleavage | Spike_Glycoprotein | Enterovirus 71 | VP1 |
| hsa-miR378 | Cleavage | Nucleocapsid | Hepatitis C virus | IFN beta |
| hsa-miR98 | Cleavage | Spike_glycoprotein | Hepatitis C virus | IFN beta |

Intriguingly, our analysis (S. Figure 2) revealed that there is only a single host miRNA (hsa-miR-27b) uniquely targeting the India SARS-CoV2.Strikingly, the hsa-miR-27b has been linked to target host IFN-gamma signaling pathways in certain viral infections. Studies on SARS-CoV (30,31) shows that Interferon-beta and interferon-gamma synergistically inhibit the replication of severe acute respiratory syndrome-associated coronavirus. Interestingly, 52 host miRNAs were predicted to have same targets in all the SARS-CoV2 genomes (Wuhan, Italy and USA), except an extra miRNA uniquely targeting Indian genome. Further, we also identified host anti-viral miRNAs predicted to target the spikeglycoprotein of SARS-CoV-2, as shown in Figure 3.

**Figure 3:**
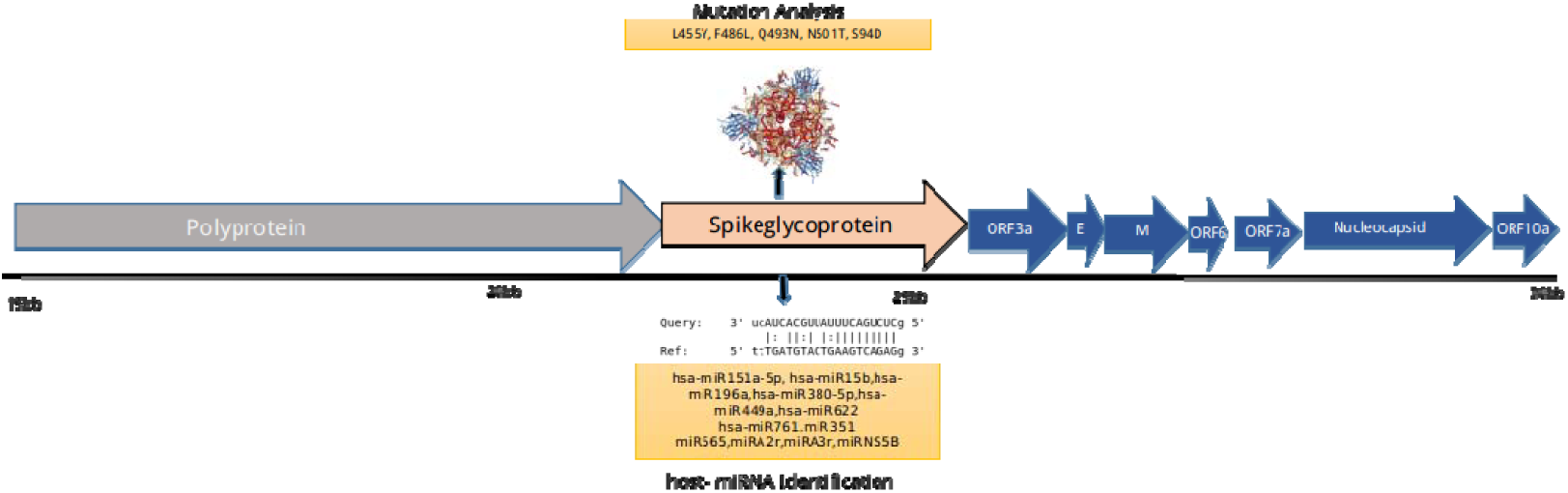
Spike glycoprotein with their host-antiviral miRNAs

### Immunogenic properties of SARS-CoV2 proteins

CTL and B- cell epitopes for each protein of SARS-CoV 2 were determined using CTLpred and ABCPred servers (Table 5).

**Table 5:**
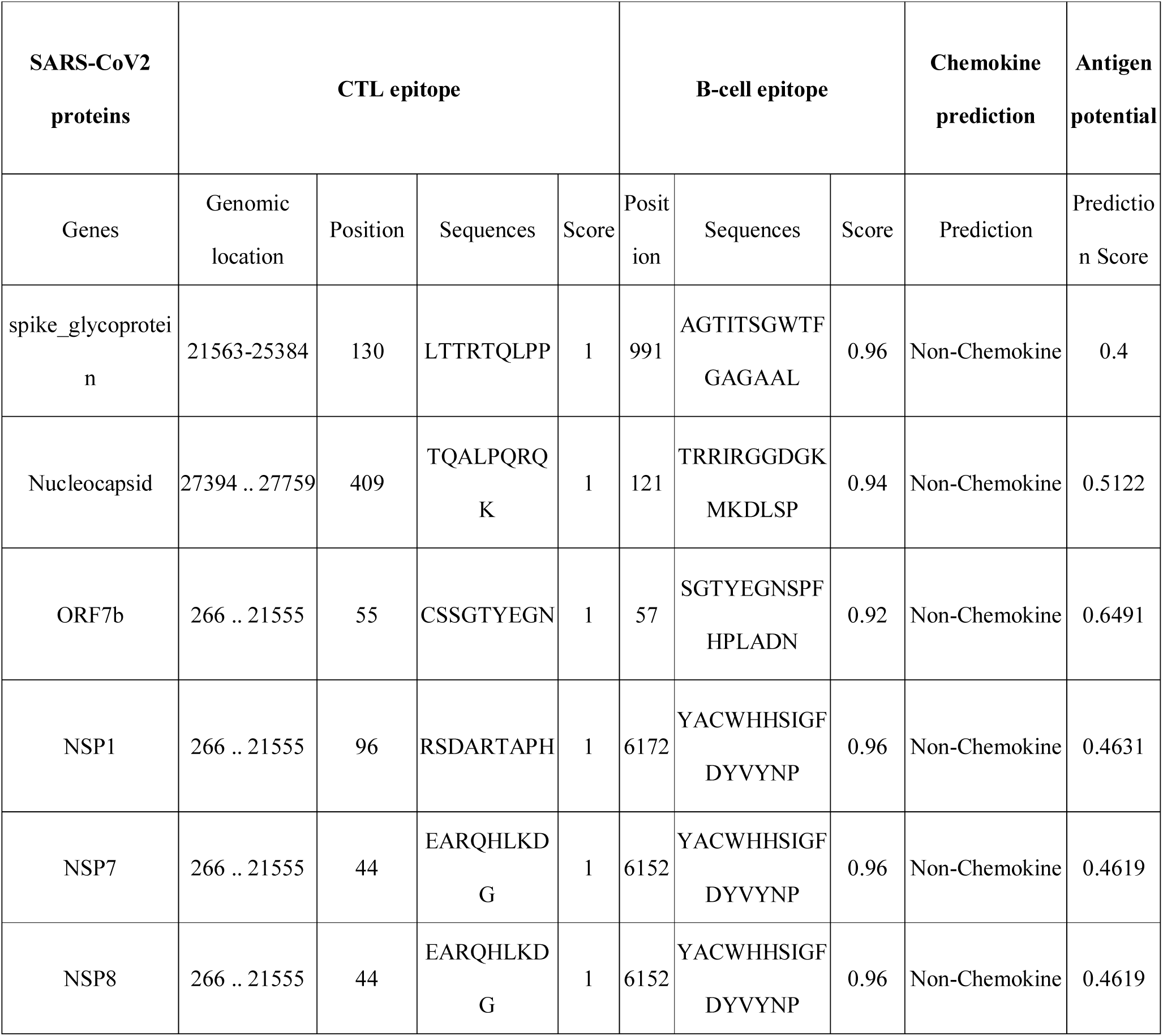
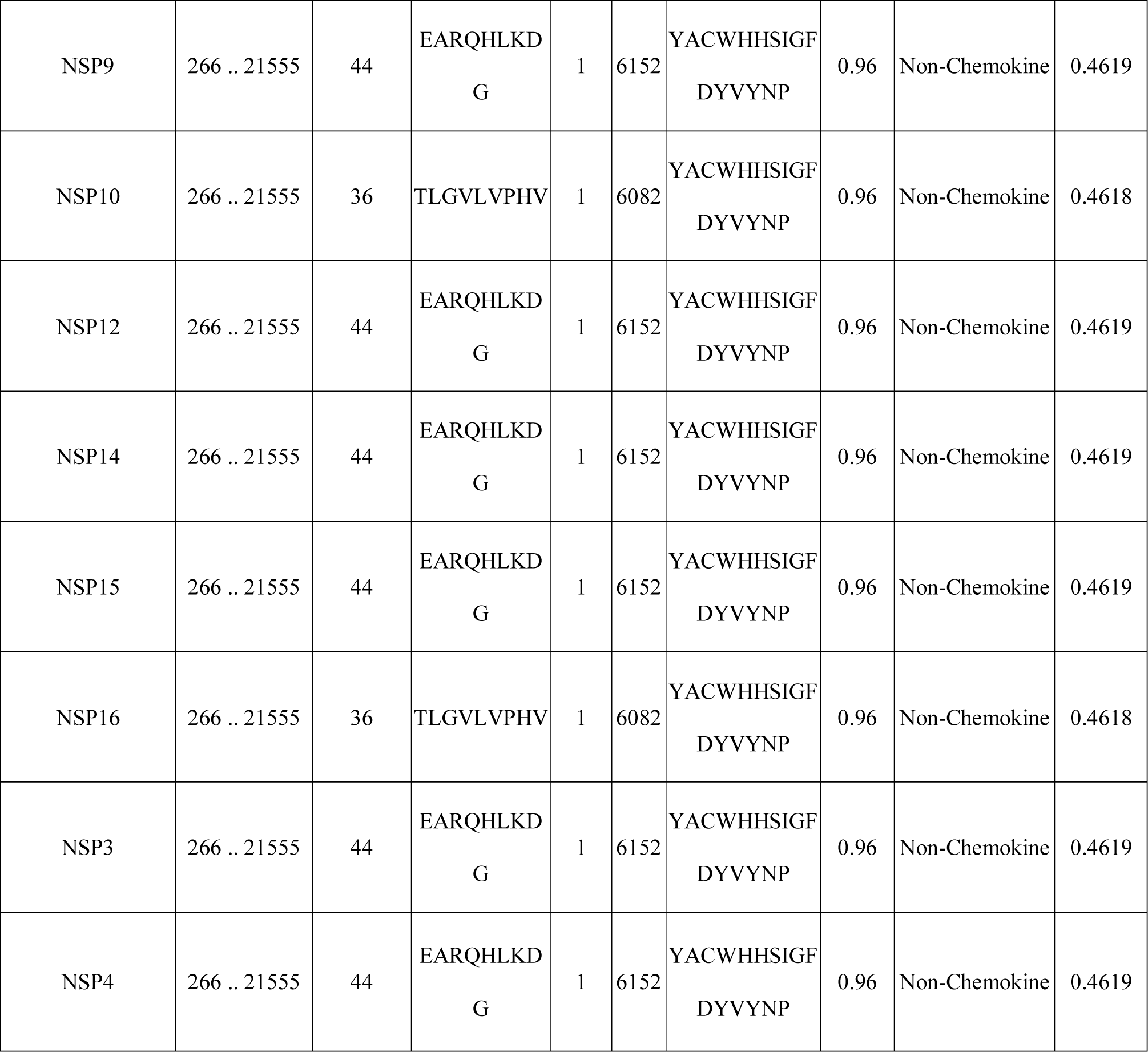
Epitopes (CTL and B) and probable antigen prediction of SARS-CoV genes

Among CTL predictions for spike surface glycoprotein LTTRTQLPP peptide at 130 position was highly antigenic with score 1. Whereas B-cell epitope AGTITSGWTFGAGAAL peptide displayed the highest antigenic score of 0.96 at position 991. Non-structural proteins display similar CTL and B-Cell epitopes i.e YACWHHSIGFDYVYNP and SGTYEGNSPFHPLADN with a prediction score of more than 0.9.

CTL and B-cell epitopes for Nucleocapsid protein are TQALPQRQK and TRRIRGGDGKMKDLSP with the highest predictive score at 121 positions. The highest overall protective antigen prediction probability was observed for ORF7b with a score of 0.64 as reported by VaxiJen server.

## Discussion and conclusion

We have used bioinformatics tools to investigate SARS-CoV2 sequences from different geographical locations. The Phylogenetic analysis of the genomes, the nucleotide sequence diversity analysis of the genomes, the predicted antiviral host miRNAs specific to the genomes and the prediction of immune active sequences in the genomes have yielded some interesting facts, including unique features. For the phylogenetic analysis, we compared the sequences of 6 SARS-CoV2 isolates from different countries namely, Wuhan, India, Italy, USA and Nepal along with other corona virus species (Figure 1). As reported earlier too(*19, 20*), the virus from Wuhan showed higher similarity with SARS-CoV. There was no phylogenetic segregation of the genomes based on geographic origin, whether from the same continent or a neighboring country (Figure 1) but, ambiguously showing varied clustering like Italy and Nepal clustered together, followed by India and USA. This reiterates the findings indicating the massive exchange and importation of the carriers between the epicenter Wuhan and these countries. However, a detail analysis, complemented with more sequences and patient met data will give further evolutionary insights regarding the fast spreading pandemic.

### Genomic comparison

The phylogenetics heterogeneity between different strains is explored by genome variation profiling to find alterations in genetic information during the course of evolution, outbreak, and clinical spectrum caused by the different strains. In case of SARS-CoV2 and SARS-CoV too, few clinical characteristics differentiate them among themselves and with other seasonal influenza infections as well, as reported recently (*21*). Interestingly in the present analysis, in comparison to SARS-CoV, we observed at least one of the variations like indels, deletions, misaligned and frameshift in all the SARS-CoV2 proteins except ORF6, ORF10 and ORF14 (Table S1).

The spread and containment of the SARS-CoV2 outbreak is varying among different countries (https://www.gisaid.org/). Amongst other factors, sequence of the host invasion factors of the virus strain may possibly play an important role in degree of transmission, virulence and pathogenicity of the virus. Therefore, we set out to investigate the genetic variations present in the spike surface glycoprotein (S) which plays a significant role in binding of the virus to the host cell, and determining host tropism and pathogenicity(*22*). Going well with the expectations from a rapidly transmitting pandemic virus, in our analysis, we observed various mutations located in the regions associated with immune response (Table S2). These mutations may have significant impact on the antigenic and immunogenic changes responsible for differences in the severity of the outbreak in different geographical regions.

To gain further insights, we compared the genetic mutation spectrum identified in the four countries, namely USA, Italy, India and Nepal. Surprisingly, the mutation spectrums were different among these countries (Table 3). While sequence from Nepal showed no variations, the maximum number of mutations were identified in Indian sequence-located in ORF1ab, nsp2, nsp3, helicase, ORF8 protein and spike surface glycoprotein (Table3). Probably presence of the unique mutations identified in the genome from Italy are responsible for the sudden upsurge in the number of affected cases and deaths (*2*), combined with other factors-a speculation which maybe verified with more evidences. From this analysis, we also speculate that the presence of country specific mutation spectrum may also be able to explain the current scenario in these countries like severity of illness, containment of the outbreak, the extent and timing of exposures to a symptomatic carrier etc.

Non-structural proteins have their specific roles in replication and transcription (*23*). Previous studies on SARS-CoV revealed Nsp15 as a potential candidate for the therapeutic target(*24*). It is noteworthy to mention that in the present study; various mutations have been identified in all the non-structural proteins suggesting them to be an important and potential player in proposing therapeutic targets and should be explored experimentally.

Many studies have reported that miRNAs not only act as the signature of tissue expression and function but also as potential biomarkers playing important role in regulating disease pathophysiology(*25*). In viral infections, host antiviral miRNAs play a crucial role in the regulation of immune response to virus infection depending upon the viral agent. Many known human miRNAs appear to be able to target viral genes and their functions like interfering with replication, translation and expression. In the present study, we tried to predict the antiviral host-miRNAs specific for SARS-CoV2.Firstly, we compared SARS-CoV2 and SARS-CoV and identified a list of 6 miRNAs unique to SARS-CoV2 and previously reported to be associated with other virus diseases, including HIV (Table. 4).

Clinical investigations have suggested that patients with cardiac diseases, hypertension, or diabetes, who are treated with ACE2-increasing drugs including inhibitors and blockers show increased expression of ACE2 and thus are at higher risk of getting the SARS-CoV2 infection (*26*). Also there are studies on the regulatory role of miRNA hsa-mir-27b-3p described in ACE2 Signaling (*27*). The results of the present study suggest a strong correlation between miRNA hsa-mir-27b-3p and ACE2 which needs to be confirmed experimentally in SARS-CoV2 cases.

Further, we tried to compare the miRNAs in the genomes and observed some striking findings. We observed that out of all the miRNAs, hsa-miR-27b is the only unique miRNA specific to India SARS-CoV2 and showed no significant complementarity-based nucleotide-binding with the strain of SARS-CoV2 from other countries. This is surprising and is of utmost importance along with our other novel finding of a unique mutation identified in the spike surface glycoprotein (A930V (24351C>T)) in the Indian sequence. The target of the hsa-miR-27b sequence is the location containing A930V mutation in the spike surface glycoprotein which perhaps, should be explored further, experimentally. Reports have demonstrated the ability of miR-27b to inhibit HIV-1 replication (28). There are contradicting reports regarding the use of HIV antiviral regimen for SARS-CoV2 infection showing no effects in China (29) while the Indian Council of Medical Research (India) has given the guidelines for the use of HIV treatment in these cases as some of the patients showed improvement with the administration. A few other drugs, including FDA approved antiviral and antimalarials are being used with significant claimed-successes. Needless to say, the variation in the protein sequences may have some influence on the therapeutic effects of these drugs.

Based on our analysis, we speculate an important regulatory role of miR-27b in SARS-CoV2 infection. The contradictory treatment outcomes may be due to the presence of the miR-27b target in the Indian genome specifically. It probably indicates that the specific genetic and miRNA spectrum should be considered as the basis of the treatment management.

The findings in the study have revealed unique features of the SARS-CoV2 genomes, which may be explored further. For example, one may analyse the link between severity of diseases to each of the variants, expression of the predicted host antiviral miRNAs can be checked in the patients, the predicted epitopes may be explored for their immunogenicity, difference in treatment outcomes may also be correlated with genome variations, lastly the potential of the unique segments of the virus proteins and the unique host miRNAs may be explored in development of novel antiviral therapies.

## Acknowledgements

This work was financially supported by the Department of Biotechnology (DBT), Government of India, grant BT/BI/25/066/2012, awarded to D.G. Financial support provided by the Indian Council of Medical Research (ICMR), India to RS as Senior Research Fellowship is duly acknowledged (2019-5850). DS received fellowship from the Council of Scientific and industrial Research (CSIR, India, 09/0512(0207)/2016/EMR-1). SB is the recipient of National Post-doctoral fellowship from SERB, Department of Science & Technology, India (PDF/2017/001326).

## Supplementary figures

**S. Figure1:**
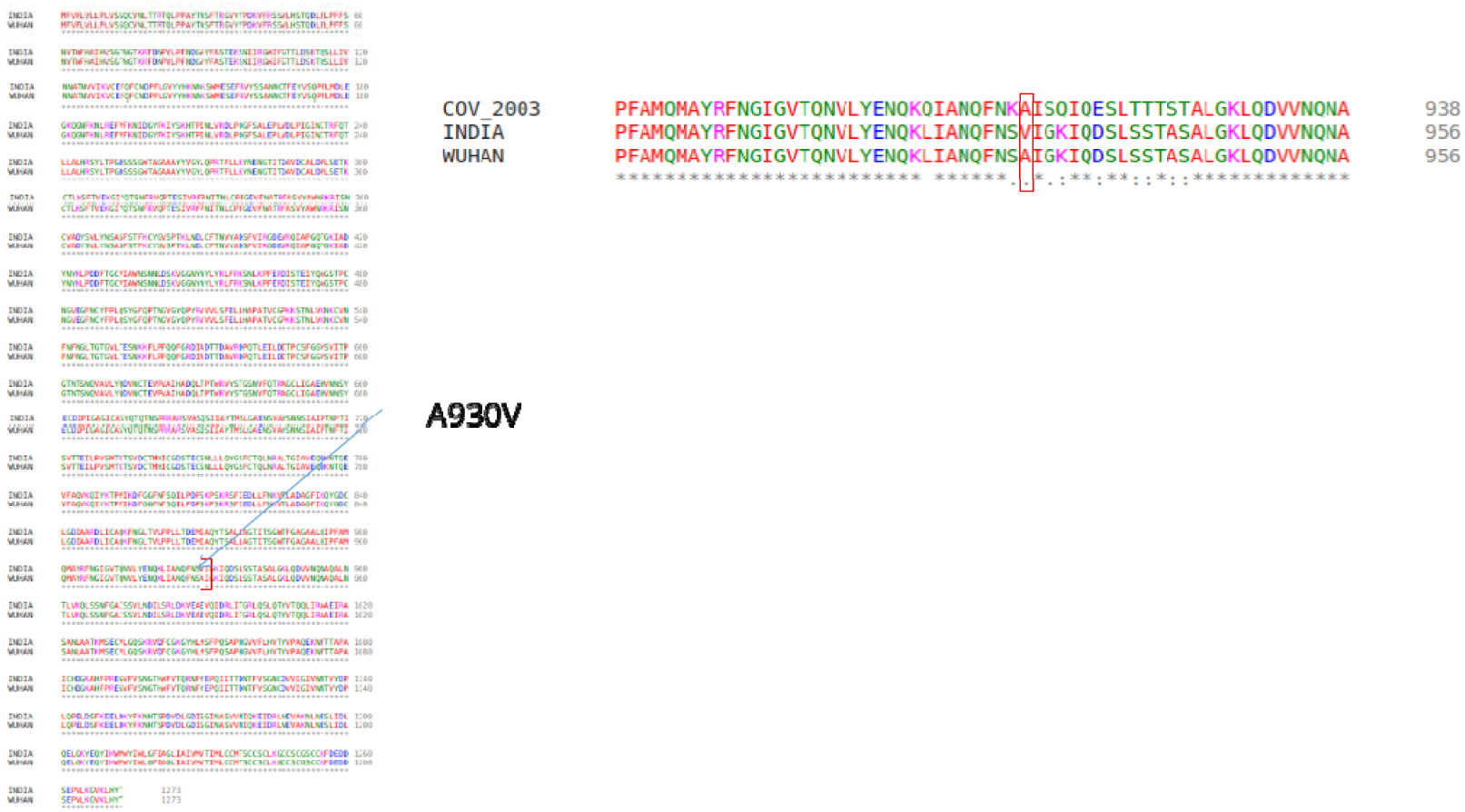
Sequence alignments of SARS-CoV2 spike glycoprotein sequences from Indian, Chinese (Wuhan) and SARS (right pane only) genomes. The alignments show mutation at 930 position in the Indian SARS-CoV2 genome. SARS and SARS-CoV2 sequences are conserved, except mutation at position 930 in the Indian SARS-CoV2

**S. Figure2:**
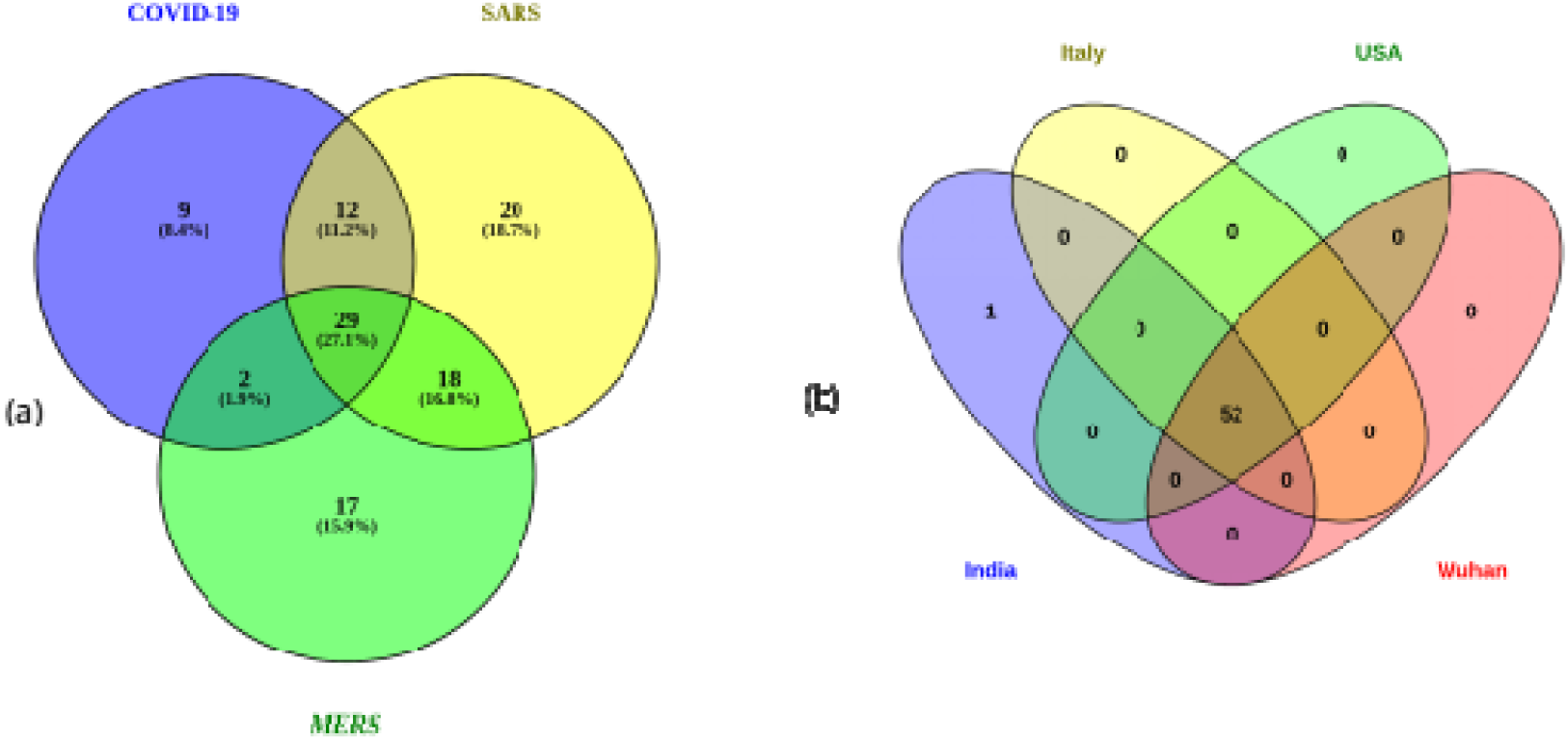
Venn diagram showing (a) Host-antiviral miRNAs in COVID-19, SARS and MERS viruses. 9 miRNAs are predicted to uniquely target COVID-19 genes (b) Host-antiviral miRNAs targeting SARS-CoV2 genes in the virus genomes from different geographical locations. There is one miRNAs uniquely targeting the Indian SARS-CoV2 genes only.

