## Supplementary Table S2 for "Comparative analyses of SAR-CoV2 genomes from different geographical locations and other coronavirus family genomes reveals unique features potentially consequential to host-virus interaction and pathogenesis"

*** Red color highlighted residues are regions which are found to be associated with high immune response**

*** Blue color regions are the reported mutations in RBD domain**

**Spike:**

V3I (21569G>A), V6L (21578G>T 21580T>A), L7F (21583A>T), P9T (21587C>A 21589A>T), V11T (21593G>A 21594T>C 21595C>T), S13G (21599A>G),

S13_Q14insSDLD (21601_21602insAGTGACCTTGAC), Q14R (21603A>G), V16T (21608G>A 21609T>C 21610T>C), N17T (21612A>C), L18F (21614C>T),

T19D (21617A>G 21618C>A 21619A>T), T20D (21620A>G 21621C>A 21622C>T), R21del (21623_21625delAGA), T22V (21626A>G 21627C>T), L24del

(21632_21634delTTA), P25A (21635C>G 21637C>T), A27N (21641G>A 21642C>A 21643A>T), T29_N30insQH (21649_21650insCAACAT), N30T

(21651A>C), F32S (21657T>C 21658C>T), T33M (21660C>T 21661A>G), K41E (21683A>G), V42I (21686G>A), S46D (21698T>G 21699C>A 21700A>C),

V47T (21701G>A 21702T>C), H49Y (21707C>T), S50L (21711C>T), F59Y (21738T>A), W64G (21752T>G), A67T (21761G>A), H69_T73del

(21767_21781delCATGTCTCTGGGACC), G75H (21785G>C 21786G>A), K77_R78del (21790_21795delTAAGAG), D80G (21801A>G 21802T>C), L84I

(21812C>A), N87K (21823T>G), V90I (21830G>A), S94A (21842T>G), I100V (21860A>G 21862A>T), I101V (21863A>G 21865A>C), I105V (21875A>G),

T108S (21884A>T), L110M (21890T>A 21892A>G), D111N (21893G>A 21895T>C), S112N (21896T>A 21897C>A 21898G>C), T114S (21902A>T

21904C>A), L117V (21911C>G 21913A>G), L118I (21914C>A), V120I (21920G>A), A123S (21929G>T), K129R (21947A>C 21948A>G), V130A (21951T>C

21952C>A), E132N (21956G>A 21958A>C), Q134E (21962C>G), F135L (21967T>G), N137D (21971A>G 21973T>C), D138N (21974G>A 21976T>C), L141F

(21985G>T), G142A (21987G>C), Y144S (21993A>C), Y145K (21995T>A 21997C>A), H146P (21999A>C), K147M (22002A>T 22003A>G), N148_S151del

(22004_22015delAACAACAAAAGT), W152G (22016T>G 22018G>T), M153T (22020T>C 22021G>A), E154Q (22022G>C 22024A>G), S155T (22026G>C

22027T>A), E156H (22028G>C 22030G>T), F157T (22031T>A 22032T>C 22033C>T), R158M (22035G>T 22036A>G), V159I (22037G>A 22039T>A), Y160F

(22041A>T 22042T>C), S161D (22043T>G 22044C>A), S162N (22047G>A), N164F (22052A>T 22053A>T), V171I (22073G>A 22075C>A), Q173D

(22079C>G 22081G>T), P174A (22082C>G 22084T>C), L176S (22088C>T 22089T>C 22090T>G), M177L (22091A>C 22093G>T), L179V (22097C>G),

E180S (22100G>T 22101A>C), G181E (22104G>A), Q183S (22109C>T 22110A>C 22111G>A), N188H (22124A>C 22126T>C), I197K (22152T>A

22153T>A), Y200F (22161A>T), F201L (22163T>C 22165T>C), K202Y (22166A>T 22168A>T), I203V (22169A>G 22171A>T), S205K (22175T>A 22176C>A

22177T>G), K206G (22178A>G 22179A>G 22180G>C), H207Y (22181C>T 22183C>T), T208Q (22184A>C 22185C>A 22186G>A), N211D (22193A>G),

L212V (22196T>G), Q218S (22214C>T 22215A>C 22216G>T), S221N (22223T>A 22224C>A 22225G>C), A222T (22226G>A), E224K (22232G>A), L226I

(22238T>A 22240G>T), V227F (22241G>T 22243A>T), D228K (22244G>A 22246T>G), I231L (22253A>C 22255A>T), R237N (22272G>A 22273G>T),

Q239R (22277C>A 22278A>G), T240A (22280A>G 22282T>C), L241I (22283T>A 22285A>T), A243T (22289G>A 22291T>A), L244A (22292T>G 22293T>C

22294A>C), H245_Y248del (22295_22306delCATAGAAGTTAT), L249F (22309G>T), T250S (22310A>T 22312T>A), G252A (22317G>C), D253Q (22319G>C

22321T>A), S254D (22322T>G 22323C>A 22324T>C), S255I (22325T>A 22326C>T), S256_G257del (22328_22333delTCAGGT), T259G (22337A>G

22338C>G 22339A>C), A260T (22340G>A 22342T>G), G261S (22343G>T 22344G>C 22345T>A), Y266F (22359A>T), Q271K (22373C>A 22375A>G),

R273T (22380G>C 22381G>T), L276M (22388C>A 22390A>G), N280D (22400A>G), A292S (22436G>T 22438A>T), L293Q (22440T>A 22441T>A), D294N

(22442G>A 22444C>T), S297A (22451T>G 22453A>T), T299L (22457A>C 22458C>T 22459A>C), T302S (22466A>T 22468G>T), L303V (22469T>G

22471G>T), T307E (22481A>G 22482C>A 22483T>G), V308I (22484G>A 22486A>T), E309D (22489A>C), Q321V (22523C>G 22524A>T 22525A>T), T323S

(22529A>T), E324G (22533A>G), S325D (22535T>G 22536C>A), I326V (22538A>G), R346K (22599G>A), A348P (22604G>C 22606A>T), N354E (22622A>G

22624C>G), R357K (22632G>A), A372T (22676G>A), S373F (22680C>T 22681A>T), P384A (22712C>G 22714T>C), T393S (22739A>T 22741T>C), I402V

(22766A>G 22768T>C), R403K (22770G>A 22771A>G), E406D (22780A>T), K417V (22811A>G 22812A>T 22813G>T), T430M (22851C>T 22852A>G),

I434L (22862A>C 22864A>T), S438T (22874T>A), N439R (22878A>G 22879C>G), L441I (22883C>A), S443A (22889T>G), K444T (22893A>C 22894G>T),

V445S (22895G>T 22896T>C 22897T>A), G446T (22898G>A 22899G>C), L452K (22916C>A 22917T>A 22918G>A), **L455Y** (22926T>A 22927G>T), F456L

(22928T>C), K458H (22934A>C 22936G>T), S459G (22937T>G 22938C>G 22939T>C), N460K (22942T>G), K462R (22947A>G 22948A>G), T470N

(22971C>A), E471V (22974A>T 22975A>G), I472P (22976A>C 22977T>C 22978C>T), Y473F (22980A>T 22981T>C), Q474S (22982C>T 22983A>C

22984G>C), A475P (22985G>C 22987C>T), G476D (22989G>A), S477G (22991A>G), T478K (22995C>A), N481T (23004A>C 23005T>C), G482P

(23006G>C 23007G>C 23008T>A), V483del (23009_23011delGTT), E484P (23012G>C 23013A>C 23014A>T), G485A (23016G>C), **F486L** (23018T>C),

F490W (23031T>G 23032T>G), **Q493N** (23039C>A 23041A>T), **S494D** (23042T>G 23043C>A 23044A>T), Q498Y (23054C>T 23056A>C), P499T

(23057C>A), **N501T** (23064A>C), V503I (23069G>A), H519N (23117C>A), K529L (23147A>T 23148A>T 23149G>A), N532D (23156A>G 23158T>C), V534I

(23162G>A), K537Q (23171A>C 23173A>G), E554P (23222G>C 23223A>C 23224G>T), N556S (23228A>T 23229A>C 23230C>A), K558R (23235A>G

23236G>A), L560Q (23241T>A 23242G>A), I569V (23267A>G), A570S (23270G>T), T572F (23276A>T 23277C>T 23278T>C), A575S (23285G>T

23287T>C), Q580K (23300C>A 23302G>A), L582S (23306C>T 23307T>C), T588S (23324A>T), S591A (23333T>G), T604A (23372A>G), N606S (23378A>T

23379A>C 23380C>T), Q607E (23381C>G 23383G>A), E619D (23419A>T), P621S (23423C>T), V622T (23426G>A 23427T>C 23428T>A), T632A

(23456A>G), V635I (23465G>A 23467T>A), S640N (23480T>A 23481C>A 23482T>C), R646Q (23499G>A 23500T>A), N657D (23531A>G), N658T

(23535A>C 23536C>T), Q675H (23587G>T), Q677V (23591C>G 23592A>T 23593G>T), T678S (23594A>T), N679L (23597A>T 23598A>T 23599T>A),

S680_R683del (23601_23612delCTCCTCGGCGGG), A684L (23601_23612delCTCCTCGGCGGG 23613C>T), V687T (23621G>A 23622T>C 23623A>T),

A688S (23624G>A 23625C>G 23626T>C), S689Q (23627A>C 23628G>A 23629T>A), Q690K (23630C>A), I693V (23639A>G 23641T>G), E702D

(23668A>T), N703S (23670A>G), V705I (23675G>A), S711T (23693T>A 23695T>C), T719S (23717A>T 23719T>A), V722I (23726G>A), I726V (23738A>G

23740T>A), L727M (23741C>A 23743A>G), T732A (23756A>G 23758C>T), T739N (23778C>A 23779A>T), S750A (23810A>G 23811G>C 23812C>T),

T768S (23864A>T 23866T>A), V772A (23877T>C), K776R (23888A>C 23889A>G 23890A>C), Q779R (23898A>G 23899A>T), I788M (23926T>G), P793T (23939C>A 23941A>T), I794L (23942A>T 23944T>G), D796Y (23948G>T), S810L (23990T>C 23991C>T), S813T (24000G>C 24001C>T), I834M (24064C>G), D839E (24079T>A), A845N (24095G>A 24096C>A), E868D (24166A>T), Q872A (24176C>G 24177A>C 24178A>C), S875A (24185T>G), L878V(24194T>G 24196A>T), A879S (24197G>A 24198C>G 24199G>T), I882A (24206A>G 24207T>C), S884A (24212T>G), L922Q (24326T>C 24327T>A24328G>A), S929K (24348G>A 24349T>G), G932S (24356G>A 24358C>T), K933Q (24359A>C), D936E (24370C>A), S939T (24377T>A 24379T>A), S940T

(24380T>A 24382C>A), A942S (24386G>T), S943T (24390G>C), S1055A (24725T>G), A1070S (24770G>T 24772A>C), K1073R (24780A>G), D1084E

(24814T>A), H1088Y (24824C>T), S1097F (24852C>T 24853A>T), H1101S (24863C>T 24864A>C 24865C>T), V1104I (24872G>A 24874A>T), Y1110F

(24891A>T), E1111S (24893G>T 24894A>C 24895A>T), V1133I (24959G>A 24961C>T), I1216V (25208A>G), M1233L (25259A>T), C1247A (25301T>G

25302G>C 25303T>A)
