## Supplementary Table S1 for "Comparative analyses of SAR-CoV2 genomes from different geographical locations and other coronavirus family genomes reveals unique features potentially consequential to host-virus interaction and pathogenesis"

### Results Summary

#### Name: NC_004718.3

| | |
|---|---|
| **Length** | 29751 |
| **Species** | Severe acute respiratory syndrome-related coronavirus |
| **NT Identity (%)** | 79.5772 |
| **AA Identity (%)** | 83.3672 |
| **Genome** | 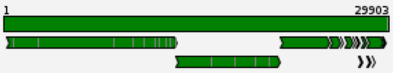 |

### Detailed Results of NC_004718.3

#### Sequence Assignment

| | |
|---|---|
| **Length** | 29751 |

##### Assignment

| | |
|---|---|
| **Type** | Severe acute respiratory syndrome-related coronavirus (Taxonomy ID: 694009) |
| **Reference Genome** | NC_045512.3 (Length: 29903bp) |
| **Host(s)** | Homo sapiens / Paguma larvata |
| **NT Identity (%)** | 79.5772 |
| **AA Identity (%)** | 83.3672 |
| **NT Quality** | 1.15879 |
| **AA Quality** | 5.92939 |
| **Number of stop codons** | 14 |
| **Number of CDS** | 14 |

##### Alignment

| | |
|---|---|
| **Alignment score** | 34641 (NT) + 84980 (AA) = 119621 |
| **Concordance (%)** | 75.9503 |
| **Alignment method** | Global, seeded, nucleotide + amino acids (AGA) |

##### Genome region

Sequence starts at position 1 and ends at position 29894 relative to NC_045512.3 reference sequence.

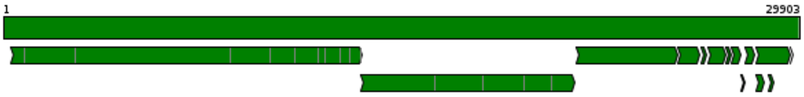


##### Alignment Detailed statistics

| | Begin | End | Coverage | Score | Concordance | Matches | Identities | I/D/M/F* | Stop Codons |
|---|---|---|---|---|---|---|---|---|---|
| **NT** | 1 | 29894 | 99.9% | 34641.0 | 58.8% | 29674 (99.0%) | 23675 (79.0%) | 77/220 | |
| **CDS** | | | | | | | | | |
| 1_orf1ab | 1 | 7097 | 100% | 44033.0 | 89.3% | 7070 (99.6%) | 6125 (86.3%) | 4/27/0/0 | 1 |
| 2_orf1a | 1 | 4406 | 100% | 25296.0 | 84.3% | 4379 (99.3%) | 3552 (80.5%) | 4/27/0/0 | 1 |
| 3_S | 1 | 1274 | 100% | 7176.0 | 82.1% | 1250 (97.7%) | 974 (76.1%) | 6/24/0/0 | 1 |
| 4_ORF3a | 1 | 276 | 100% | 1490.0 | 76.7% | 275 (99.6%) | 200 (72.5%) | 0/1/0/0 | 1 |

| | Begin | End | Coverage | Score | Concordance | Matches | Identities | I/D/M/F* | Stop Codons |
|---|---|---|---|---|---|---|---|---|---|
| 5_E | 1 | 76 | 100% | 447.0 | 94.3% | 76 (98.7%) | 73 (94.8%) | 1/0/0/0 | 1 |
| 6_M | 1 | 223 | 100% | 1419.0 | 92.9% | 222 (99.6%) | 202 (90.6%) | 0/1/0/0 | 1 |
| 7_ORF6 | 1 | 62 | 100% | 300.0 | 75.0% | 62 (100%) | 42 (67.7%) | 0/0/0/0 | 0 |
| 8_ORF7a | 1 | 122 | 100% | 758.0 | 89.6% | 122 (99.2%) | 105 (85.4%) | 1/0/0/0 | 1 |
| 9_ORF7b | 1 | 44 | 100% | 252.0 | 79.7% | 44 (97.8%) | 36 (80.0%) | 1/0/0/0 | 1 |
| 10_ORF8 | 1 | 120 | 99.2% | 117.0 | 15.6% | 104 (82.5%) | 37 (29.4%) | 5/17/1/1 | 0 |
| 11_N | 1 | 420 | 100% | 2641.0 | 92.1% | 420 (99.3%) | 382 (90.3%) | 3/0/0/0 | 1 |
| 12_ORF9b | 1 | 98 | 100% | 451.0 | 73.0% | 98 (99.0%) | 72 (72.7%) | 1/0/0/0 | 1 |
| 13_ORF10 | 1 | 39 | 100% | 212.0 | 78.8% | 39 (100%) | 32 (82.1%) | 0/0/0/0 | 2 |
| 14_ORF14 | 1 | 74 | 100% | 388.0 | 77.3% | 74 (100%) | 57 (77.0%) | 0/0/0/0 | 2 |
| **Proteins** | | | | | | | | | |
| orf1ab polyprotein (YP_009724389.1) | 1 | 7097 | 100% | 44033.0 | 89.3% | 7070 (99.6%) | 6125 (86.3%) | 4/27/0/0 | 1 |

| Begin | End | Coverage | Score | Concordance | Matches | Identities | I/D/M/F* | Stop Codons |
|---|---|---|---|---|---|---|---|---|

P6L (282C>T), F8V (287T>G), V38A (378T>C), Q44E (395C>G), D48N (407G>A), V56L (431G>C 433T>G), R77L (494C>T 495G>T), T78S (498C>G 499T>C), A79T (500G>A 502A>C), P80N (503C>A 504C>A), V84K (515G>A 516T>A 517T>G), M85V (518A>G 520G>C), L92M (539C>A 541C>G), E93D (544A>C), E102I (569G>A 570A>T 571G>A), I114T (606T>C 607A>C), V116I (611G>A 613G>T), K120N (625G>T), A138I (677G>A 678C>T), F143Y (693T>A), Y154I (725T>A 726A>T), F157Y (735T>A), Q158E (737C>G), E159Q (740G>C), S166G (761A>G), V169A (771T>C 772T>A), T170L (773A>C 774C>T), M174T (786T>C 787G>T), Y182V (809T>G 810A>T), E199D (862G>T), L204F (875C>T), A211S (896G>T 898T>A), S212M (899T>A 900C>T 901A>G), F221Y (927T>A 928T>C), D223E (934C>G), T224S (935A>T 937T>G), E233D (964A>C), Y241F (987A>T), E246D (1003A>T), L251H (1016T>C 1017T>A 1018G>C), L259S (1040T>A 1041T>G 1042G>T), N267K (1066T>A), N272K (1081T>G), I280K (1104T>A), I281V (1106A>G), T283V (1112A>G 1113C>T 1114T>C), L293T (1142C>A 1143T>C), D294E (1147T>G), N310Q (1193A>C 1195T>G), Q314N (1205C>A 1207A>T), C316H (1211T>C 1212G>A), D324N (1235G>A), G327D (1245G>A), T329V (1250A>G 1251C>T), G334C (1265G>T), V337L (1274G>C 1276T>G), F343H (1292T>C 1293T>A), T350V (1313A>G 1314C>T), K351I (1317A>T 1318A>T), A354P (1325G>C 1327C>T), Q362T (1349C>A 1350A>C 1351A>T), I368M (1369T>G), Y369P (1370T>C 1371A>C 1372T>A), H374Q (1387C>A), N375D (1388A>G 1390T>C), S376P (1391T>C), V378I (1397G>A 1399A>T), L384V (1415C>G), E386D (1423A>T), E390H (1433G>C 1435A>C), G392N (1439G>A 1440G>A), L393I (1442T>A 1444G>T), K394E (1445A>G), I396R (1451A>C 1452T>G 1453T>A), I404R (1476T>G 1477T>A), A405C (1478G>T 1479C>G 1480C>T), S412A (1499T>G 1501T>C), H417Y (1514C>T), C420R (1523T>C), N430D (1553A>G 1555C>T), C433S (1563G>C 1564T>A), N434G (1565A>G 1566A>G), V438I (1577G>A), V439T (1580G>A 1581T>C), E441D (1588A>C), G442N (1589G>A 1590G>A), S443V (1592T>G 1593C>T 1594C>G), G445T (1598G>A 1599G>C 1600T>C), D448E (1609C>G), N449D (1610A>G 1612C>T), Q455S (1628C>A 1629A>G 1630A>T), K456R (1631A>C 1632A>G 1633A>T), K458R (1637A>C 1638A>G 1639A>T), K468H (1667A>C 1669A>T), I473V (1682A>G 1684C>T), V488I (1727G>A 1729G>T), E489D (1732A>C), V491I (1736G>A 1738G>A), G493S (1742G>A), A498S (1757G>T 1759A>T), Q501T (1766C>A 1767A>C 1768A>C), F509Y (1791T>A), A516P (1811G>C 1813T>C), K517V (1814A>G 1815A>T), E525Q (1838G>C), K527R (1845A>G), I529V (1850A>G 1852A>T), S531T (1857G>C 1858T>A), Y534C (1866A>G), A535G (1869C>G 1870A>T), A537P (1874G>C 1876A>C), E539Q (1880G>C), R542G (1889C>G), V544I (1895G>A 1897A>C), S549A (1910T>G 1912C>G), E553D (1924A>T), T554A (1925A>G 1927T>A), Q556N (1931C>A 1933A>C), N557H (1934A>C 1936T>C), V559I (1940G>A 1942G>T), R560P (1944G>C), V561D (1947T>A), K564R (1956A>G 1957G>A), I567V (1964A>G 1966A>C), Q575E (1988C>G 1990G>A), Y576Q (1991T>C 1993T>G), I581V (2006A>G 2008T>C), M585V (2018A>G 2020G>T), F586Y (2022T>A 2023C>T), A591L (2036G>C 2037C>T 2038T>C), N594S (2046A>G), L595V (2048C>G 2050A>C), V596I (2051G>A), V597I (2054G>A 2056A>T), I601V (2066A>G 2068T>A), V605L (2078G>C), L608Q (2087T>C 2088T>A), T614S (2105A>T), I616L (2111A>C 2113C>T), F617L (2116T>G), V620T (2123G>A 2124T>C), Y621V (2126T>G 2127A>T), K625R (2139A>G 2140A>G), V627I (2144G>A), L628F (2147C>T), D629E (2152T>A), L631I (2156C>A), E633A (2163A>C), F635L (2168T>C), K636S (2172A>G 2173G>T), E637A (2175A>C), R643K (2193G>A 2194A>G), G645A (2199G>C), V649L (2210G>C 2212T>C), I652L (2219A>C), S653I (2222T>A 2223C>T 2224A>T), C655G (2228T>G), A656V (2232C>T), C657F (2235G>T), E658D (2239A>C), G661K (2246G>A 2247G>A 2248T>G), V665Q (2258G>C 2259T>A 2260C>G), T666V (2261A>G 2262C>T 2263C>T), C667A (2264T>G 2265G>C), A668S (2267G>T), K669D (2270A>G 2272G>T), E670N (2273G>A 2275A>C), E673D (2284G>T), S674C (2285A>T), Q676K (2291C>A 2293G>A), T677C (2294A>T 2295C>G 2296A>C), F679I (2300T>A), K680D (2303A>G 2305G>T), L681V (2306C>G), F685A (2318T>G 2319T>C 2320T>A), A687E (2325C>A 2326T>A), L688M (2327T>A), A690I (2333G>A 2334C>T), S692Q (2339T>C 2340C>A 2341T>A), I693V (2342A>G), I694T (2346T>C), G696A (2352G>C), K701R (2366A>C 2367A>G), A702S (2369G>T 2371C>A), T708V (2387A>G 2388C>T 2389A>C), V710I (2393G>A), T711A (2396A>G 2398G>T), H712Q (2401C>A), K719Q (2420A>C), V721I (2426G>A 2428T>A), K722R (2429A>C 2430A>G 2431A>T), S723G (2432T>G 2433C>G), R724K (2436G>A 2437A>G), E726Q (2441G>C 2443A>G), T727L (2444A>C 2445C>T 2446T>G), G728Q (2447G>C 2448G>A 2449C>A), I739V (2480A>G 2482T>A), I740T (2484T>C), E745D (2500A>T), T746S (2501A>T), L747H (2505T>A), P748D (2507C>G 2508C>A), E750V (2514A>T), V751L (2516G>C 2518G>T), L752T (2519T>A 2520T>C 2521A>C), T753S (2522A>T 2524A>T), T760N (2544C>A 2545T>C), D762E (2551T>A), Q764E (2555C>G), P765A (2558C>G), Q768T (2567C>A 2568A>C 2569A>G), T770V (2573A>G 2574C>T), S771D (2576A>G 2577G>A), E772S (2579G>A 2580A>G 2581A>C), A773F (2582G>T 2583C>T 2584T>C), V774T (2585G>A 2586T>C 2587T>A), E775N (2588G>A 2590A>T), A776G (2592C>G 2593T>A), P777A (2594C>G 2596A>T), L778I (2597T>A 2599G>C), I785V (2618A>G 2620T>A), T796K (2652C>A), K798Q (2657A>C 2659G>A), A803S (2672G>T 2674A>T), N805G (2678A>G 2679A>G), M806L (2681A>T 2683G>A), M807L (2684A>C), V808A (2688T>C 2689A>T), T812V (2699A>G 2700C>T), T814R (2705A>C 2706C>G 2707A>C), T821I (2727C>T 2728A>T), K822_V823insG (2731_2732insGGT), D827E (2746T>A), I831W (2756A>T 2757T>G 2758A>G), S838N (2778G>A), N840R (2784A>G 2785T>A), I849V (2810A>G), A859V (2841C>T), L864S (2855C>T 2856T>C), N869T (2871A>C), D877E (2896T>G), I880V (2903A>G 2905A>G), E888D (2929A>T), P892N (2939C>A 2940C>A 2941A>C), L893M (2942C>A), M902V (2969A>G 2971G>A), Y905F (2979A>T), E910D (2995G>T), S911A (2996T>G), F914E (3005T>G 3006T>A 3007T>A), K915N (3010A>C), L916F (3013G>T), A917S (3014G>T 3016T>A), H919R (3021A>G), D930E (3055T>A), E933D (3064A>C), G934D (3066G>A), G934_D935insA (3067_3068insGCA), D935E (3070T>G), F941I (3086T>A), E942D (3091G>T), P943E (3092C>G 3093C>A), S944T (3095T>A 3097A>C), T945C (3098A>T 3099C>G), Q946E (3101C>G), Y947H (3104T>C), K958L (3137A>C 3138A>T 3139A>C), T965S (3158A>T 3160T>A), S966A (3161T>G), A967E (3165C>A 3166T>A), A968T (3167G>A 3169T>A), L969V (3170C>G), Q970R (3174A>G), P971V (3176C>G 3177C>T), Q975E (3188C>G), D983_T999del (3212_3262delGATAGTCAACAAACTGTTGGTCAACAAGACGGCAGTGAGGACAATCAGACA), I1002E (3269A>G 3270T>A 3271T>G), T1004S (3275A>T), I1005_V1006del (3278_3283delATTGTT), V1008I (3287G>A), Q1009E (3290C>G 3292A>G), Q1011E (3296C>G), L1012P (3299T>C 3300T>C), E1013_M1014del (3302_3307delGAGATG), L1016P (3312T>C), V1019E (3321T>A 3322T>A), V1020E (3324T>A 3325T>A), Q1021_I1023del (3327_3335delAGACTATTG), E1024P (3327_3335delAGACTATTG 3336A>C), S1027Q (3344A>C 3345G>A 3346T>G), S1029T (3351G>C), Y1039A (3380T>G 3381A>C), N1042C (3389A>T 3390A>G), A1043V (3393C>T 3394A>T), E1047K (3404G>A 3406A>G), K1050Q (3413A>C), K1051S (3417A>G 3418G>T), V1052A (3420T>C 3421A>T), K1053N (3424A>T), T1055M (3429C>T 3430A>G), V1057I (3434G>A), V1063I (3452G>A 3454T>A), Y1064H (3455T>C), N1081G (3506A>G 3507A>G), V1085K (3518G>A 3519T>A 3520T>G), A1092K (3539G>A 3540C>A 3541T>G), T1093L (3542A>C 3543C>T 3544T>A), K1098T (3558A>C), V1104L (3575G>T 3577T>G), H1113K (3602C>A 3604C>G), V1122L (3629G>C 3631T>A), K1124A (3635A>G 3636A>C), S1133A (3662A>G 3663G>C 3664T>A), Q1140S (3683C>T 3684A>C 3685G>A), H1141Q (3688C>G), E1142D (3691A>C), V1143I (3692G>A 3694T>C), D1157K (3734G>A 3736C>A), I1159L (3740A>C 3742A>T), H1160Q (3745T>G), R1163Q (3752A>C 3753G>A), D1167Q (3764G>C 3766T>G), N1172Q (3779A>C 3781T>G), L1175I (3788T>A 3790A>T), F1178N (3797T>A 3798T>A), N1181A (3806A>G 3807A>C), D1184E (3817C>G), K1185Q (3818A>C 3820A>G), L1186V (3821C>G), S1188M (3827T>A 3828C>T 3829A>G), S1189D (3830A>G 3831G>A 3832C>T), F1190Y (3834T>A), E1192D (3841A>T), M1193N (3843T>A 3844G>C), K1194L (3845A>C 3846A>T), S1195K (3849G>A 3850T>G), E1196P (3851G>C 3852A>C 3853A>T), K1197R (3855A>G 3856G>A), Q1198V (3857C>G 3858A>T 3859A>G), V1199E (3861T>A 3862T>A), E1200A (3864A>C), Q1201P (3867A>C 3868A>T), I1203Q (3872A>C 3873T>A 3874C>A), A1204E (3876C>A 3877T>G), I1206P (3881A>C 3882T>C 3883T>A), K1208N (3889A>C), E1209T (3890G>A 3891A>C 3892G>A), V1211D (3897T>A), K1212_P1213del (3899_3904delAAGCCA), F1214S (3906T>C 3907T>C), I1215K (3909T>A), S1218E (3917A>G 3918G>A 3919T>G), P1220del (3923_3925delCCT), E1223V (3933A>T), R1225K (3939G>A 3940A>G), K1226P (3941A>C 3942A>C 3943A>T), Q1227V (3944C>G 3945A>T 3946A>C), D1229V (3951A>T 3952T>G), K1230_K1231insP (3955_3956insCCA), V1236I (3971G>A), E1237D (3976A>T), E1251N (4016G>A 4018A>T), N1252K (4021C>G), Y1256F (4032A>T), I1257A (4034A>G 4035T>C), N1262K (4051T>G), H1264Y (4055C>T 4057T>C), P1265H (4059C>A 4060A>T), A1268Q (4067G>C 4068C>A 4069C>G), T1269N (4071C>A 4072T>C), L1270M (4073C>A 4075T>G), V1271L (4076G>C), S1272R (4081T>A), D1273G (4083A>G 4084C>T), I1274E (4085A>G 4086T>A 4087T>A), I1276M (4093C>G), T1277S (4094A>T), K1280E (4103A>G), I1286M (4123A>G), V1291I (4136G>A 4138T>C), Q1292T (4139C>A 4140A>C 4141A>T), E1293S (4142G>A 4143A>G 4144G>T), V1295D (4149T>A), L1296I (4151T>A 4153A>C), A1298C (4157G>T 4158C>G), T1303S (4172A>T 4174T>C), A1314S (4205G>T 4207G>A), K1315R (4209A>G), R1318K (4218G>A 4219A>G), T1322V (4229A>G 4230C>T 4231A>T), N1324E (4235A>G 4237T>G), L1334C (4266T>G 4267A>T), N1335A (4268A>G 4269A>C), V1339L (4280G>C 4282A>T), V1345A (4299T>C 4300G>T), I1355V (4328A>G 4330T>A), I1359E (4340A>G 4341T>A 4342T>A), I1360A (4343A>G 4344T>C 4345C>A), S1361P (4346T>C), E1363A (4353A>C 4354G>T), Q1365E (4358C>G), V1391I (4436G>A 4438C>A), V1393M (4442G>A), E1394D (4447A>T), T1395V (4448A>G 4449C>T), K1396R (4452A>G), V1399M (4460G>A 4462T>G), S1400A (4463T>G), V1415I (4508G>A 4510G>C), A1420V (4524C>T 4525T>C), Y1423F (4533A>T), T1429E (4550A>G 4551C>A 4552A>G), T1430P (4553A>C), L1434I (4565C>A), N1436T (4572A>C 4573C>G), T1437K (4575C>A 4576A>G), D1440S (4583G>T 4584A>C), T1444P (4595A>C 4597T>G), L1450I (4613C>A), L1457F (4636A>T), Y1465C (4659A>G), V1471A (4677T>C 4678G>T), T1474V (4685A>G 4686C>T), A1485T (4718G>A 4720G>A), P1497S (4754C>T), I1502V (4769A>G 4771T>A), I1505V (4778A>G 4780C>T), K1512R (4800A>G), S1520R (4823T>C 4824C>G), Q1522E (4829C>G 4831A>G), I1525V (4838A>G 4840A>T), S1534I (4866G>T), Y1537H (4874T>C), T1538_S1539insL (4879_4880insCTG), S1539E (4880A>G 4881G>A 4882T>G), N1540S (4884A>G 4885T>C), T1542V (4889A>G 4890C>T), T1543E (4892A>G 4893C>A 4894A>G), I1551L (4916A>C 4918C>T), T1552S (4919A>T 4921C>A), F1553L (4922T>C), N1555K (4930T>A), T1558S (4938C>G 4939A>T), R1566K (4962G>A), I1577T (4995T>C), V1583L (5012G>C), N1611V (5096A>G 5097A>T 5098T>A), S1612N (5099T>A 5100C>A 5101A>T), Y1619F (5121A>T), N1623S (5133A>G), V1629S (5150G>A 5151T>G), T1638L (5177A>C 5178C>T), P1640E (5183C>G 5184C>A 5185T>G), Y1658F (5238A>T 5239C>T), N1662G (5249A>G 5250A>G), A1677S (5294G>T 5296C>T), T1678S (5298C>G), A1679V (5301C>T 5302A>T), T1682A (5309A>G), I1686L (5321A>C 5323A>T), L1688V (5327T>G 5329G>C), P1692A (5339C>G), D1697E (5356T>G), E1706D (5383A>T), C1718S (5417T>A), S1733T (5463G>C 5464T>C), Y1734H (5465T>C 5467C>T), F1736L (5471T>C 5473T>A), D1742E (5491T>A), C1744A (5495T>G 5496G>C 5497C>A), T1754H (5525A>C 5526C>A), Q1758K (5537C>A 5539G>A), Q1759T (5540C>A 5541A>C 5542G>T), K1763T (5553A>C), E1777D (5596A>T), Q1778N (5597C>A 5599A>T), F1779L (5600T>C), K1781T (5607A>C), Q1784S (5615C>T 5616A>C 5617G>C), T1788V (5627A>G 5628C>T), K1791R (5636A>C 5637A>G 5638A>T), Q1792D (5639C>G 5641A>T), K1795Q (5648A>C), P1803S (5672C>T), Q1813E (5702C>G), E1815K (5708G>A), K1817Q (5714A>C), H1818Q (5719T>A), T1822L (5729A>T 5730C>T 5731T>A), S1825N (5739G>A), K1837T (5775A>C 5776A>T), S1841A (5786T>G), C1847R (5804T>C 5806C>T), L1852H (5819T>C 5820T>A 5821A>C), S1856M (5831T>A 5832C>T 5833C>G), I1863V (5852A>G 5854T>G), N1871T (5877A>C 5878C>A), T1881S (5906A>T 5908T>G), V1888T (5927G>A 5928T>C), C1889Y (5931G>A 5932T>C), D1893E (5944C>A), N1898G (5957A>G 5958A>G 5959T>G), S1905A (5978T>G), F1907Y (5985T>A 5986C>T), N1917T (6015A>C 6016C>T), Y1920L (6024A>T 6025T>A), F1930L (6053T>C 6055T>C), V1931T (6056G>A 6057T>C), D1933S (6062G>T 6063A>C), I1935T (6069T>C 6070C>A), L1944M (6095T>A 6097A>G), Y1947F (6105A>T 6106T>C), K1948T (6108A>C 6109G>A), K1956S (6131A>T 6132A>C 6133A>T), K1973R (6183A>G), T1976S (6191A>T), P1977A (6194C>G 6196C>G), V1994I (6245G>A), N1996Q (6251A>C 6253T>G), N1999T (6261A>C 6262T>C), A2001T (6266G>A 6268C>A), Y2003F (6273A>T 6274T>C), I2010L (6293A>T), E2020D (6325A>T), D2026E (6343T>A), K2029A (6350A>G 6351A>C 6352G>A), S2030V (6353T>G 6354C>T), A2033T (6362G>A 6364G>A), D2043S (6392G>A 6393A>G), L2044Q (6396T>A), K2045Q (6398A>C), V2047T (6404G>A 6405T>C), D2060E (6445C>A), L2062I (6449C>A 6451T>A), N2065D (6458A>G 6460T>C), D2074N (6485G>A 6487C>T), I2075V (6488A>G 6490T>C), A2080S (6503G>T), N2081D (6506A>G), N2082E (6509A>G 6511T>A), S2083G (6512A>G), L2084V (6515T>G 6517A>T), I2086V (6521A>G 6523T>A), E2088Q (6527G>C), V2090L (6533G>T 6535T>A), T2093E (6542A>G 6543C>A 6544A>G), D2101E (6568C>A), S2103T (6572T>A 6574T>A), L2105I (6578C>A), R2115L (6608A>C 6609G>T), V2116A (6612T>C 6613A>C), L2122I (6629C>A), L2127I (6644T>A 6646A>T), V2130I (6653G>A), D2136S (6671G>A 6672A>G), T2137K (6675C>A 6676T>A), A2139L (6680G>T 6681C>T 6682T>G), N2140A (6683A>G 6684A>C), A2142V (6690C>T 6691T>C), N2147G (6704A>G 6705A>G 6706C>A), K2148Q (6707A>C), V2149A (6711T>C 6712T>A), V2150A (6714T>C 6715T>A), S2151I (6717G>T), T2154S (6725A>T 6727T>A), I2156C (6731A>T 6732T>G 6733A>C), V2157A (6735T>C), T2158K (6738C>A 6739A>G), C2160L (6744G>T 6745T>A), L2161A (6746T>G 6747T>C), N2162Q (6749A>C 6751C>A), C2165F (6759G>T), T2166N (6762C>A 6763T>C), F2172V (6779T>G 6781C>G), L2177F (6794C>T 6796A>C), R2184K (6816G>A), K2191R (6837A>G), M2194L (6845A>C 6847G>A), T2202S (6870C>G), G2207A (6885G>C), F2209L (6892T>A), E2212D (6901G>T), S2214G (6905T>G 6906C>G 6907A>C), F2215I (6908T>A), L2218V (6917T>G), N2222K (6931T>A), I2227F (6944A>T 6946A>C), N2228T (6948A>C 6949T>A), I2230A (6953A>G 6954T>C 6955A>T), I2231M (6958T>G), F2233L (6962T>C 6964T>A), V2238I (6977G>A), Y2245C (6999A>G 7000C>T), S2246V (7001T>G 7002C>T), L2250F (7015A>T), M2254L (7025A>T 7027G>A), L2257F (7036A>T), M2259A (7040A>G 7041T>C 7042G>T), T2264N (7056C>A), Y2266V (7061T>G 7062A>T 7063C>T), G2269L (7070G>T 7071G>T 7072C>G), T2274S (7085A>T), I2278T (7098T>C), A2279M (7100G>A 7101C>T 7102A>G), T2280D (7103A>G 7104C>A 7105C>T), Y2281F (7107A>T), T2283E

| | | | | | | | | | |
|---|---|---|---|---|---|---|---|---|---|
| | (7112A>G 7113C>A 7114T>A), I2286F (7121A>T 7123A>T), V2290I (7133G>A), T2300S (7163A>T 7165C>T), S2303A (7172T>G), I2309V (7190A>G 7192T>G), F2314Y (7206T>A 7207T>C), W2316L (7211T>C 7212G>T 7213G>A), A2320I (7223G>A 7224C>T), F2321L (7228T>A), V2324A (7236T>C 7237T>C), F2328V (7247T>G), I2332M (7261T>G), R2336K (7272G>A 7273G>A), V2340L (7283G>T), A2344S (7295G>T 7297T>A), L2349V (7310T>G), S2352G (7319A>G), V2356S (7331G>A 7332T>G 7333A>T), L2367F (7366A>T), N2370S (7374A>G), L2371I (7376C>A), I2377V (7394A>G), V2393I (7442G>A), V2400I (7463G>A 7465T>C), V2401M (7466G>A 7468A>G), N2405T (7479A>C 7480T>C), V2430M (7553G>A 7555T>G), R2431K (7557G>A 7558A>G), K2442R (7589A>C 7590A>G 7591A>T), L2447T (7604C>A 7605T>C 7606A>T), V2453L (7622G>C 7624T>C), A2460T (7643G>A), T2495A (7748A>G 7750A>T), S2500A (7763T>G 7765C>G), I2501L (7766A>C 7768C>T), S2517P (7814T>C 7816T>G), E2550D (7915A>C), S2553A (7922T>G), A2554S (7925G>T 7927A>T), A2584T (8015G>A 8017G>T), A2587S (8024G>T 8026A>C), N2596D (8051A>G 8053T>C), S2600A (8063T>G), N2603S (8073A>G 8074C>T), T2611A (8096A>G), E2617H (8114G>C 8116A>C), A2618S (8117G>A 8118C>G 8119T>C), N2623G (8132A>G 8133A>G), S2625A (8138T>G 8140C>T), N2628G (8147A>G 8148A>G), I2634V (8165A>G 8167T>G), F2641V (8186T>G), S2644T (8195T>A 8197A>C), E2647D (8206A>C), V2652I (8219G>A), Q2660H (8245A>C), I2663L (8252A>T), Y2673F (8283A>T 8284T>C), S2695N (8349G>A), I2709V (8390A>G), A2710S (8393G>T 8395T>A), F2718Y (8418T>A), L2738I (8477T>A), K2741R (8487A>G 8488G>A), V2754I (8525G>A), A2759S (8540G>T), N2767S (8565A>G), N2768T (8568A>C), W2769C (8572G>T), L2770F (8575G>T), Q2772L (8580A>T 8581G>T), L2773M (8582T>A 8584A>G), I2774L (8585A>C), V2776A (8592T>C 8593T>C), V2779L (8600G>T), F2780C (8604T>G), L2781V (8606C>G), F2782L (8609T>C), V2783A (8613T>C), A2785L (8618G>T 8619C>T 8620T>G), I2786V (8621A>G), F2787C (8625T>G 8626C>T), L2789I (8630T>A 8632A>C), I2790V (8633A>G 8635A>T), T2791M (8637C>T 8638A>G), V2795T (8648G>A 8649T>C 8650C>A), M2796L (8651A>T), K2798I (8658A>T 8659A>C), T2800D (8663A>G 8664C>A), D2801G (8667A>G 8668C>T), F2802Y (8670T>A 8671T>C), S2803T (8672T>A), S2804N (8676G>A), D2813Q (8702G>C 8704T>G), G2814D (8706G>A), A2821I (8726G>A 8727C>T 8728A>T), T2825D (8738A>G 8739C>A), D2833G (8763A>G), T2836A (8771A>G), T2846K (8802C>A 8803T>A), A2850S (8813G>A 8814C>G 8815T>C), L2853V (8822T>G 8824G>A), I2854V (8825A>G 8827T>A), V2857I (8834G>A), V2862I (8849G>A 8851G>T), V2865I (8858G>A 8860C>A), I2873V (8882A>G 8884A>G), T2876A (8891A>G), T2877I (8895C>T 8896T>C), T2906S (8982C>G), S2926M (9041T>A 9042C>T 9043T>G), V2938L (9077G>T 9079A>G), V2943I (9092G>A), A2944S (9095G>T), E2946S (9101G>A 9102A>G 9103A>T), S2947E (9104A>G 9105G>A 9106T>G), S2981A (9206T>G), A2994V (9246C>T 9247T>A), V2996I (9251G>A), V2998L (9257G>C), D3009E (9292T>G), Y3010H (9293T>C), S3013A (9302T>G), P3015S (9308C>T), V3024M (9335G>A 9337A>G), L3027I (9344C>A 9346T>A), T3028A (9347A>G), M3030I (9355G>C), I3035V (9368A>G 9370T>G), I3038V (9377A>G 9379T>G), I3043V (9392A>G 9394A>G), I3047V (9404A>G), V3053I (9422G>A 9424A>T), V3056L (9431G>T 9433A>G), L3060A (9443C>G 9444T>C), R3066K (9462G>A 9463G>A), A3070V (9474C>T), S3075N (9489G>A 9490T>C), F3080A (9503T>G 9504T>C), T3082A (9509A>G 9511T>A), V3091I (9536G>A), T3095V (9548A>G 9549C>T), V3097A (9555T>C), I3108F (9587A>T), L3116F (9611C>T 9613T>C), I3126L (9641A>C), M3129F (9650A>T 9652G>T), V3130A (9654T>C 9655T>C), T3133S (9662A>T 9664A>T), L3135I (9668T>A 9670A>T), I3142A (9689A>G 9690T>C 9691T>A), A3143I (9692G>A 9693C>T 9694T>C), I3145V (9698A>G 9700C>A), I3146F (9701A>T 9703T>C), T3150L (9713A>C 9714C>T 9715A>G), F3153C (9723T>G), Y3154H (9725T>C), S3158N (9738G>A 9739T>C), K3162R (9750A>G), R3163K (9753G>A), V3166M (9761G>A 9763C>G), S3171T (9776T>A 9778C>A), D3196E (9853T>G), V3197T (9854G>A 9855T>C 9856G>A), M3221L (9926A>T 9928G>A), S3246A (10001T>G), V3298T (10157G>A 10158T>C), S3309A (10190T>G 10192T>A), N3328S (10248A>G 10249T>C), V3349L (10310G>C 10312A>G), K3351R (10317A>G), A3357S (10334G>T 10336C>T), F3397H (10454T>C 10455T>A 10456C>T), N3443K (10594C>A), V3465L (10658G>T 10660T>A), S3530A (10853T>G 10855A>T), A3548T (10907G>A), L3549I (10910T>A 10912A>T), S3570G (10973A>G), A3571K (10976G>A 10977C>A 10978A>G), V3572F (10979G>T 10981G>C), R3574K (10986G>A), T3575I (10989C>T 10990A>T), I3576V (10991A>G 10993C>T), L3583M (11012T>A), I3587F (11024A>T 11026T>C), V3593I (11042G>A), L3606V (11081T>G 11083G>T), A3615T (11108G>A), M3616L (11111A>C 11113G>T), I3619M (11122T>G), M3621I (11128G>T), S3622A (11129T>G), F3624C (11136T>G), M3627L (11144A>C), F3628L (11147T>C), D3668E (11269T>A), M3669L (11270A>T), V3670A (11274T>C), F3677Y (11295T>A), K3678R (11298A>G), V3689L (11330G>T), G3704A (11376G>C), L3715I (11408T>A 11410G>T), I3737V (11474A>G 11476A>T), V3751I (11516G>A), G3757A (11535G>C), M3761V (11546A>G), C3766Y (11562G>A), I3768L (11567A>T 11569T>G), F3769L (11572C>A), F3789C (11631T>G), T3791C (11636A>T 11637C>G 11638T>C), N3833S (11763A>G), V3847I (11804G>A), K3929R (12051A>G), F3957Y (12135T>A), I4074V (12485A>G 12487A>C), N4078G (12497A>G 12498A>G 12499C>T), T4087N (12525C>A 12526A>C), S4115N (12609G>A 12610T>C), T4174N (12786C>A 12787A>T), T4175S (12788A>T 12790A>G), L4188H (12827T>C 12828T>A 12829A>C), A4276P (13091G>C), K4366R (13362A>G), M4390L (13433A>T), L4391M (13436C>A 13438T>G), Q4397S (13454C>T 13455A>C), S4398T (13457T>A), D4432E (13560T>A), D4454E (13626T>A), D4455G (13628A>G), I4458L (13636A>T 13638T>A), F4469M (13669T>A 13671C>G), L4482V (13708C>G), K4490V (13732A>G 13733A>T 13734A>C), I4498V (13756A>G), A4577S (13993G>T 13995T>A), N4590D (14032A>G), I4615V (14107A>G), T4617V (14113A>G 14114C>T 14115C>A), T4618A (14116A>G 14118G>A), S4621C (14125A>T 14127T>C), V4625I (14137G>A), T4644A (14194A>G), V4649M (14209G>A 14211T>G), T4651A (14215A>G), T4654A (14224A>G), Y4657L (14233T>C 14234A>T 14235C>T), K4673C (14281A>T 14282A>G 14283A>T), V4691I (14335G>A), N5003T (15272A>C 15273C>T), T5035N (15368C>A 15369A>C), S5039N (15380G>A), T5131H (15655A>C 15656C>A 15657A>T), D5132E (15660C>A), N5135D (15667A>G), F5158Y (15737T>A 15738C>T), T5161N (15746C>A 15747T>C), S5164A (15754T>G), S5176A (15790T>G), V5894I (17944G>A 17946A>T), V5939I (18079G>A 18081A>C), T5956I (18131C>T 18132T>A), E6024D (18336A>T), P6053E (18421C>G 18422C>A 18423T>A), D6057E (18435T>A), S6059T (18439T>A), S6062N (18449G>A), N6101G (18565A>G 18566A>G 18567T>A), R6137K (18674G>A), A6145S (18697G>T 18699T>A), H6153N (18721C>A), I6156V (18730A>G 18732T>G), L6184Q (18815T>A 18816G>A), Y6185H (18817T>C), T6218S (18916A>T), I6219V (18919A>G), K6229R (18950A>G), I6230V (18952A>G), A6232S (18958G>T 18960G>T), A6244S (18994G>T), D6270E (19074T>A), S6299H (19159T>C 19160C>A 19161T>C), S6321A (19225T>G 19227C>A), V6362T (19348G>A 19349T>C), L6418Q (19516T>C 19517T>A), V6435I (19567G>A), F6459Y (19640T>A), Q6470H (19674A>C), Q6471A (19675C>G 19676A>C 19677G>C), V6474A (19685T>C), T6482A (19708A>G), V6490I (19732G>A), L6494I (19744T>A 19746G>C), V6521I (19825G>A 19827G>T), D6543E (19893T>A), I6548V (19906A>G), S6555T (19927T>A 19929T>A), T6566S (19961C>G 19962G>T), I6567A (19963A>G 19964T>C), A6569S (19969G>T 19971A>T), P6570S (19972C>T), F6574L (19986T>G), D6580E (20004T>A), Q6604T (20074C>A 20075A>C), V6607K (20083G>A 20084T>A 20085A>G), K6610A (20092A>G 20093A>C), L6614V (20104C>G 20106T>C), A6623S (20131G>T 20133C>A), Y6631F (20156A>T), V6637I (20173G>A), V6638I (20176G>A 20178C>T), N6651D (20215A>G 20217T>C), Q6653E (20221C>G 20223A>G), E6654D (20226A>T), I6663T (20252T>C), E6675Q (20287G>C 20289A>G), S6695G (20347A>G 20349T>A), L6703M (20371C>A), F6710S (20393T>C 20394T>A), K6711Q (20395A>C 20397G>A), E6712D (20400A>T), F6715L (20407T>C), E6716K (20410G>A), V6766I (20560G>A), T6777A (20593A>G 20595A>T), S6799A (20659T>G 20661T>A), D6830E (20754T>A), S6831N (20756G>A), T6833V (20761A>G 20762C>T 20763A>T), L6834I (20764T>A), K6933R (21062A>G), N6936H (21070A>C), I6951L (21115A>C 21117T>G), Q6956K (21130C>A 21132A>G), V6965I (21157G>A 21159G>A), I6967V (21163A>G), A6986S (21220G>T), C7007A (21283T>G 21284G>C), R7014K (21304C>A 21305G>A 21306C>G), V7021T (21325G>A 21326T>C), G7063N (21451G>A 21452G>A), L7070Y (21473T>A 21474A>T), S7074E (21484A>G 21485G>A 21486T>A), I7088V (21526A>G), V7092I (21538G>A) | | | | | | | | |
| leader protein (YP_009725297.1) | 1 | 180 | 100% | 1059.0 | 86.0% | 180 (100%) | 152 (84.4%) | 0/0/0/0 | 0 |
| | P6L (282C>T), F8V (287T>G), V38A (378T>C), Q44E (395C>G), D48N (407G>A), V56L (431G>C 433T>G), R77L (494C>T 495G>T), T78S (498C>G 499T>C), A79T (500G>A 502A>C), P80N (503C>A 504C>A), V84K (515G>A 516T>A 517T>G), M85V (518A>G 520G>C), L92M (539C>A 541C>G), E93D (544A>C), E102I (569G>A 570A>T 571G>A), I114T (606T>C 607A>C), V116I (611G>A 613G>T), K120N (625G>T), A138I (677G>A 678C>T), F143Y (693T>A), Y154I (725T>A 726A>T), F157Y (735T>A), Q158E (737C>G), E159Q (740G>C), S166G (761A>G), V169A (771T>C 772T>A), T170L (773A>C 774C>T), M174T (786T>C 787G>T) | | | | | | | | |
| nsp2 (YP_009725298.1) | 1 | 638 | 100% | 3197.0 | 72.6% | 638 (100%) | 436 (68.3%) | 0/0/0/0 | 0 |
| | Y2V (809T>G 810A>T), E19D (862G>T), L24F (875C>T), A31S (896G>T 898T>A), S32M (899T>A 900C>T 901A>G), F41Y (927T>A 928T>C), D43E (934C>G), T44S (935A>T 937T>G), E53D (964A>C), Y61F (987A>T), E66D (1003A>T), L71H (1016T>C 1017T>A 1018G>C), L79S (1040T>A 1041T>G 1042G>T), N87K (1066T>A), N92K (1081T>G), I100K (1104T>A), I101V (1106A>G), T103V (1112A>G 1113C>T 1114T>C), L113T (1142C>A 1143T>C), D114E (1147T>G), N130Q (1193A>C 1195T>G), Q134N (1205C>A 1207A>T), C136H (1211T>C 1212G>A), D144N (1235G>A), G147D (1245G>A), T149V (1250A>G 1251C>T), G154C (1265G>T), V157L (1274G>C 1276T>G), F163H (1292T>C 1293T>A), T170V (1313A>G 1314C>T), K171I (1317A>T 1318A>T), A174P (1325G>C 1327C>T), Q182T (1349C>A 1350A>C 1351A>T), I188M (1369T>G), Y189P (1370T>C 1371A>C 1372T>A), H194Q (1387C>A), N195D (1388A>G 1390T>C), S196P (1391T>C), V198I (1397G>A 1399A>T), L204V (1415C>G), E206D (1423A>T), E210H (1433G>C 1435A>C), G212N (1439G>A 1440G>A), L213I (1442T>A 1444G>T), K214E (1445A>G), I216R (1451A>C 1452T>G 1453T>A), I224R (1476T>G 1477T>A), A225C (1478G>T 1479C>G 1480C>T), S232A (1499T>G 1501T>C), H237Y (1514C>T), C240R (1523T>C), N250D (1553A>G 1555C>T), C253S (1563G>C 1564T>A), N254G (1565A>G 1566A>G), V258I (1577G>A), V259T (1580G>A 1581T>C), E261D (1588A>C), G262N (1589G>A 1590G>A), S263V (1592T>G 1593C>T 1594C>G), G265T (1598G>A 1599G>C 1600T>C), D268E (1609C>G), N269D (1610A>G 1612C>T), Q275S (1628C>A 1629A>G 1630A>T), K276R (1631A>C 1632A>G 1633A>T), K278R (1637A>C 1638A>G 1639A>T), K288H (1667A>C 1669A>T), I293V (1682A>G 1684C>T), V308I (1727G>A 1729G>T), E309D (1732A>C), V311I (1736G>A 1738G>A), G313S (1742G>A), A318S (1757G>T 1759A>T), Q321T (1766C>A 1767A>C 1768A>C), F329Y (1791T>A), A336P (1811G>C 1813T>C), K337V (1814A>G 1815A>T), E345Q (1838G>C), K347R (1845A>G), I349V (1850A>G 1852A>T), S351T (1857G>C 1858T>A), Y354C (1866A>G), A355G (1869C>G 1870A>T), A357P (1874G>C 1876A>C), E359Q (1880G>C), R362G (1889C>G), V364I (1895G>A 1897A>C), S369A (1910T>G 1912C>G), E373D (1924A>T), T374A (1925A>G 1927T>A), Q376N (1931C>A 1933A>C), N377H (1934A>C 1936T>C), V379I (1940G>A 1942G>T), R380P (1944G>C), V381D (1947T>A), K384R (1956A>G 1957G>A), I387V (1964A>G 1966A>C), Q395E (1988C>G 1990G>A), Y396Q (1991T>C 1993T>G), I401V (2006A>G 2008T>C), M405V (2018A>G 2020G>T), F406Y (2022T>A 2023C>T), A411L (2036G>C 2037C>T 2038T>C), N414S (2046A>G), L415V (2048C>G 2050A>C), V416I (2051G>A), V417I (2054G>A 2056A>T), I421V (2066A>G 2068T>A), V425L (2078G>C), L428Q (2087T>C 2088T>A), T434S (2105A>T), I436L (2111A>C 2113C>T), F437L (2116T>G), V440T (2123G>A 2124T>C), Y441V (2126T>G 2127A>T), K445R (2139A>G 2140A>G), V447I (2144G>A), L448F (2147C>T), D449E (2152T>A), L451I (2156C>A), E453A (2163A>C), F455L (2168T>C), K456S (2172A>G 2173G>T), E457A (2175A>C), R463K (2193G>A 2194A>G), G465A (2199G>C), V469L (2210G>C 2212T>C), I472L (2219A>C), S473I (2222T>A 2223C>T 2224A>T), C475G (2228T>G), A476V (2232C>T), C477F (2235G>T), E478D (2239A>C), G481K (2246G>A 2247G>A 2248T>G), V485Q (2258G>C 2259T>A 2260C>G), T486V (2261A>G 2262C>T 2263C>T), C487A (2264T>G 2265G>C), A488S (2267G>T), K489D (2270A>G 2272G>T), E490N (2273G>A 2275A>C), E493D (2284G>T), S494C (2285A>T), Q496K (2291C>A 2293G>A), T497C (2294A>T 2295C>G 2296A>C), F499I (2300T>A), K500D (2303A>G 2305G>T), L501V (2306C>G), F505A (2318T>G 2319T>C 2320T>A), A507E (2325C>A 2326T>A), L508M (2327T>A), A510I (2333G>A 2334C>T), S512Q (2339T>C 2340C>A 2341T>A), I513V (2342A>G), I514T (2346T>C), G516A (2352G>C), K521R (2366A>C 2367A>G), A522S (2369G>T 2371C>A), T528V (2387A>G 2388C>T 2389A>C), V530I (2393G>A), T531A (2396A>G 2398G>T), H532Q (2401C>A), K539Q (2420A>C), V541I (2426G>A 2428T>A), K542R (2429A>C 2430A>G 2431A>T), S543G (2432T>G 2433C>G), R544K (2436G>A 2437A>G), E546Q (2441G>C 2443A>G), T547L (2444A>C 2445C>T 2446T>G), G548Q (2447G>C 2448G>A 2449C>A), I559V (2480A>G 2482T>A), I560T (2484T>C), E565D (2500A>T), T566S (2501A>T), L567H (2505T>A), P568D (2507C>G 2508C>A), E570V (2514A>T), V571L (2516G>C 2518G>T), L572T (2519T>A 2520T>C 2521A>C), T573S (2522A>T 2524A>T), T580N (2544C>A 2545T>C), D582E (2551T>A), Q584E (2555C>G), P585A (2558C>G), Q588T (2567C>A 2568A>C 2569A>G), T590V (2573A>G 2574C>T), S591D (2576A>G 2577G>A), E592S (2579G>A 2580A>G 2581A>C), A593F (2582G>T 2583C>T 2584T>C), V594T (2585G>A 2586T>C 2587T>A), E595N (2588G>A 2590A>T), A596G (2592C>G 2593T>A), P597A (2594C>G 2596A>T), L598I (2597T>A 2599G>C), I605V (2618A>G 2620T>A), T616K (2652C>A), K618Q (2657A>C 2659G>A), A623S (2672G>T 2674A>T), N625G (2678A>G 2679A>G), M626L (2681A>T 2683G>A), M627L (2684A>C), V628A (2688T>C 2689A>T), T632V (2699A>G 2700C>T), T634R (2705A>C 2706C>G 2707A>C) | | | | | | | | |

| | Begin | End | Coverage | Score | Concordance | Matches | Identities | I/D/M/F* | Stop Codons |
|---|---|---|---|---|---|---|---|---|---|
| nsp3 (YP_009725299.1) | 1 | 1945 | 100% | 10414.0 | 80.5% | 1918 (98.4%) | 1485 (76.2%) | 4/27/0/0 | 0 |
| | T3I (2727C>T 2728A>T), K4_V5insG (2731_2732insGGT), D9E (2746T>A), I13W (2756A>T 2757T>G 2758A>G), S20N (2778G>A), N22R (2784A>G 2785T>A), I31V (2810A>G), A41V (2841C>T), L46S (2855C>T 2856T>C), N51T (2871A>C), D59E (2896T>G), I62V (2903A>G 2905A>G), E70D (2929A>T), P74N (2939C>A 2940C>A 2941A>C), L75M (2942C>A), M84V (2969A>G 2971G>A), Y87F (2979A>T), E92D (2995G>T), S93A (2996T>G), F96E (3005T>G 3006T>A 3007T>A), K97N (3010A>C), L98F (3013G>T), A99S (3014G>T 3016T>A), H101R (3021A>G), D112E (3055T>A), E115D (3064A>C), G116D (3066G>A), G116_D117insA (3067_3068insGCA), D117E (3070T>G), F123I (3086T>A), E124D (3091G>T), P125E (3092C>G 3093C>A), S126T (3095T>A 3097A>C), T127C (3098A>T 3099C>G), Q128E (3101C>G), Y129H (3104T>C), K140L (3137A>C 3138A>T 3139A>C), T147S (3158A>T 3160T>A), S148A (3161T>G), A149E (3165C>A 3166T>A), A150T (3167G>A 3169T>A), L151V (3170C>G), Q152R (3174A>G), P153V (3176C>G 3177C>T), Q157E (3188C>G), D165_T181del (3212_3262delGATAGTCAACAAACTGTTGGTCAACAAGACGGCAGTGAGGACAATCAGACA), I184E (3269A>G 3270T>A 3271T>G), T186S (3275A>T), I187_V188del (3278_3283delATTGTT), V190I (3287G>A), Q191E (3290C>G 3292A>G), Q193E (3296C>G), L194P (3299T>C 3300T>C), E195_M196del (3302_3307delGAGATG), L198P (3312T>C), V201E (3321T>A 3322T>A), V202E (3324T>A 3325T>A), Q203_I205del (3327_3335delAGACTATTG), E206P (3327_3335delAGACTATTG 3336A>C), S209Q (3344A>C 3345G>A 3346T>G), S211T (3351G>C), Y221A (3380T>G 3381A>C), N224C (3389A>T 3390A>G), A225V (3393C>T 3394A>T), E229K (3404G>A 3406A>G), K232Q (3413A>C), K233S (3417A>G 3418G>T), V234A (3420T>C 3421A>T), K235N (3424A>T), T237M (3429C>T 3430A>G), V239I (3434G>A), V245I (3452G>A 3454T>A), Y246H (3455T>C), N263G (3506A>G 3507A>G), V267K (3518G>A 3519T>A 3520T>G), A274K (3539G>A 3540C>A 3541T>G), T275L (3542A>C 3543C>T 3544T>A), K280T (3558A>C), V286L (3575G>T 3577T>G), H295K (3602C>A 3604C>G), V304L (3629G>C 3631T>A), K306A (3635A>G 3636A>C), S315A (3662A>G 3663G>C 3664T>A), Q322S (3683C>T 3684A>C 3685G>A), H323Q (3688C>G), E324D (3691A>C), V325I (3692G>A 3694T>C), D339K (3734G>A 3736C>A), I341L (3740A>C 3742A>T), H342Q (3745T>G), R345Q (3752A>C 3753G>A), D349Q (3764G>C 3766T>G), N354Q (3779A>C 3781T>G), L357I (3788T>A 3790A>T), F360N (3797T>A 3798T>A), N363A (3806A>G 3807A>C), D366E (3817C>G), K367Q (3818A>C 3820A>G), L368V (3821C>G), S370M (3827T>A 3828C>T 3829A>G), S371D (3830A>G 3831G>A 3832C>T), F372Y (3834T>A), E374D (3841A>T), M375N (3843T>A 3844G>C), K376L (3845A>C 3846A>T), S377K (3849G>A 3850T>G), E378P (3851G>C 3852A>C 3853A>T), K379R (3855A>G 3856G>A), Q380V (3857C>G 3858A>T 3859A>G), V381E (3861T>A 3862T>A), E382A (3864A>C), Q383P (3867A>C 3868A>T), I385Q (3872A>C 3873T>A 3874C>A), A386E (3876C>A 3877T>G), I388P (3881A>C 3882T>C 3883T>A), K390N (3889A>C), E391T (3890G>A 3891A>C 3892G>A), V393D (3897T>A), K394_P395del (3899_3904delAAGCCA), F396S (3906T>C 3907T>C), I397K (3909T>A), S400E (3917A>G 3918G>A 3919T>G), P402del (3923_3925delCCT), E405V (3933A>T), R407K (3939G>A 3940A>G), K408P (3941A>C 3942A>C 3943A>T), Q409V (3944C>G 3945A>T 3946A>C), D411V (3951A>T 3952T>G), K412_K413insP (3955_3956insCCA), V418I (3971G>A), E419D (3976A>T), E433N (4016G>A 4018A>T), N434K (4021C>G), Y438F (4032A>T), I439A (4034A>G 4035T>C), N444K (4051T>G), H446Y (4055C>T 4057T>C), P447H (4059C>A 4060A>T), A450Q (4067G>C 4068C>A 4069C>G), T451N (4071C>A 4072T>C), L452M (4073C>A 4075T>G), V453L (4076G>C), S454R (4081T>A), D455G (4083A>G 4084C>T), I456E (4085A>G 4086T>A 4087T>A), I458M (4093C>G), T459S (4094A>T), K462E (4103A>G), I468M (4123A>G), V473I (4136G>A 4138T>C), Q474T (4139C>A 4140A>C 4141A>T), E475S (4142G>A 4143A>G 4144G>T), V477D (4149T>A), L478I (4151T>A 4153A>C), A480C (4157G>T 4158C>G), T485S (4172A>T 4174T>C), A496S (4205G>T 4207G>A), K497R (4209A>G), R500K (4218G>A 4219A>G), T504V (4229A>G 4230C>T 4231A>T), N506E (4235A>G 4237T>G), L516C (4266T>G 4267A>T), N517A (4268A>G 4269A>C), V521L (4280G>C 4282A>T), V527A (4299T>C 4300G>T), I537V (4328A>G 4330T>A), I541E (4340A>G 4341T>A 4342T>A), I542A (4343A>G 4344T>C 4345C>A), S543P (4346T>C), E545A (4353A>C 4354G>T), Q547E (4358C>G), V573I (4436G>A 4438C>A), V575M (4442G>A), E576D (4447A>T), T577V (4448A>G 4449C>T), K578R (4452A>G), V581M (4460G>A 4462T>G), S582A (4463T>G), V597I (4508G>A 4510G>C), A602V (4524C>T 4525T>C), Y605F (4533A>T), T611E (4550A>G 4551C>A 4552A>G), T612P (4553A>C), L616I (4565C>A), N618T (4572A>C 4573C>G), T619K (4575C>A 4576A>G), D622S (4583G>T 4584A>C), T626P (4595A>C 4597T>G), L632I (4613C>A), L639F (4636A>T), Y647C (4659A>G), V653A (4677T>C 4678G>T), T656V (4685A>G 4686C>T), A667T (4718G>A 4720G>A), P679S (4754C>T), I684V (4769A>G 4771T>A), I687V (4778A>G 4780C>T), K694R (4800A>G), S702R (4823T>C 4824C>G), Q704E (4829C>G 4831A>G), I707V (4838A>G 4840A>T), S716I (4866G>T), Y719H (4874T>C), T720_S721insL (4879_4880insCTG), S721E (4880A>G 4881G>A 4882T>G), N722S (4884A>G 4885T>C), T724V (4889A>G 4890C>T), T725E (4892A>G 4893C>A 4894A>G), I733L (4916A>C 4918C>T), T734S (4919A>T 4921C>A), F735L (4922T>C), N737K (4930T>A), T740S (4938C>G 4939A>T), R748K (4962G>A), I759T (4995T>C), V765L (5012G>C), N793V (5096A>G 5097A>T 5098T>A), S794N (5099T>A 5100C>A 5101A>T), Y801F (5121A>T), N805S (5133A>G), V811S (5150G>A 5151T>G), T820L (5177A>C 5178C>T), P822E (5183C>G 5184C>A 5185T>G), Y840F (5238A>T 5239C>T), N844G (5249A>G 5250A>G), A859S (5294G>T 5296C>T), T860S (5298C>G), A861V (5301C>T 5302A>T), T864A (5309A>G), I868L (5321A>C 5323A>T), L870V (5327T>G 5329G>C), P874A (5339C>G), D879E (5356T>G), E888D (5383A>T), C900S (5417T>A), S915T (5463G>C 5464T>C), Y916H (5465T>C 5467C>T), F918L (5471T>C 5473T>A), D924E (5491T>A), C926A (5495T>G 5496G>C 5497C>A), T936H (5525A>C 5526C>A), Q940K (5537C>A 5539G>A), Q941T (5540C>A 5541A>C 5542G>T), K945T (5553A>C), E959D (5596A>T), Q960N (5597C>A 5599A>T), F961L (5600T>C), K963T (5607A>C), Q966S (5615C>T 5616A>C 5617G>C), T970V (5627A>G 5628C>T), K973R (5636A>C 5637A>G 5638A>T), Q974D (5639C>G 5641A>T), K977Q (5648A>C), P985S (5672C>T), Q995E (5702C>G), E997K (5708G>A), K999Q (5714A>C), H1000Q (5719T>A), T1004L (5729A>T 5730C>T 5731T>A), S1007N (5739G>A), K1019T (5775A>C 5776A>T), S1023A (5786T>G), C1029R (5804T>C 5806C>T), L1034H (5819T>C 5820T>A 5821A>C), S1038M (5831T>A 5832C>T 5833C>G), I1045V (5852A>G 5854T>G), N1053T (5877A>C 5878C>A), T1063S (5906A>T 5908T>G), V1070T (5927G>A 5928T>C), C1071Y (5931G>A 5932T>C), D1075E (5944C>A), N1080G (5957A>G 5958A>G 5959T>G), S1087A (5978T>G), F1089Y (5985T>A 5986C>T), N1099T (6015A>C 6016C>T), Y1102L (6024A>T 6025T>A), F1112L (6053T>C 6055T>C), V1113T (6056G>A 6057T>C), D1115S (6062G>T 6063A>C), I1117T (6069T>C 6070C>A), L1126M (6095T>A 6097A>G), Y1129F (6105A>T 6106T>C), K1130T (6108A>C 6109G>A), K1138S (6131A>T 6132A>C 6133A>T), K1155R (6183A>G), T1158S (6191A>T), P1159A (6194C>G 6196C>G), V1176I (6245G>A), N1178Q (6251A>C 6253T>G), N1181T (6261A>C 6262T>C), A1183T (6266G>A 6268C>A), Y1185F (6273A>T 6274T>C), I1192L (6293A>T), E1202D (6325A>T), D1208E (6343T>A), K1211A (6350A>G 6351A>C 6352G>A), S1212V (6353T>G 6354C>T), A1215T (6362G>A 6364G>A), D1225S (6392G>A 6393A>G), L1226Q (6396T>A), K1227Q (6398A>C), V1229T (6404G>A 6405T>C), D1242E (6445C>A), L1244I (6449C>A 6451T>A), N1247D (6458A>G 6460T>C), D1256N (6485G>A 6487C>T), I1257V (6488A>G 6490T>C), A1262S (6503G>T), N1263D (6506A>G), N1264E (6509A>G 6511T>A), S1265G (6512A>G), L1266V (6515T>G 6517A>T), I1268V (6521A>G 6523T>A), E1270Q (6527G>C), V1272L (6533G>T 6535T>A), T1275E (6542A>G 6543C>A 6544A>G), D1283E (6568C>A), S1285T (6572T>A 6574T>A), L1287I (6578C>A), R1297L (6608A>C 6609G>T), V1298A (6612T>C 6613A>C), L1304I (6629C>A), L1309I (6644T>A 6646A>T), V1312I (6653G>A), D1318S (6671G>A 6672A>G), T1319K (6675C>A 6676T>A), A1321L (6680G>T 6681C>T 6682T>G), N1322A (6683A>G 6684A>C), A1324V (6690C>T 6691T>C), N1329G (6704A>G 6705A>G 6706C>A), K1330Q (6707A>C), V1331A (6711T>C 6712T>A), V1332A (6714T>C 6715T>A), S1333I (6717G>T), T1336S (6725A>T 6727T>A), I1338C (6731A>T 6732T>G 6733A>C), V1339A (6735T>C), T1340K (6738C>A 6739A>G), C1342L (6744G>T 6745T>A), L1343A (6746T>G 6747T>C), N1344Q (6749A>C 6751C>A), C1347F (6759G>T), T1348N (6762C>A 6763T>C), F1354V (6779T>G 6781C>G), L1359F (6794C>T 6796A>C), R1366K (6816G>A), K1373R (6837A>G), M1376L (6845A>C 6847G>A), T1384S (6870C>G), G1389A (6885G>C), F1391L (6892T>A), E1394D (6901G>T), S1396G (6905T>G 6906C>G 6907A>C), F1397I (6908T>A), L1400V (6917T>G), N1404K (6931T>A), I1409F (6944A>T 6946A>C), N1410T (6948A>C 6949T>A), I1412A (6953A>G 6954T>C 6955A>T), I1413M (6958T>G), F1415L (6962T>C 6964T>A), V1420I (6977G>A), Y1427C (6999A>G 7000C>T), S1428V (7001T>G 7002C>T), L1432F (7015A>T), M1436L (7025A>T 7027G>A), L1439F (7036A>T), M1441A (7040A>G 7041T>C 7042G>T), T1446N (7056C>A), Y1448V (7061T>G 7062A>T 7063C>T), G1451L (7070G>T 7071G>T 7072C>G), T1456S (7085A>T), I1460T (7098T>C), A1461M (7100G>A 7101C>T 7102A>G), T1462D (7103A>G 7104C>A 7105C>T), Y1463F (7107A>T), T1465E (7112A>G 7113C>A 7114T>A), I1468F (7121A>T 7123A>T), V1472I (7133G>A), T1482S (7163A>T 7165C>T), S1485A (7172T>G), I1491V (7190A>G 7192T>G), F1496Y (7206T>A 7207T>C), W1498L (7211T>C 7212G>T 7213G>A), A1502I (7223G>A 7224C>T), F1503L (7228T>A), V1506A (7236T>C 7237T>C), F1510V (7247T>G), I1514M (7261T>G), R1518K (7272G>A 7273G>A), V1522L (7283G>T), A1526S (7295G>T 7297T>A), L1531V (7310T>G), S1534G (7319A>G), V1538S (7331G>A 7332T>G 7333A>T), L1549F (7366A>T), N1552S (7374A>G), L1553I (7376C>A), I1559V (7394A>G), V1575I (7442G>A), V1582I (7463G>A 7465T>C), V1583M (7466G>A 7468A>G), N1587T (7479A>C 7480T>C), V1612M (7553G>A 7555T>G), R1613K (7557G>A 7558A>G), K1624R (7589A>C 7590A>G 7591A>T), L1629T (7604C>A 7605T>C 7606A>T), V1635L (7622G>C 7624T>C), A1642T (7643G>A), T1677A (7748A>G 7750A>T), S1682A (7763T>G 7765C>G), I1683L (7766A>C 7768C>T), S1699P (7814T>C 7816T>G), E1732D (7915A>C), S1735A (7922T>G), A1736S (7925G>T 7927A>T), A1766T (8015G>A 8017G>T), A1769S (8024G>T 8026A>C), N1778D (8051A>G 8053T>C), S1782A (8063T>G), N1785S (8073A>G 8074C>T), T1793A (8096A>G), E1799H (8114G>C 8116A>C), A1800S (8117G>A 8118C>G 8119T>C), N1805G (8132A>G 8133A>G), S1807A (8138T>G 8140C>T), N1810G (8147A>G 8148A>G), I1816V (8165A>G 8167T>G), F1823V (8186T>G), S1826T (8195T>A 8197A>C), E1829D (8206A>C), V1834I (8219G>A), Q1842H (8245A>C), I1845L (8252A>T), Y1855F (8283A>T 8284T>C), S1877N (8349G>A), I1891V (8390A>G), A1892S (8393G>T 8395T>A), F1900Y (8418T>A), L1920I (8477T>A), K1923R (8487A>G 8488G>A), V1936I (8525G>A), A1941S (8540G>T) | | | | | | | | |
| nsp4 (YP_009725300.1) | 1 | 500 | 100% | 3038.0 | 86.1% | 500 (100%) | 400 (80.0%) | 0/0/0/0 | 0 |
| | N4S (8565A>G), N5T (8568A>C), W6C (8572G>T), L7F (8575G>T), Q9L (8580A>T 8581G>T), L10M (8582T>A 8584A>G), I11L (8585A>C), V13A (8592T>C 8593T>C), V16L (8600G>T), F17C (8604T>G), L18V (8606C>G), F19L (8609T>C), V20A (8613T>C), A22L (8618G>T 8619C>T 8620T>G), I23V (8621A>G), F24C (8625T>G 8626C>T), L26I (8630T>A 8632A>C), I27V (8633A>G 8635A>T), T28M (8637C>T 8638A>G), V32T (8648G>A 8649T>C 8650C>A), M33L (8651A>T), K35I (8658A>T 8659A>C), T37D (8663A>G 8664C>A), D38G (8667A>G 8668C>T), F39Y (8670T>A 8671T>C), S40T (8672T>A), S41N (8676G>A), D50Q (8702G>C 8704T>G), G51D (8706G>A), A58I (8726G>A 8727C>T 8728A>T), T62D (8738A>G 8739C>A), D70G (8763A>G), T73A (8771A>G), T83K (8802C>A 8803T>A), A87S (8813G>A 8814C>G 8815T>C), L90V (8822T>G 8824G>A), I91V (8825A>G 8827T>A), V94I (8834G>A), V99I (8849G>A 8851G>T), V102I (8858G>A 8860C>A), I110V (8882A>G 8884A>G), T113A (8891A>G), T114I (8895C>T 8896T>C), T143S (8982C>G), S163M (9041T>A 9042C>T 9043T>G), V175L (9077G>T 9079A>G), V180I (9092G>A), A181S (9095G>T), E183S (9101G>A 9102A>G 9103A>T), S184E (9104A>G 9105G>A 9106T>G), S218A (9206T>G), A231V (9246C>T 9247T>A), V233I (9251G>A), V235L (9257G>C), D246E (9292T>G), Y247H (9293T>C), S250A (9302T>G), P252S (9308C>T), V261M (9335G>A 9337A>G), L264I (9344C>A 9346T>A), T265A (9347A>G), M267I (9355G>C), I272V (9368A>G 9370T>G), I275V (9377A>G 9379T>G), I280V (9392A>G 9394A>G), I284V (9404A>G), V290I (9422G>A 9424A>T), V293L (9431G>T 9433A>G), L297A (9443C>G 9444T>C), R303K (9462G>A 9463G>A), A307V (9474C>T), S312N (9489G>A 9490T>C), F317A (9503T>G 9504T>C), T319A (9509A>G 9511T>A), V328I (9536G>A), T332V (9548A>G 9549C>T), V334A (9555T>C), I345F (9587A>T), L353F (9611C>T 9613T>C), I363L (9641A>C), M366F (9650A>T 9652G>T), V367A (9654T>C 9655T>C), T370S (9662A>T 9664A>T), L372I (9668T>A 9670A>T), I379A (9689A>G 9690T>C 9691T>A), A380I (9692G>A 9693C>T 9694T>C), I382V (9698A>G 9700C>A), I383F (9701A>T 9703T>C), T387L (9713A>C 9714C>T 9715A>G), F390C (9723T>G), Y391H (9725T>C), S395N (9738G>A 9739T>C), K399R (9750A>G), R400K (9753G>A), V403M (9761G>A 9763C>G), S408T (9776T>A 9778C>A), D433E (9853T>G), V434T (9854G>A 9855T>C 9856G>A), M458L (9926A>T 9928G>A), S483A (10001T>G) | | | | | | | | |
| 3C-like proteinase (YP_009725301.1) | 1 | 306 | 100% | 2172.0 | 97.4% | 306 (100%) | 294 (96.1%) | 0/0/0/0 | 0 |

| | Begin | End | Coverage | Score | Concordance | Matches | Identities | I/D/M/F* | Stop Codons |
|---|---|---|---|---|---|---|---|---|---|
| | V35T (10157G>A 10158T>C), S46A (10190T>G 10192T>A), N65S (10248A>G 10249T>C), V86L (10310G>C 10312A>G), K88R (10317A>G), A94S (10334G>T 10336C>T), F134H (10454T>C 10455T>A 10456C>T), N180K (10594C>A), V202L (10658G>T 10660T>A), S267A (10853T>G 10855A>T), A285T (10907G>A), L286I (10910T>A 10912A>T) | | | | | | | | |
| nsp6 (YP_009725302.1) | 1 | 290 | 100% | 1813.0 | 89.9% | 290 (100%) | 253 (87.2%) | 0/0/0/0 | 0 |
| | S1G (10973A>G), A2K (10976G>A 10977C>A 10978A>G), V3F (10979G>T 10981G>C), R5K (10986G>A), T6I (10989C>T 10990A>T), I7V (10991A>G 10993C>T), L14M (11012T>A), I18F (11024A>T 11026T>C), V24I (11042G>A), L37V (11081T>G 11083G>T), A46T (11108G>A), M47L (11111A>C 11113G>T), I50M (11122T>G), M52I (11128G>T), S53A (11129T>G), F55C (11136T>G), M58L (11144A>C), F59L (11147T>C), D99E (11269T>A), M100L (11270A>T), V101A (11274T>C), F108Y (11295T>A), K109R (11298A>G), V120L (11330G>T), G135A (11376G>C), L146I (11408T>A 11410G>T), I168V (11474A>G 11476A>T), V182I (11516G>A), G188A (11535G>C), M192V (11546A>G), C197Y (11562G>A), I199L (11567A>T 11569T>G), F200L (11572C>A), F220C (11631T>G), T222C (11636A>T 11637C>G 11638T>C), N264S (11763A>G), V278I (11804G>A) | | | | | | | | |
| nsp7 (YP_009725303.1) | 1 | 83 | 100% | 508.0 | 99.4% | 83 (100%) | 82 (98.8%) | 0/0/0/0 | 0 |
| | K70R (12051A>G) | | | | | | | | |
| nsp8 (YP_009725304.1) | 1 | 198 | 100% | 1210.0 | 98.0% | 198 (100%) | 193 (97.5%) | 0/0/0/0 | 0 |
| | F15Y (12135T>A), I132V (12485A>G 12487A>C), N136G (12497A>G 12498A>G 12499C>T), T145N (12525C>A 12526A>C), S173N (12609G>A 12610T>C) | | | | | | | | |
| nsp9 (YP_009725305.1) | 1 | 113 | 100% | 752.0 | 98.4% | 113 (100%) | 110 (97.3%) | 0/0/0/0 | 0 |
| | T34N (12786C>A 12787A>T), T35S (12788A>T 12790A>G), L48H (12827T>C 12828T>A 12829A>C) | | | | | | | | |
| nsp10 (YP_009725306.1) | 1 | 139 | 100% | 1061.0 | 98.7% | 139 (100%) | 135 (97.1%) | 0/0/0/0 | 0 |
| | A23P (13091G>C), K113R (13362A>G), M137L (13433A>T), L138M (13436C>A 13438T>G) | | | | | | | | |
| RNA-dependent RNA polymerase (YP_009725307.1) | 1 | 932 | 100% | 6561.0 | 97.4% | 932 (100%) | 898 (96.4%) | 0/0/0/0 | 0 |
| | Q5S (13454C>T 13455A>C), S6T (13457T>A), D40E (13560T>A), D62E (13626T>A), D63G (13628A>G), I66L (13636A>T 13638T>A), F77M (13669T>A 13671C>G), L90V (13708C>G), K98V (13732A>G 13733A>T 13734A>C), I106V (13756A>G), A185S (13993G>T 13995T>A), N198D (14032A>G), I223V (14107A>G), T225V (14113A>G 14114C>T 14115C>A), T226A (14116A>G 14118G>A), S229C (14125A>T 14127T>C), V233I (14137G>A), T252A (14194A>G), V257M (14209G>A 14211T>G), T259A (14215A>G), T262A (14224A>G), Y265L (14233T>C 14234A>T 14235C>T), K281C (14281A>T 14282A>G 14283A>T), V299I (14335G>A), N611T (15272A>C 15273C>T), T643N (15368C>A 15369A>C), S647N (15380G>A), T739H (15655A>C 15656C>A 15657A>T), D740E (15660C>A), N743D (15667A>G), F766Y (15737T>A 15738C>T), T769N (15746C>A 15747T>C), S772A (15754T>G), S784A (15790T>G) | | | | | | | | |
| helicase (YP_009725308.1) | 1 | 601 | 100% | 4241.0 | 99.9% | 601 (100%) | 600 (99.8%) | 0/0/0/0 | 0 |
| | V570I (17944G>A 17946A>T) | | | | | | | | |
| 3'-to-5' exonuclease (YP_009725309.1) | 1 | 527 | 100% | 3864.0 | 96.8% | 527 (100%) | 501 (95.1%) | 0/0/0/0 | 0 |
| | V14I (18079G>A 18081A>C), T31I (18131C>T 18132T>A), E99D (18336A>T), P128E (18421C>G 18422C>A 18423T>A), D132E (18435T>A), S134T (18439T>A), S137N (18449G>A), N176G (18565A>G 18566A>G 18567T>A), R212K (18674G>A), A220S (18697G>T 18699T>A), H228N (18721C>A), I231V (18730A>G 18732T>G), L259Q (18815T>A 18816G>A), Y260H (18817T>C), T293S (18916A>T), I294V (18919A>G), K304R (18950A>G), I305V (18952A>G), A307S (18958G>T 18960G>T), A319S (18994G>T), D345E (19074T>A), S374H (19159T>C 19160C>A 19161T>C), S396A (19225T>G 19227C>A), V437T (19348G>A 19349T>C), L493Q (19516T>C 19517T>A), V510I (19567G>A) | | | | | | | | |
| endoRNAse (YP_009725310.1) | 1 | 346 | 100% | 2174.0 | 92.0% | 346 (100%) | 307 (88.7%) | 0/0/0/0 | 0 |
| | F7Y (19640T>A), Q18H (19674A>C), Q19A (19675C>G 19676A>C 19677G>C), V22A (19685T>C), T30A (19708A>G), V38I (19732G>A), L42I (19744T>A 19746G>C), V69I (19825G>A 19827G>T), D91E (19893T>A), I96V (19906A>G), S103T (19927T>A 19929T>A), T114S (19961C>G 19962G>T), I115A (19963A>G 19964T>C), A117S (19969G>T 19971A>T), P118S (19972C>T), F122L (19986T>G), D128E (20004T>A), Q152T (20074C>A 20075A>C), V155K (20083G>A 20084T>A 20085A>G), K158A (20092A>G 20093A>C), L162V (20104C>G 20106T>C), A171S (20131G>T 20133C>A), Y179F (20156A>T), V185I (20173G>A), V186I (20176G>A 20178C>T), N199D (20215A>G 20217T>C), Q201E (20221C>G 20223A>G), E202D (20226A>T), I211T (20252T>C), E223Q (20287G>C 20289A>G), S243G (20347A>G 20349T>A), L251M (20371C>A), F258S (20393T>C 20394T>A), K259Q (20395A>C 20397G>A), E260D (20400A>T), F263L (20407T>C), E264K (20410G>A), V314I (20560G>A), T325A (20593A>G 20595A>T) | | | | | | | | |
| 2'-O-ribose methyltransferase (YP_009725311.1) | 1 | 298 | 100% | 1968.0 | 95.3% | 298 (100%) | 278 (93.3%) | 0/0/0/0 | 0 |
| | S1A (20659T>G 20661T>A), D32E (20754T>A), S33N (20756G>A), T35V (20761A>G 20762C>T 20763A>T), L36I (20764T>A), K135R (21062A>G), N138H (21070A>C), I153L (21115A>C 21117T>G), Q158K (21130C>A 21132A>G), V167I (21157G>A 21159G>A), I169V (21163A>G), A188S (21220G>T), C209A (21283T>G 21284G>C), R216K (21304C>A 21305G>A 21306C>G), V223T (21325G>A 21326T>C), G265N (21451G>A 21452G>A), L272Y (21473T>A 21474A>T), S276E (21484A>G 21485G>A 21486T>A), I290V (21526A>G), V294I (21538G>A) | | | | | | | | |
| orf1a polyprotein (YP_009725295.1) | 1 | 4406 | 100% | 25296.0 | 84.3% | 4379 (99.3%) | 3552 (80.5%) | 4/27/0/0 | 1 |

| Begin | End | Coverage | Score | Concordance | Matches | Identities | I/D/M/F* | Stop Codons |
|---|---|---|---|---|---|---|---|---|

P6L (282C>T), F8V (287T>G), V38A (378T>C), Q44E (395C>G), D48N (407G>A), V56L (431G>C 433T>G), R77L (494C>T 495G>T), T78S (498C>G 499T>C), A79T (500G>A 502A>C), P80N (503C>A 504C>A), V84K (515G>A 516T>A 517T>G), M85V (518A>G 520G>C), L92M (539C>A 541C>G), E93D (544A>C), E102I (569G>A 570A>T 571G>A), I114T (606T>C 607A>C), V116I (611G>A 613G>T), K120N (625G>T), A138I (677G>A 678C>T), F143Y (693T>A), Y154I (725T>A 726A>T), F157Y (735T>A), Q158E (737C>G), E159Q (740G>C), S166G (761A>G), V169A (771T>C 772T>A), T170L (773A>C 774C>T), M174T (786T>C 787G>T), Y182V (809T>G 810A>T), E199D (862G>T), L204F (875C>T), A211S (896G>T 898T>A), S212M (899T>A 900C>T 901A>G), F221Y (927T>A 928T>C), D223E (934C>G), T224S (935A>T 937T>G), E233D (964A>C), Y241F (987A>T), E246D (1003A>T), L251H (1016T>C 1017T>A 1018G>C), L259S (1040T>A 1041T>G 1042G>T), N267K (1066T>A), N272K (1081T>G), I280K (1104T>A), I281V (1106A>G), T283V (1112A>G 1113C>T 1114T>C), L293T (1142C>A 1143T>C), D294E (1147T>G), N310Q (1193A>C 1195T>G), Q314N (1205C>A 1207A>T), C316H (1211T>C 1212G>A), D324N (1235G>A), G327D (1245G>A), T329V (1250A>G 1251C>T), G334C (1265G>T), V337L (1274G>C 1276T>G), F343H (1292T>C 1293T>A), T350V (1313A>G 1314C>T), K351I (1317A>T 1318A>T), A354P (1325G>C 1327C>T), Q362T (1349C>A 1350A>C 1351A>T), I368M (1369T>G), Y369P (1370T>C 1371A>C 1372T>A), H374Q (1387C>A), N375D (1388A>G 1390T>C), S376P (1391T>C), V378I (1397G>A 1399A>T), L384V (1415C>G), E386D (1423A>T), E390H (1433G>C 1435A>C), G392N (1439G>A 1440G>A), L393I (1442T>A 1444G>T), K394E (1445A>G), I396R (1451A>C 1452T>G 1453T>A), I404R (1476T>G 1477T>A), A405C (1478G>T 1479C>G 1480C>T), S412A (1499T>G 1501T>C), H417Y (1514C>T), C420R (1523T>C), N430D (1553A>G 1555C>T), C433S (1563G>C 1564T>A), N434G (1565A>G 1566A>G), V438I (1577G>A), V439T (1580G>A 1581T>C), E441D (1588A>C), G442N (1589G>A 1590G>A), S443V (1592T>G 1593C>T 1594C>G), G445T (1598G>A 1599G>C 1600T>C), D448E (1609C>G), N449D (1610A>G 1612C>T), Q455S (1628C>A 1629A>G 1630A>T), K456R (1631A>C 1632A>G 1633A>T), K458R (1637A>C 1638A>G 1639A>T), K468H (1667A>C 1669A>T), I473V (1682A>G 1684C>T), V488I (1727G>A 1729G>T), E489D (1732A>C), V491I (1736G>A 1738G>A), G493S (1742G>A), A498S (1757G>T 1759A>T), Q501T (1766C>A 1767A>C 1768A>C), F509Y (1791T>A), A516P (1811G>C 1813T>C), K517V (1814A>G 1815A>T), E525Q (1838G>C), K527R (1845A>G), I529V (1850A>G 1852A>T), S531T (1857G>C 1858T>A), Y534C (1866A>G), A535G (1869C>G 1870A>T), A537P (1874G>C 1876A>C), E539Q (1880G>C), R542G (1889C>G), V544I (1895G>A 1897A>C), S549A (1910T>G 1912C>G), E553D (1924A>T), T554A (1925A>G 1927T>A), Q556N (1931C>A 1933A>C), N557H (1934A>C 1936T>C), V559I (1940G>A 1942G>T), R560P (1944G>C), V561D (1947T>A), K564R (1956A>G 1957G>A), I567V (1964A>G 1966A>C), Q575E (1988C>G 1990G>A), Y576Q (1991T>C 1993T>G), I581V (2006A>G 2008T>C), M585V (2018A>G 2020G>T), F586Y (2022T>A 2023C>T), A591L (2036G>C 2037C>T 2038T>C), N594S (2046A>G), L595V (2048C>G 2050A>C), V596I (2051G>A), V597I (2054G>A 2056A>T), I601V (2066A>G 2068T>A), V605L (2078G>C), L608Q (2087T>C 2088T>A), T614S (2105A>T), I616L (2111A>C 2113C>T), F617L (2116T>G), V620T (2123G>A 2124T>C), Y621V (2126T>G 2127A>T), K625R (2139A>G 2140A>G), V627I (2144G>A), L628F (2147C>T), D629E (2152T>A), L631I (2156C>A), E633A (2163A>C), F635L (2168T>C), K636S (2172A>G 2173G>T), E637A (2175A>C), R643K (2193G>A 2194A>G), G645A (2199G>C), V649L (2210G>C 2212T>C), I652L (2219A>C), S653I (2222T>A 2223C>T 2224A>T), C655G (2228T>G), A656V (2232C>T), C657F (2235G>T), E658D (2239A>C), G661K (2246G>A 2247G>A 2248T>G), V665Q (2258G>C 2259T>A 2260C>G), T666V (2261A>G 2262C>T 2263C>T), C667A (2264T>G 2265G>C), A668S (2267G>T), K669D (2270A>G 2272G>T), E670N (2273G>A 2275A>C), E673D (2284G>T), S674C (2285A>T), Q676K (2291C>A 2293G>A), T677C (2294A>T 2295C>G 2296A>C), F679I (2300T>A), K680D (2303A>G 2305G>T), L681V (2306C>G), F685A (2318T>G 2319T>C 2320T>A), A687E (2325C>A 2326T>A), L688M (2327T>A), A690I (2333G>A 2334C>T), S692Q (2339T>C 2340C>A 2341T>A), I693V (2342A>G), I694T (2346T>C), G696A (2352G>C), K701R (2366A>C 2367A>G), A702S (2369G>T 2371C>A), T708V (2387A>G 2388C>T 2389A>C), V710I (2393G>A), T711A (2396A>G 2398G>T), H712Q (2401C>A), K719Q (2420A>C), V721I (2426G>A 2428T>A), K722R (2429A>C 2430A>G 2431A>T), S723G (2432T>G 2433C>G), R724K (2436G>A 2437A>G), E726Q (2441G>C 2443A>G), T727L (2444A>C 2445C>T 2446T>G), G728Q (2447G>C 2448G>A 2449C>A), I739V (2480A>G 2482T>A), I740T (2484T>C), E745D (2500A>T), T746S (2501A>T), L747H (2505T>A), P748D (2507C>G 2508C>A), E750V (2514A>T), V751L (2516G>C 2518G>T), L752T (2519T>A 2520T>C 2521A>C), T753S (2522A>T 2524A>T), T760N (2544C>A 2545T>C), D762E (2551T>A), Q764E (2555C>G), P765A (2558C>G), Q768T (2567C>A 2568A>C 2569A>G), T770V (2573A>G 2574C>T), S771D (2576A>G 2577G>A), E772S (2579G>A 2580A>G 2581A>C), A773F (2582G>T 2583C>T 2584T>C), V774T (2585G>A 2586T>C 2587T>A), E775N (2588G>A 2590A>T), A776G (2592C>G 2593T>A), P777A (2594C>G 2596A>T), L778I (2597T>A 2599G>C), I785V (2618A>G 2620T>A), T796K (2652C>A), K798Q (2657A>C 2659G>A), A803S (2672G>T 2674A>T), N805G (2678A>G 2679A>G), M806L (2681A>T 2683G>A), M807L (2684A>C), V808A (2688T>C 2689A>T), T812V (2699A>G 2700C>T), T814R (2705A>C 2706C>G 2707A>C), T821I (2727C>T 2728A>T), K822_V823insG (2731_2732insGGT), D827E (2746T>A), I831W (2756A>T 2757T>G 2758A>G), S838N (2778G>A), N840R (2784A>G 2785T>A), I849V (2810A>G), A859V (2841C>T), L864S (2855C>T 2856T>C), N869T (2871A>C), D877E (2896T>G), I880V (2903A>G 2905A>G), E888D (2929A>T), P892N (2939C>A 2940C>A 2941A>C), L893M (2942C>A), M902V (2969A>G 2971G>A), Y905F (2979A>T), E910D (2995G>T), S911A (2996T>G), F914E (3005T>G 3006T>A 3007T>A), K915N (3010A>C), L916F (3013G>T), A917S (3014G>T 3016T>A), H919R (3021A>G), D930E (3055T>A), E933D (3064A>C), G934D (3066G>A), G934_D935insA (3067_3068insGCA), D935E (3070T>G), F941I (3086T>A), E942D (3091G>T), P943E (3092C>G 3093C>A), S944T (3095T>A 3097A>C), T945C (3098A>T 3099C>G), Q946E (3101C>G), Y947H (3104T>C), K958L (3137A>C 3138A>T 3139A>C), T965S (3158A>T 3160T>A), S966A (3161T>G), A967E (3165C>A 3166T>A), A968T (3167G>A 3169T>A), L969V (3170C>G), Q970R (3174A>G), P971V (3176C>G 3177C>T), Q975E (3188C>G), D983_T999del (3212_3262delGATAGTCAACAAACTGTTGGTCAACAAGACGGCAGTGAGGACAATCAGACA), I1002E (3269A>G 3270T>A 3271T>G), T1004S (3275A>T), I1005_V1006del (3278_3283delATTGTT), V1008I (3287G>A), Q1009E (3290C>G 3292A>G), Q1011E (3296C>G), L1012P (3299T>C 3300T>C), E1013_M1014del (3302_3307delGAGATG), L1016P (3312T>C), V1019E (3321T>A 3322T>A), V1020E (3324T>A 3325T>A), Q1021_I1023del (3327_3335delAGACTATTG), E1024P (3327_3335delAGACTATTG 3336A>C), S1027Q (3344A>C 3345G>A 3346T>G), S1029T (3351G>C), Y1039A (3380T>G 3381A>C), N1042C (3389A>T 3390A>G), A1043V (3393C>T 3394A>T), E1047K (3404G>A 3406A>G), K1050Q (3413A>C), K1051S (3417A>G 3418G>T), V1052A (3420T>C 3421A>T), K1053N (3424A>T), T1055M (3429C>T 3430A>G), V1057I (3434G>A), V1063I (3452G>A 3454T>A), Y1064H (3455T>C), N1081G (3506A>G 3507A>G), V1085K (3518G>A 3519T>A 3520T>G), A1092K (3539G>A 3540C>A 3541T>G), T1093L (3542A>C 3543C>T 3544T>A), K1098T (3558A>C), V1104L (3575G>T 3577T>G), H1113K (3602C>A 3604C>G), V1122L (3629G>C 3631T>A), K1124A (3635A>G 3636A>C), S1133A (3662A>G 3663G>C 3664T>A), Q1140S (3683C>T 3684A>C 3685G>A), H1141Q (3688C>G), E1142D (3691A>C), V1143I (3692G>A 3694T>C), D1157K (3734G>A 3736C>A), I1159L (3740A>C 3742A>T), H1160Q (3745T>G), R1163Q (3752A>C 3753G>A), D1167Q (3764G>C 3766T>G), N1172Q (3779A>C 3781T>G), L1175I (3788T>A 3790A>T), F1178N (3797T>A 3798T>A), N1181A (3806A>G 3807A>C), D1184E (3817C>G), K1185Q (3818A>C 3820A>G), L1186V (3821C>G), S1188M (3827T>A 3828C>T 3829A>G), S1189D (3830A>G 3831G>A 3832C>T), F1190Y (3834T>A), E1192D (3841A>T), M1193N (3843T>A 3844G>C), K1194L (3845A>C 3846A>T), S1195K (3849G>A 3850T>G), E1196P (3851G>C 3852A>C 3853A>T), K1197R (3855A>G 3856G>A), Q1198V (3857C>G 3858A>T 3859A>G), V1199E (3861T>A 3862T>A), E1200A (3864A>C), Q1201P (3867A>C 3868A>T), I1203Q (3872A>C 3873T>A 3874C>A), A1204E (3876C>A 3877T>G), I1206P (3881A>C 3882T>C 3883T>A), K1208N (3889A>C), E1209T (3890G>A 3891A>C 3892G>A), V1211D (3897T>A), K1212_P1213del (3899_3904delAAGCCA), F1214S (3906T>C 3907T>C), I1215K (3909T>A), S1218E (3917A>G 3918G>A 3919T>G), P1220del (3923_3925delCCT), E1223V (3933A>T), R1225K (3939G>A 3940A>G), K1226P (3941A>C 3942A>C 3943A>T), Q1227V (3944C>G 3945A>T 3946A>C), D1229V (3951A>T 3952T>G), K1230_K1231insP (3955_3956insCCA), V1236I (3971G>A), E1237D (3976A>T), E1251N (4016G>A 4018A>T), N1252K (4021C>G), Y1256F (4032A>T), I1257A (4034A>G 4035T>C), N1262K (4051T>G), H1264Y (4055C>T 4057T>C), P1265H (4059C>A 4060A>T), A1268Q (4067G>C 4068C>A 4069C>G), T1269N (4071C>A 4072T>C), L1270M (4073C>A 4075T>G), V1271L (4076G>C), S1272R (4081T>A), D1273G (4083A>G 4084C>T), I1274E (4085A>G 4086T>A 4087T>A), I1276M (4093C>G), T1277S (4094A>T), K1280E (4103A>G), I1286M (4123A>G), V1291I (4136G>A 4138T>C), Q1292T (4139C>A 4140A>C 4141A>T), E1293S (4142G>A 4143A>G 4144G>T), V1295D (4149T>A), L1296I (4151T>A 4153A>C), A1298C (4157G>T 4158C>G), T1303S (4172A>T 4174T>C), A1314S (4205G>T 4207G>A), K1315R (4209A>G), R1318K (4218G>A 4219A>G), T1322V (4229A>G 4230C>T 4231A>T), N1324E (4235A>G 4237T>G), L1334C (4266T>G 4267A>T), N1335A (4268A>G 4269A>C), V1339L (4280G>C 4282A>T), V1345A (4299T>C 4300G>T), I1355V (4328A>G 4330T>A), I1359E (4340A>G 4341T>A 4342T>A), I1360A (4343A>G 4344T>C 4345C>A), S1361P (4346T>C), E1363A (4353A>C 4354G>T), Q1365E (4358C>G), V1391I (4436G>A 4438C>A), V1393M (4442G>A), E1394D (4447A>T), T1395V (4448A>G 4449C>T), K1396R (4452A>G), V1399M (4460G>A 4462T>G), S1400A (4463T>G), V1415I (4508G>A 4510G>C), A1420V (4524C>T 4525T>C), Y1423F (4533A>T), T1429E (4550A>G 4551C>A 4552A>G), T1430P (4553A>C), L1434I (4565C>A), N1436T (4572A>C 4573C>G), T1437K (4575C>A 4576A>G), D1440S (4583G>T 4584A>C), T1444P (4595A>C 4597T>G), L1450I (4613C>A), L1457F (4636A>T), Y1465C (4659A>G), V1471A (4677T>C 4678G>T), T1474V (4685A>G 4686C>T), A1485T (4718G>A 4720G>A), P1497S (4754C>T), I1502V (4769A>G 4771T>A), I1505V (4778A>G 4780C>T), K1512R (4800A>G), S1520R (4823T>C 4824C>G), Q1522E (4829C>G 4831A>G), I1525V (4838A>G 4840A>T), S1534I (4866G>T), Y1537H (4874T>C), T1538_S1539insL (4879_4880insCTG), S1539E (4880A>G 4881G>A 4882T>G), N1540S (4884A>G 4885T>C), T1542V (4889A>G 4890C>T), T1543E (4892A>G 4893C>A 4894A>G), I1551L (4916A>C 4918C>T), T1552S (4919A>T 4921C>A), F1553L (4922T>C), N1555K (4930T>A), T1558S (4938C>G 4939A>T), R1566K (4962G>A), I1577T (4995T>C), V1583L (5012G>C), N1611V (5096A>G 5097A>T 5098T>A), S1612N (5099T>A 5100C>A 5101A>T), Y1619F (5121A>T), N1623S (5133A>G), V1629S (5150G>A 5151T>G), T1638L (5177A>C 5178C>T), P1640E (5183C>G 5184C>A 5185T>G), Y1658F (5238A>T 5239C>T), N1662G (5249A>G 5250A>G), A1677S (5294G>T 5296C>T), T1678S (5298C>G), A1679V (5301C>T 5302A>T), T1682A (5309A>G), I1686L (5321A>C 5323A>T), L1688V (5327T>G 5329G>C), P1692A (5339C>G), D1697E (5356T>G), E1706D (5383A>T), C1718S (5417T>A), S1733T (5463G>C 5464T>C), Y1734H (5465T>C 5467C>T), F1736L (5471T>C 5473T>A), D1742E (5491T>A), C1744A (5495T>G 5496G>C 5497C>A), T1754H (5525A>C 5526C>A), Q1758K (5537C>A 5539G>A), Q1759T (5540C>A 5541A>C 5542G>T), K1763T (5553A>C), E1777D (5596A>T), Q1778N (5597C>A 5599A>T), F1779L (5600T>C), K1781T (5607A>C), Q1784S (5615C>T 5616A>C 5617G>C), T1788V (5627A>G 5628C>T), K1791R (5636A>C 5637A>G 5638A>T), Q1792D (5639C>G 5641A>T), K1795Q (5648A>C), P1803S (5672C>T), Q1813E (5702C>G), E1815K (5708G>A), K1817Q (5714A>C), H1818Q (5719T>A), T1822L (5729A>T 5730C>T 5731T>A), S1825N (5739G>A), K1837T (5775A>C 5776A>T), S1841A (5786T>G), C1847R (5804T>C 5806C>T), L1852H (5819T>C 5820T>A 5821A>C), S1856M (5831T>A 5832C>T 5833C>G), I1863V (5852A>G 5854T>G), N1871T (5877A>C 5878C>A), T1881S (5906A>T 5908T>G), V1888T (5927G>A 5928T>C), C1889Y (5931G>A 5932T>C), D1893E (5944C>A), N1898G (5957A>G 5958A>G 5959T>G), S1905A (5978T>G), F1907Y (5985T>A 5986C>T), N1917T (6015A>C 6016C>T), Y1920L (6024A>T 6025T>A), F1930L (6053T>C 6055T>C), V1931T (6056G>A 6057T>C), D1933S (6062G>T 6063A>C), I1935T (6069T>C 6070C>A), L1944M (6095T>A 6097A>G), Y1947F (6105A>T 6106T>C), K1948T (6108A>C 6109G>A), K1956S (6131A>T 6132A>C 6133A>T), K1973R (6183A>G), T1976S (6191A>T), P1977A (6194C>G 6196C>G), V1994I (6245G>A), N1996Q (6251A>C 6253T>G), N1999T (6261A>C 6262T>C), A2001T (6266G>A 6268C>A), Y2003F (6273A>T 6274T>C), I2010L (6293A>T), E2020D (6325A>T), D2026E (6343T>A), K2029A (6350A>G 6351A>C 6352G>A), S2030V (6353T>G 6354C>T), A2033T (6362G>A 6364G>A), D2043S (6392G>A 6393A>G), L2044Q (6396T>A), K2045Q (6398A>C), V2047T (6404G>A 6405T>C), D2060E (6445C>A), L2062I (6449C>A 6451T>A), N2065D (6458A>G 6460T>C), D2074N (6485G>A 6487C>T), I2075V (6488A>G 6490T>C), A2080S (6503G>T), N2081D (6506A>G), N2082E (6509A>G 6511T>A), S2083G (6512A>G), L2084V (6515T>G 6517A>T), I2086V (6521A>G 6523T>A), E2088Q (6527G>C), V2090L (6533G>T 6535T>A), T2093E (6542A>G 6543C>A 6544A>G), D2101E (6568C>A), S2103T (6572T>A 6574T>A), L2105I (6578C>A), R2115L (6608A>C 6609G>T), V2116A (6612T>C 6613A>C), L2122I (6629C>A), L2127I (6644T>A 6646A>T), V2130I (6653G>A), D2136S (6671G>A 6672A>G), T2137K (6675C>A 6676T>A), A2139L (6680G>T 6681C>T 6682T>G), N2140A (6683A>G 6684A>C), A2142V (6690C>T 6691T>C), N2147G (6704A>G 6705A>G 6706C>A), K2148Q (6707A>C), V2149A (6711T>C 6712T>A), V2150A (6714T>C 6715T>A), S2151I (6717G>T), T2154S (6725A>T 6727T>A), I2156C (6731A>T 6732T>G 6733A>C), V2157A (6735T>C), T2158K (6738C>A 6739A>G), C2160L (6744G>T 6745T>A), L2161A (6746T>G 6747T>C), N2162Q (6749A>C 6751C>A), C2165F (6759G>T), T2166N (6762C>A 6763T>C), F2172V (6779T>G 6781C>G), L2177F (6794C>T 6796A>C), R2184K (6816G>A), K2191R (6837A>G), M2194L (6845A>C 6847G>A), T2202S (6870C>G), G2207A (6885G>C), F2209L (6892T>A), E2212D (6901G>T), S2214G (6905T>G 6906C>G 6907A>C), F2215I (6908T>A), L2218V (6917T>G), N2222K (6931T>A), I2227F (6944A>T 6946A>C), N2228T (6948A>C 6949T>A), I2230A (6953A>G 6954T>C 6955A>T), I2231M (6958T>G), F2233L (6962T>C 6964T>A), V2238I (6977G>A), Y2245C (6999A>G 7000C>T), S2246V (7001T>G 7002C>T), L2250F (7015A>T), M2254L (7025A>T 7027G>A), L2257F (7036A>T), M2259A (7040A>G 7041T>C 7042G>T), T2264N (7056C>A), Y2266V (7061T>G 7062A>T 7063C>T), G2269L (7070G>T 7071G>T 7072C>G), T2274S (7085A>T), I2278T (7098T>C), A2279M (7100G>A 7101C>T 7102A>G), T2280D (7103A>G 7104C>A 7105C>T), Y2281F (7107A>T), T2283E

| | | | | | | | | | |
|---|---|---|---|---|---|---|---|---|---|
| | (7112A>G 7113C>A 7114T>A), I2286F (7121A>T 7123A>T), V2290I (7133G>A), T2300S (7163A>T 7165C>T), S2303A (7172T>G), I2309V (7190A>G 7192T>G), F2314Y (7206T>A 7207T>C), W2316L (7211T>C 7212G>T 7213G>A), A2320I (7223G>A 7224C>T), F2321L (7228T>A), V2324A (7236T>C 7237T>C), F2328V (7247T>G), I2332M (7261T>G), R2336K (7272G>A 7273G>A), V2340L (7283G>T), A2344S (7295G>T 7297T>A), L2349V (7310T>G), S2352G (7319A>G), V2356S (7331G>A 7332T>G 7333A>T), L2367F (7366A>T), N2370S (7374A>G), L2371I (7376C>A), I2377V (7394A>G), V2393I (7442G>A), V2400I (7463G>A 7465T>C), V2401M (7466G>A 7468A>G), N2405T (7479A>C 7480T>C), V2430M (7553G>A 7555T>G), R2431K (7557G>A 7558A>G), K2442R (7589A>C 7590A>G 7591A>T), L2447T (7604C>A 7605T>C 7606A>T), V2453L (7622G>C 7624T>C), A2460T (7643G>A), T2495A (7748A>G 7750A>T), S2500A (7763T>G 7765C>G), I2501L (7766A>C 7768C>T), S2517P (7814T>C 7816T>G), E2550D (7915A>C), S2553A (7922T>G), A2554S (7925G>T 7927A>T), A2584T (8015G>A 8017G>T), A2587S (8024G>T 8026A>C), N2596D (8051A>G 8053T>C), S2600A (8063T>G), N2603S (8073A>G 8074C>T), T2611A (8096A>G), E2617H (8114G>C 8116A>C), A2618S (8117G>A 8118C>G 8119T>C), N2623G (8132A>G 8133A>G), S2625A (8138T>G 8140C>T), N2628G (8147A>G 8148A>G), I2634V (8165A>G 8167T>G), F2641V (8186T>G), S2644T (8195T>A 8197A>C), E2647D (8206A>C), V2652I (8219G>A), Q2660H (8245A>C), I2663L (8252A>T), Y2673F (8283A>T 8284T>C), S2695N (8349G>A), I2709V (8390A>G), A2710S (8393G>T 8395T>A), F2718Y (8418T>A), L2738I (8477T>A), K2741R (8487A>G 8488G>A), V2754I (8525G>A), A2759S (8540G>T), N2767S (8565A>G), N2768T (8568A>C), W2769C (8572G>T), L2770F (8575G>T), Q2772L (8580A>T 8581G>T), L2773M (8582T>A 8584A>G), I2774L (8585A>C), V2776A (8592T>C 8593T>C), V2779L (8600G>T), F2780C (8604T>G), L2781V (8606C>G), F2782L (8609T>C), V2783A (8613T>C), A2785L (8618G>T 8619C>T 8620T>G), I2786V (8621A>G), F2787C (8625T>G 8626C>T), L2789I (8630T>A 8632A>C), I2790V (8633A>G 8635A>T), T2791M (8637C>T 8638A>G), V2795T (8648G>A 8649T>C 8650C>A), M2796L (8651A>T), K2798I (8658A>T 8659A>C), T2800D (8663A>G 8664C>A), D2801G (8667A>G 8668C>T), F2802Y (8670T>A 8671T>C), S2803T (8672T>A), S2804N (8676G>A), D2813Q (8702G>C 8704T>G), G2814D (8706G>A), A2821I (8726G>A 8727C>T 8728A>T), T2825D (8738A>G 8739C>A), D2833G (8763A>G), T2836A (8771A>G), T2846K (8802C>A 8803T>A), A2850S (8813G>A 8814C>G 8815T>C), L2853V (8822T>G 8824G>A), I2854V (8825A>G 8827T>A), V2857I (8834G>A), V2862I (8849G>A 8851G>T), V2865I (8858G>A 8860C>A), I2873V (8882A>G 8884A>G), T2876A (8891A>G), T2877I (8895C>T 8896T>C), T2906S (8982C>G), S2926M (9041T>A 9042C>T 9043T>G), V2938L (9077G>T 9079A>G), V2943I (9092G>A), A2944S (9095G>T), E2946S (9101G>A 9102A>G 9103A>T), S2947E (9104A>G 9105G>A 9106T>G), S2981A (9206T>G), A2994V (9246C>T 9247T>A), V2996I (9251G>A), V2998L (9257G>C), D3009E (9292T>G), Y3010H (9293T>C), S3013A (9302T>G), P3015S (9308C>T), V3024M (9335G>A 9337A>G), L3027I (9344C>A 9346T>A), T3028A (9347A>G), M3030I (9355G>C), I3035V (9368A>G 9370T>G), I3038V (9377A>G 9379T>G), I3043V (9392A>G 9394A>G), I3047V (9404A>G), V3053I (9422G>A 9424A>T), V3056L (9431G>T 9433A>G), L3060A (9443C>G 9444T>C), R3066K (9462G>A 9463G>A), A3070V (9474C>T), S3075N (9489G>A 9490T>C), F3080A (9503T>G 9504T>C), T3082A (9509A>G 9511T>A), V3091I (9536G>A), T3095V (9548A>G 9549C>T), V3097A (9555T>C), I3108F (9587A>T), L3116F (9611C>T 9613T>C), I3126L (9641A>C), M3129F (9650A>T 9652G>T), V3130A (9654T>C 9655T>C), T3133S (9662A>T 9664A>T), L3135I (9668T>A 9670A>T), I3142A (9689A>G 9690T>C 9691T>A), A3143I (9692G>A 9693C>T 9694T>C), I3145V (9698A>G 9700C>A), I3146F (9701A>T 9703T>C), T3150L (9713A>C 9714C>T 9715A>G), F3153C (9723T>G), Y3154H (9725T>C), S3158N (9738G>A 9739T>C), K3162R (9750A>G), R3163K (9753G>A), V3166M (9761G>A 9763C>G), S3171T (9776T>A 9778C>A), D3196E (9853T>G), V3197T (9854G>A 9855T>C 9856G>A), M3221L (9926A>T 9928G>A), S3246A (10001T>G), V3298T (10157G>A 10158T>C), S3309A (10190T>G 10192T>A), N3328S (10248A>G 10249T>C), V3349L (10310G>C 10312A>G), K3351R (10317A>G), A3357S (10334G>T 10336C>T), F3397H (10454T>C 10455T>A 10456C>T), N3443K (10594C>A), V3465L (10658G>T 10660T>A), S3530A (10853T>G 10855A>T), A3548T (10907G>A), L3549I (10910T>A 10912A>T), S3570G (10973A>G), A3571K (10976G>A 10977C>A 10978A>G), V3572F (10979G>T 10981G>C), R3574K (10986G>A), T3575I (10989C>T 10990A>T), I3576V (10991A>G 10993C>T), L3583M (11012T>A), I3587F (11024A>T 11026T>C), V3593I (11042G>A), L3606V (11081T>G 11083G>T), A3615T (11108G>A), M3616L (11111A>C 11113G>T), I3619M (11122T>G), M3621I (11128G>T), S3622A (11129T>G), F3624C (11136T>G), M3627L (11144A>C), F3628L (11147T>C), D3668E (11269T>A), M3669L (11270A>T), V3670A (11274T>C), F3677Y (11295T>A), K3678R (11298A>G), V3689L (11330G>T), G3704A (11376G>C), L3715I (11408T>A 11410G>T), I3737V (11474A>G 11476A>T), V3751I (11516G>A), G3757A (11535G>C), M3761V (11546A>G), C3766Y (11562G>A), I3768L (11567A>T 11569T>G), F3769L (11572C>A), F3789C (11631T>G), T3791C (11636A>T 11637C>G 11638T>C), N3833S (11763A>G), V3847I (11804G>A), K3929R (12051A>G), F3957Y (12135T>A), I4074V (12485A>G 12487A>C), N4078G (12497A>G 12498A>G 12499C>T), T4087N (12525C>A 12526A>C), S4115N (12609G>A 12610T>C), T4174N (12786C>A 12787A>T), T4175S (12788A>T 12790A>G), L4188H (12827T>C 12828T>A 12829A>C), A4276P (13091G>C), K4366R (13362A>G), M4390L (13433A>T), L4391M (13436C>A 13438T>G), Q4397S (13454C>T 13455A>C), S4398T (13457T>A) | | | | | | | | |
| nsp11 (YP_009725312.1) | 1 | 13 | 100% | 71.0 | 86.6% | 13 (100%) | 11 (84.6%) | 0/0/0/0 | 0 |
| | Q5S (13454C>T 13455A>C), S6T (13457T>A) | | | | | | | | |
| surface glycoprotein (YP_009724390.1) | 1 | 1274 | 100% | 7176.0 | 82.1% | 1250 (97.7%) | 974 (76.1%) | 6/24/0/0 | 1 |
| | V3I (21569G>A), V6L (21578G>T 21580T>A), L7F (21583A>T), P9T (21587C>A 21589A>T), V11T (21593G>A 21594T>C 21595C>T), S13G (21599A>G), S13_Q14insSDLD (21601_21602insAGTGACCTTGAC), Q14R (21603A>G), V16T (21608G>A 21609T>C 21610T>C), N17T (21612A>C), L18F (21614C>T), T19D (21617A>G 21618C>A 21619A>T), T20D (21620A>G 21621C>A 21622C>T), R21del (21623_21625delAGA), T22V (21626A>G 21627C>T), L24del (21632_21634delTTA), P25A (21635C>G 21637C>T), A27N (21641G>A 21642C>A 21643A>T), T29_N30insQH (21649_21650insCAACAT), N30T (21651A>C), F32S (21657T>C 21658C>T), T33M (21660C>T 21661A>G), K41E (21683A>G), V42I (21686G>A), S46D (21698T>G 21699C>A 21700A>C), V47T (21701G>A 21702T>C), H49Y (21707C>T), S50L (21711C>T), F59Y (21738T>A), W64G (21752T>G), A67T (21761G>A), H69_T73del (21767_21781delCATGTCTCTGGGACC), G75H (21785G>C 21786G>A), K77_R78del (21790_21795delTAAGAG), D80G (21801A>G 21802T>C), L84I (21812C>A), N87K (21823T>G), V90I (21830G>A), S94A (21842T>G), I100V (21860A>G 21862A>T), I101V (21863A>G 21865A>C), I105V (21875A>G), T108S (21884A>T), L110M (21890T>A 21892A>G), D111N (21893G>A 21895T>C), S112N (21896T>A 21897C>A 21898G>C), T114S (21902A>T 21904C>A), L117V (21911C>G 21913A>G), L118I (21914C>A), V120I (21920G>A), A123S (21929G>T), K129R (21947A>C 21948A>G), V130A (21951T>C 21952C>A), E132N (21956G>A 21958A>C), Q134E (21962C>G), F135L (21967T>G), N137D (21971A>G 21973T>C), D138N (21974G>A 21976T>C), L141F (21985G>T), G142A (21987G>C), Y144S (21993A>C), Y145K (21995T>A 21997C>A), H146P (21999A>C), K147M (22002A>T 22003A>G), N148_S151del (22004_22015delAACAACAAAAGT), W152G (22016T>G 22018G>T), M153T (22020T>C 22021G>A), E154Q (22022G>C 22024A>G), S155T (22026G>C 22027T>A), E156H (22028G>C 22030G>T), F157T (22031T>A 22032T>C 22033C>T), R158M (22035G>T 22036A>G), V159I (22037G>A 22039T>A), Y160F (22041A>T 22042T>C), S161D (22043T>G 22044C>A), S162N (22047G>A), N164F (22052A>T 22053A>T), V171I (22073G>A 22075C>A), Q173D (22079C>G 22081G>T), P174A (22082C>G 22084T>C), L176S (22088C>T 22089T>C 22090T>G), M177L (22091A>C 22093G>T), L179V (22097C>G), E180S (22100G>T 22101A>C), G181E (22104G>A), Q183S (22109C>T 22110A>C 22111G>A), N188H (22124A>C 22126T>C), I197K (22152T>A 22153T>A), Y200F (22161A>T), F201L (22163T>C 22165T>C), K202Y (22166A>T 22168A>T), I203V (22169A>G 22171A>T), S205K (22175T>A 22176C>A 22177T>G), K206G (22178A>G 22179A>G 22180G>C), H207Y (22181C>T 22183C>T), T208Q (22184A>C 22185C>A 22186G>A), N211D (22193A>G), L212V (22196T>G), Q218S (22214C>T 22215A>C 22216G>T), S221N (22223T>A 22224C>A 22225G>C), A222T (22226G>A), E224K (22232G>A), L226I (22238T>A 22240G>T), V227F (22241G>T 22243A>T), D228K (22244G>A 22246T>G), I231L (22253A>C 22255A>T), R237N (22272G>A 22273G>T), Q239R (22277C>A 22278A>G), T240A (22280A>G 22282T>C), L241I (22283T>A 22285A>T), A243T (22289G>A 22291T>A), L244A (22292T>G 22293T>C 22294A>C), H245_Y248del (22295_22306delCATAGAAGTTAT), L249F (22309G>T), T250S (22310A>T 22312T>A), G252A (22317G>C), D253Q (22319G>C 22321T>A), S254D (22322T>G 22323C>A 22324T>C), S255I (22325T>A 22326C>T), S256_G257del (22328_22333delTCAGGT), T259G (22337A>G 22338C>G 22339A>C), A260T (22340G>A 22342T>G), G261S (22343G>T 22344G>C 22345T>A), Y266F (22359A>T), Q271K (22373C>A 22375A>G), R273T (22380G>C 22381G>T), L276M (22388C>A 22390A>G), N280D (22400A>G), A292S (22436G>T 22438A>T), L293Q (22440T>A 22441T>A), D294N (22442G>A 22444C>T), S297A (22451T>G 22453A>T), T299L (22457A>C 22458C>T 22459A>C), T302S (22466A>T 22468G>T), L303V (22469T>G 22471G>T), T307E (22481A>G 22482C>A 22483T>G), V308I (22484G>A 22486A>T), E309D (22489A>C), Q321V (22523C>G 22524A>T 22525A>T), T323S (22529A>T), E324G (22533A>G), S325D (22535T>G 22536C>A), I326V (22538A>G), R346K (22599G>A), A348P (22604G>C 22606A>T), N354E (22622A>G 22624C>G), R357K (22632G>A), A372T (22676G>A), S373F (22680C>T 22681A>T), P384A (22712C>G 22714T>C), T393S (22739A>T 22741T>C), I402V (22766A>G 22768T>C), R403K (22770G>A 22771A>G), E406D (22780A>T), K417V (22811A>G 22812A>T 22813G>T), T430M (22851C>T 22852A>G), I434L (22862A>C 22864A>T), S438T (22874T>A), N439R (22878A>G 22879C>G), L441I (22883C>A), S443A (22889T>G), K444T (22893A>C 22894G>T), V445S (22895G>T 22896T>C 22897T>A), G446T (22898G>A 22899G>C), L452K (22916C>A 22917T>A 22918G>A), L455Y (22926T>A 22927G>T), F456L (22928T>C), K458H (22934A>C 22936G>T), S459G (22937T>G 22938C>G 22939T>C), N460K (22942T>G), K462R (22947A>G 22948A>G), T470N (22971C>A), E471V (22974A>T 22975A>G), I472P (22976A>C 22977T>C 22978C>T), Y473F (22980A>T 22981T>C), Q474S (22982C>T 22983A>C 22984G>C), A475P (22985G>C 22987C>T), G476D (22989G>A), S477G (22991A>G), T478K (22995C>A), N481T (23004A>C 23005T>C), G482P (23006G>C 23007G>C 23008T>A), V483del (23009_23011delGTT), E484P (23012G>C 23013A>C 23014A>T), G485A (23016G>C), F486L (23018T>C), F490W (23031T>G 23032T>G), Q493N (23039C>A 23041A>T), S494D (23042T>G 23043C>A 23044A>T), Q498Y (23054C>T 23056A>C), P499T (23057C>A), N501T (23064A>C), V503I (23069G>A), H519N (23117C>A), K529L (23147A>T 23148A>T 23149G>A), N532D (23156A>G 23158T>C), V534I (23162G>A), K537Q (23171A>C 23173A>G), E554P (23222G>C 23223A>C 23224G>T), N556S (23228A>T 23229A>C 23230C>A), K558R (23235A>G 23236G>A), L560Q (23241T>A 23242G>A), I569V (23267A>G), A570S (23270G>T), T572F (23276A>T 23277C>T 23278T>C), A575S (23285G>T 23287T>C), Q580K (23300C>A 23302G>A), L582S (23306C>T 23307T>C), T588S (23324A>T), S591A (23333T>G), T604A (23372A>G), N606S (23378A>T 23379A>C 23380C>T), Q607E (23381C>G 23383G>A), E619D (23419A>T), P621S (23423C>T), V622T (23426G>A 23427T>C 23428T>A), T632A (23456A>G), V635I (23465G>A 23467T>A), S640N (23480T>A 23481C>A 23482T>C), R646Q (23499G>A 23500T>A), N657D (23531A>G), N658T (23535A>C 23536C>T), Q675H (23587G>T), Q677V (23591C>G 23592A>T 23593G>T), T678S (23594A>T), N679L (23597A>T 23598A>T 23599T>A), S680_R683del (23601_23612delCTCCTCGGCGGG), A684L (23601_23612delCTCCTCGGCGGG 23613C>T), V687T (23621G>A 23622T>C 23623A>T), A688S (23624G>A 23625C>G 23626T>C), S689Q (23627A>C 23628G>A 23629T>A), Q690K (23630C>A), I693V (23639A>G 23641T>G), E702D (23668A>T), N703S (23670A>G), V705I (23675G>A), S711T (23693T>A 23695T>C), T719S (23717A>T 23719T>A), V722I (23726G>A), I726V (23738A>G 23740T>A), L727M (23741C>A 23743A>G), T732A (23756A>G 23758C>T), T739N (23778C>A 23779A>T), S750A (23810A>G 23811G>C 23812C>T), T768S (23864A>T 23866T>A), V772A (23877T>C), K776R (23888A>C 23889A>G 23890A>C), Q779R (23898A>G 23899A>T), I788M (23926T>G), P793T (23939C>A 23941A>T), I794L (23942A>T 23944T>G), D796Y (23948G>T), S810L (23990T>C 23991C>T), S813T (24000G>C 24001C>T), I834M (24064C>G), D839E (24079T>A), A845N (24095G>A 24096C>A), E868D (24166A>T), Q872A (24176C>G 24177A>C 24178A>C), S875A (24185T>G), L878V (24194T>G 24196A>T), A879S (24197G>A 24198C>G 24199G>T), I882A (24206A>G 24207T>C), S884A (24212T>G), L922Q (24326T>C 24327T>A 24328G>A), S929K (24348G>A 24349T>G), G932S (24356G>A 24358C>T), K933Q (24359A>C), D936E (24370C>A), S939T (24377T>A 24379T>A), S940T (24380T>A 24382C>A), A942S (24386G>T), S943T (24390G>C), S1055A (24725T>G), A1070S (24770G>T 24772A>C), K1073R (24780A>G), D1084E (24814T>A), H1088Y (24824C>T), S1097F (24852C>T 24853A>T), H1101S (24863C>T 24864A>C 24865C>T), V1104I (24872G>A 24874A>T), Y1110F (24891A>T), E1111S (24893G>T 24894A>C 24895A>T), V1133I (24959G>A 24961C>T), I1216V (25208A>G), M1233L (25259A>T), C1247A (25301T>G 25302G>C 25303T>A) | | | | | | | | |

| | Begin | End | Coverage | Score | Concordance | Matches | Identities | I/D/M/F* | Stop Codons |
|---|---|---|---|---|---|---|---|---|---|
| ORF3a protein (YP_009724391.1) | 1 | 276 | 100% | 1490.0 | 76.7% | 275 (99.6%) | 200 (72.5%) | 0/1/0/0 | 1 |
| | I7F (25411A>T 25413C>T), I10L (25420A>C), G11R (25423G>A), T12S (25426A>T 25428T>A), V13I (25429G>A 25431A>T), L15A (25435T>G 25436T>C 25437G>A), K16Q (25438A>C), Q17P (25442A>C), G18V (25445G>T 25446T>A), E19K (25447G>A), K21D (25453A>G 25455G>C), D22N (25456G>A), T24S (25462A>T), S26A (25468T>G), D27S (25471G>A 25472A>G), F28T (25474T>A 25475T>C), R30H (25481G>A 25482C>T), I37L (25501A>C), I47V (25531A>G), V48I (25534G>A), L52F (25546C>T), S60T (25570T>A), T64A (25582A>G 25584C>G), K66N (25590A>T), S74Y (25613C>A 25614C>T), V77F (25621G>T 25623T>C), H78Q (25626C>G), V80I (25630G>A), V90I (25660G>A 25662T>C), L101M (25693C>A 25695T>G), P104Q (25703C>A 25704T>A), V112I (25726G>A 25728C>A), S117C (25741A>T 25743T>C), F120A (25750T>G 25751T>C 25752T>A), V121C (25753G>T 25754T>G 25755A>T), L127C (25771C>T 25772T>G), R134K (25792C>A 25793G>A 25794T>A), L147V (25831C>G), N152H (25846A>C), C153N (25849T>A 25850G>A 25851T>C), S165D (25885T>G 25886C>A), S166T (25888T>A), I169V (25897A>G), S171E (25903T>G 25904C>A), T175I (25916C>T 25917A>T), T176S (25918A>T), S177T (25922G>C 25923T>A), I179K (25928T>A 25929T>A), S180L (25930T>C 25931C>T 25932T>C), E181K (25933G>A), H182E (25936C>G 25938T>A), T190S (25960A>T), K192D (25966A>G 25968A>T), W193R (25969T>A), E194H (25972G>C 25974A>C), C200Y (25991G>A), L203V (25999T>G), S205G (26005A>G 26007T>C), S209E (26017T>G 26018C>A), D210V (26021A>T 26022C>T), Y215E (26035T>G 26037C>G), L219I (26047T>A 26049G>T), S220T (26051G>C), V225I (26065G>A), H227N (26071C>A), V228A (26075T>C), Y233F (26090A>T 26091C>T), I236L (26098A>C), D238K (26104G>A 26106T>A), E239D (26109G>C), E241P (26113G>C 26114A>C 26115A>G), E242del (26116_26118delGAA), H243N (26119C>A), V256A (26159T>C), V259A (26168T>C), E261D (26175A>T) | | | | | | | | |
| envelope protein (YP_009724392.1) | 1 | 76 | 100% | 447.0 | 94.3% | 76 (98.7%) | 73 (94.8%) | 1/0/0/0 | 1 |
| | S55T (26407T>A 26409T>G), F56V (26410T>G), S68_R69insE (26448_26449insGAA), R69G (26449A>G) | | | | | | | | |
| membrane glycoprotein (YP_009724393.1) | 1 | 223 | 100% | 1419.0 | 92.9% | 222 (99.6%) | 202 (90.6%) | 0/1/0/0 | 1 |
| | S4del (26531_26533delTTC), K15Q (26565A>C 26567G>A), T30A (26610A>G 26612A>C), C33M (26619T>A 26620G>T 26621T>G), A40S (26640G>T 26642C>T), I52V (26676A>G), I76V (26748A>G 26750C>G), L87I (26781C>A), I97V (26811A>G), H125R (26896A>G 26897T>G), L129V (26907C>G), L134M (26922C>A 26924A>G), L145I (26955C>A), I151M (26975T>G), H155S (26985C>T 26986A>C 26987T>C), A188G (27085C>G 27086A>C), G189T (27087G>A 27088G>C), S197N (27112G>A 27113T>C), S211A (27153T>G), S212G (27156A>G), S214N (27163G>A 27164T>C) | | | | | | | | |
| ORF6 protein (YP_009724394.1) | 1 | 62 | 100% | 300.0 | 75.0% | 62 (100%) | 42 (67.7%) | 0/0/0/0 | 0 |
| | L16I (27247C>A 27249A>T), K23R (27269A>G 27270A>G), V24I (27271G>A), S25A (27274T>G 27276C>T), Y31V (27292T>G 27293A>T 27294C>T), N34S (27302A>G 27303C>T), L35S (27304C>T 27305T>C 27306C>A), I37V (27310A>G 27312T>G), K38R (27314A>G), N39Q (27316A>C 27318T>A), S41F (27323C>T), S43P (27328T>C 27330A>T), E46K (27337G>A), N47K (27342T>G), K48N (27345A>T), Q51E (27352C>G 27354A>G), E54D (27363A>T), Q56E (27367C>G), I60L (27379A>T 27381T>A), *62Y (27387A>T) | | | | | | | | |
| ORF7a protein (YP_009724395.1) | 1 | 122 | 100% | 758.0 | 89.6% | 122 (99.2%) | 105 (85.4%) | 1/0/0/0 | 1 |
| | A8T (27415G>A), T11V (27424A>G 27425C>T), L12F (27427C>T 27429C>T), A13T (27430G>A 27432T>A), T14S (27433A>T), S36P (27499T>C 27501T>A), F59T (27568T>A 27569T>C), Q62H (27579A>C), P68A (27595C>G), V71T (27604G>A 27605T>C 27606A>T), K72R (27607A>C 27608A>G), V74T (27613G>A 27614T>C), Q94_E95insQ (27675_27676insCAA), I100L (27691A>C), I107L (27712A>C), I110L (27721A>T), T111I (27725C>T), L116I (27739C>A 27741C>T) | | | | | | | | |
| ORF7b (YP_009725296.1) | 1 | 44 | 100% | 252.0 | 79.7% | 44 (97.8%) | 36 (80.0%) | 1/0/0/0 | 1 |
| | I2N (27760T>A), S5T (27768T>A 27770A>T), L34I (27855C>A 27857G>C), H37L (27865A>T 27866T>A), N38E (27867A>G 27869T>A), T40P (27873A>C), H42T (27879C>A 27880A>C), H42_A43insK (27881_27882insAAA), A43V (27883C>T) | | | | | | | | |
| ORF8 protein (YP_009724396.1) | 1 | 120 | 99.2% | 117.0 | 15.6% | 104 (82.5%) | 37 (29.4%) | 5/17/1/1 | 0 |
| | F3L (27900T>C), V5I (27906G>A), F6V (27909T>G 27911C>T), G8_I10del (27915_27923delGGAATCATC), T12C (27927A>T 27928C>G), V13I (27930G>A 27932A>T), A14S (27933G>T), A15L (27936G>C 27937C>T), F16C (27940T>G 27941T>C), H17S (27942C>A 27943A>G 27944C>T), Q18C (27945C>T 27946A>G 27947A>C), E19I (27948G>A 27949A>T), S21T (27955G>C), L22V (27957T>G), L22_Q23insV (27959_27960insGTA), S24R (27963T>C 27964C>G 27965A>C), T26A (27969A>G 27971T>A), Q27S (27972C>T 27973A>C 27974A>T), H28N (27975C>A), Q29K (27978C>A), Y31H (27984T>C), V33L (27990G>C), D34E (27995T>A), P38_S43del (28005_28024delCCTATTCACTTCTATTCTAA), K44X (28005_28024delCCTATTCACTTCTATTCTAA), W45R (28026T>A), I47N (28033T>A 28034T>C), R48T (28036G>C 28037A>T), V49R (28038G>A 28039T>G 28040A>G), A51N (28044G>A 28045C>A), R52T (28048G>C 28049A>T), K53Y (28050A>T 28052A>T), A55T (28056G>A 28058A>T), P56A (28059C>G), L57W (28063T>G 28064A>G), I58L (28065A>C), E59_L60del (28068_28073delGAATTG), V62A (28078T>C 28079G>T), D63L (28080G>C 28081A>T 28082T>A), E64_A65del (28083_28088delGAGGCT), S67K (28092T>A 28093C>A 28094T>G), K68V (28095A>G 28096A>T 28097A>T), S69L (28099C>T), I71F (28104A>T), Q72H (28109G>T), Y73R (28110T>A 28111A>G 28112C>A), I74W (28113A>T 28114T>G 28115C>G), D75H (28116G>C 28118T>C), I76T (28120T>C 28121C>T), G77M (28122G>A 28123G>T 28124T>G), N78V (28125A>G 28126A>T), Y79Q (28128T>C 28130T>A), V81_S82del (28134_28139delGTTTCC), L84T (28143T>A 28144T>C), P85_F86insN (28148_28149insAAT), F86V (28149T>G), E92D (28169A>T), K94A (28173A>G 28174A>C 28175A>T), L95G (28176T>G 28177T>G 28178G>T), S97A (28182A>G 28183G>C 28184T>G), V99I (28188G>A), V100A (28192T>C 28193G>T), S103W (28201C>G), F104Y (28204T>A), Y105L (28206T>C 28207A>T), E106H (28209G>C 28211A>T), D107E (28214C>A), F108G (28215T>G 28216T>G), L109H (28218T>C 28219T>A 28220A>C), E110Q (28221G>C 28223G>A), E110_Y111insTAA (28223_28224insACTGCTGCA), Y111F (28225A>T), H112R (28227C>A 28228A>G 28229T>A), R115L (28237G>T), D119N (28248G>A), F120K (28251T>A 28252T>A 28253C>A) | | | | | | | | |
| nucleocapsid phosphoprotein (YP_009724397.2) | 1 | 420 | 100% | 2641.0 | 92.1% | 420 (99.3%) | 382 (90.3%) | 3/0/0/0 | 1 |
| | Q7_N8insS (28294_28295insTCA), N11S (28305A>G), S21T (28334T>A), G25D (28347G>A), S26N (28350G>A), E31G (28365A>G), S33N (28371G>A), S37P (28382T>C), D63E (28462C>A), K65R (28467A>G), S79G (28508A>G), I94V (28553A>G), D103E (28582T>G), G120S (28631G>T 28632G>C), D128E (28657C>A), I131V (28664A>G), A152N (28727G>A 28728C>A), I157T (28743T>C), N192G (28847A>G 28848A>G 28849C>T), S193N (28851G>A), T205N (28887C>A), G212S (28907G>A), N213G (28910A>G 28911A>G 28912T>A), D216E (28921T>A), A217T (28922G>A), M234V (28973A>G 28975G>T), A267Q (29072G>C 29073C>A 29074A>G), E290D (29143A>C), T334H (29273A>C 29274C>A 29275A>T), N345Q (29306A>C 29308T>A), Q349N (29318C>A 29320A>C), A376T (29399G>A), T379A (29408A>G), A381P (29414G>C 29416C>T), Q390P (29442A>C 29443A>C), L400M (29471T>A), K405R (29487A>G), Q409N (29498C>A 29500A>T), S413G (29510A>G 29511_29512insAGCTTC), S413_A414insAS (29511_29512insAGCTTC) | | | | | | | | |
| ORF10 protein (YP_009725255.1) | 1 | 39 | 100% | 212.0 | 78.8% | 39 (100%) | 32 (82.1%) | 0/0/0/0 | 2 |
| | I4V (29567A>G), F9I (29582T>A), Y14H (29597T>C), Y26* (29635C>A), I27T (29637T>C), D31G (29649A>G), V32L (29651G>T) | | | | | | | | |

*: Inserts / Deletes / Misaligned / Frameshifts

##### Alignment Detailed statistics

| | Begin | End | Coverage | Score | Concordance | Matches | Identities | I/D/M/F* | Stop Codons |
|---|---|---|---|---|---|---|---|---|---|
| **NT** | 1 | 29894 | 99.9% | 34641.0 | 58.8% | 29674 (99.0%) | 23675 (79.0%) | 77/220 | |

| | Begin | End | Coverage | Score | Concordance | Matches | Identities | I/D/M/F* | Stop Codons |
|---|---|---|---|---|---|---|---|---|---|
| **CDS** | | | | | | | | | |
| 1_orf1ab | 1 | 7097 | 100% | 44033.0 | 89.3% | 7070 (99.6%) | 6125 (86.3%) | 4/27/0/0 | 1 |
| 2_orf1a | 1 | 4406 | 100% | 25296.0 | 84.3% | 4379 (99.3%) | 3552 (80.5%) | 4/27/0/0 | 1 |
| 3_S | 1 | 1274 | 100% | 7176.0 | 82.1% | 1250 (97.7%) | 974 (76.1%) | 6/24/0/0 | 1 |
| 4_ORF3a | 1 | 276 | 100% | 1490.0 | 76.7% | 275 (99.6%) | 200 (72.5%) | 0/1/0/0 | 1 |
| 5_E | 1 | 76 | 100% | 447.0 | 94.3% | 76 (98.7%) | 73 (94.8%) | 1/0/0/0 | 1 |
| 6_M | 1 | 223 | 100% | 1419.0 | 92.9% | 222 (99.6%) | 202 (90.6%) | 0/1/0/0 | 1 |
| 7_ORF6 | 1 | 62 | 100% | 300.0 | 75.0% | 62 (100%) | 42 (67.7%) | 0/0/0/0 | 0 |
| 8_ORF7a | 1 | 122 | 100% | 758.0 | 89.6% | 122 (99.2%) | 105 (85.4%) | 1/0/0/0 | 1 |
| 9_ORF7b | 1 | 44 | 100% | 252.0 | 79.7% | 44 (97.8%) | 36 (80.0%) | 1/0/0/0 | 1 |
| 10_ORF8 | 1 | 120 | 99.2% | 117.0 | 15.6% | 104 (82.5%) | 37 (29.4%) | 5/17/1/1 | 0 |
| 11_N | 1 | 420 | 100% | 2641.0 | 92.1% | 420 (99.3%) | 382 (90.3%) | 3/0/0/0 | 1 |
| 12_ORF9b | 1 | 98 | 100% | 451.0 | 73.0% | 98 (99.0%) | 72 (72.7%) | 1/0/0/0 | 1 |
| 13_ORF10 | 1 | 39 | 100% | 212.0 | 78.8% | 39 (100%) | 32 (82.1%) | 0/0/0/0 | 2 |
| 14_ORF14 | 1 | 74 | 100% | 388.0 | 77.3% | 74 (100%) | 57 (77.0%) | 0/0/0/0 | 2 |
| **Proteins** | | | | | | | | | |
| orf1ab polyprotein (YP_009724389.1) | 1 | 7097 | 100% | 44033.0 | 89.3% | 7070 (99.6%) | 6125 (86.3%) | 4/27/0/0 | 1 |

| Begin | End | Coverage | Score | Concordance | Matches | Identities | I/D/M/F* | Stop Codons |
|---|---|---|---|---|---|---|---|---|

P6L (282C>T), F8V (287T>G), V38A (378T>C), Q44E (395C>G), D48N (407G>A), V56L (431G>C 433T>G), R77L (494C>T 495G>T), T78S (498C>G 499T>C), A79T (500G>A 502A>C), P80N (503C>A 504C>A), V84K (515G>A 516T>A 517T>G), M85V (518A>G 520G>C), L92M (539C>A 541C>G), E93D (544A>C), E102I (569G>A 570A>T 571G>A), I114T (606T>C 607A>C), V116I (611G>A 613G>T), K120N (625G>T), A138I (677G>A 678C>T), F143Y (693T>A), Y154I (725T>A 726A>T), F157Y (735T>A), Q158E (737C>G), E159Q (740G>C), S166G (761A>G), V169A (771T>C 772T>A), T170L (773A>C 774C>T), M174T (786T>C 787G>T), Y182V (809T>G 810A>T), E199D (862G>T), L204F (875C>T), A211S (896G>T 898T>A), S212M (899T>A 900C>T 901A>G), F221Y (927T>A 928T>C), D223E (934C>G), T224S (935A>T 937T>G), E233D (964A>C), Y241F (987A>T), E246D (1003A>T), L251H (1016T>C 1017T>A 1018G>C), L259S (1040T>A 1041T>G 1042G>T), N267K (1066T>A), N272K (1081T>G), I280K (1104T>A), I281V (1106A>G), T283V (1112A>G 1113C>T 1114T>C), L293T (1142C>A 1143T>C), D294E (1147T>G), N310Q (1193A>C 1195T>G), Q314N (1205C>A 1207A>T), C316H (1211T>C 1212G>A), D324N (1235G>A), G327D (1245G>A), T329V (1250A>G 1251C>T), G334C (1265G>T), V337L (1274G>C 1276T>G), F343H (1292T>C 1293T>A), T350V (1313A>G 1314C>T), K351I (1317A>T 1318A>T), A354P (1325G>C 1327C>T), Q362T (1349C>A 1350A>C 1351A>T), I368M (1369T>G), Y369P (1370T>C 1371A>C 1372T>A), H374Q (1387C>A), N375D (1388A>G 1390T>C), S376P (1391T>C), V378I (1397G>A 1399A>T), L384V (1415C>G), E386D (1423A>T), E390H (1433G>C 1435A>C), G392N (1439G>A 1440G>A), L393I (1442T>A 1444G>T), K394E (1445A>G), I396R (1451A>C 1452T>G 1453T>A), I404R (1476T>G 1477T>A), A405C (1478G>T 1479C>G 1480C>T), S412A (1499T>G 1501T>C), H417Y (1514C>T), C420R (1523T>C), N430D (1553A>G 1555C>T), C433S (1563G>C 1564T>A), N434G (1565A>G 1566A>G), V438I (1577G>A), V439T (1580G>A 1581T>C), E441D (1588A>C), G442N (1589G>A 1590G>A), S443V (1592T>G 1593C>T 1594C>G), G445T (1598G>A 1599G>C 1600T>C), D448E (1609C>G), N449D (1610A>G 1612C>T), Q455S (1628C>A 1629A>G 1630A>T), K456R (1631A>C 1632A>G 1633A>T), K458R (1637A>C 1638A>G 1639A>T), K468H (1667A>C 1669A>T), I473V (1682A>G 1684C>T), V488I (1727G>A 1729G>T), E489D (1732A>C), V491I (1736G>A 1738G>A), G493S (1742G>A), A498S (1757G>T 1759A>T), Q501T (1766C>A 1767A>C 1768A>C), F509Y (1791T>A), A516P (1811G>C 1813T>C), K517V (1814A>G 1815A>T), E525Q (1838G>C), K527R (1845A>G), I529V (1850A>G 1852A>T), S531T (1857G>C 1858T>A), Y534C (1866A>G), A535G (1869C>G 1870A>T), A537P (1874G>C 1876A>C), E539Q (1880G>C), R542G (1889C>G), V544I (1895G>A 1897A>C), S549A (1910T>G 1912C>G), E553D (1924A>T), T554A (1925A>G 1927T>A), Q556N (1931C>A 1933A>C), N557H (1934A>C 1936T>C), V559I (1940G>A 1942G>T), R560P (1944G>C), V561D (1947T>A), K564R (1956A>G 1957G>A), I567V (1964A>G 1966A>C), Q575E (1988C>G 1990G>A), Y576Q (1991T>C 1993T>G), I581V (2006A>G 2008T>C), M585V (2018A>G 2020G>T), F586Y (2022T>A 2023C>T), A591L (2036G>C 2037C>T 2038T>C), N594S (2046A>G), L595V (2048C>G 2050A>C), V596I (2051G>A), V597I (2054G>A 2056A>T), I601V (2066A>G 2068T>A), V605L (2078G>C), L608Q (2087T>C 2088T>A), T614S (2105A>T), I616L (2111A>C 2113C>T), F617L (2116T>G), V620T (2123G>A 2124T>C), Y621V (2126T>G 2127A>T), K625R (2139A>G 2140A>G), V627I (2144G>A), L628F (2147C>T), D629E (2152T>A), L631I (2156C>A), E633A (2163A>C), F635L (2168T>C), K636S (2172A>G 2173G>T), E637A (2175A>C), R643K (2193G>A 2194A>G), G645A (2199G>C), V649L (2210G>C 2212T>C), I652L (2219A>C), S653I (2222T>A 2223C>T 2224A>T), C655G (2228T>G), A656V (2232C>T), C657F (2235G>T), E658D (2239A>C), G661K (2246G>A 2247G>A 2248T>G), V665Q (2258G>C 2259T>A 2260C>G), T666V (2261A>G 2262C>T 2263C>T), C667A (2264T>G 2265G>C), A668S (2267G>T), K669D (2270A>G 2272G>T), E670N (2273G>A 2275A>C), E673D (2284G>T), S674C (2285A>T), Q676K (2291C>A 2293G>A), T677C (2294A>T 2295C>G 2296A>C), F679I (2300T>A), K680D (2303A>G 2305G>T), L681V (2306C>G), F685A (2318T>G 2319T>C 2320T>A), A687E (2325C>A 2326T>A), L688M (2327T>A), A690I (2333G>A 2334C>T), S692Q (2339T>C 2340C>A 2341T>A), I693V (2342A>G), I694T (2346T>C), G696A (2352G>C), K701R (2366A>C 2367A>G), A702S (2369G>T 2371C>A), T708V (2387A>G 2388C>T 2389A>C), V710I (2393G>A), T711A (2396A>G 2398G>T), H712Q (2401C>A), K719Q (2420A>C), V721I (2426G>A 2428T>A), K722R (2429A>C 2430A>G 2431A>T), S723G (2432T>G 2433C>G), R724K (2436G>A 2437A>G), E726Q (2441G>C 2443A>G), T727L (2444A>C 2445C>T 2446T>G), G728Q (2447G>C 2448G>A 2449C>A), I739V (2480A>G 2482T>A), I740T (2484T>C), E745D (2500A>T), T746S (2501A>T), L747H (2505T>A), P748D (2507C>G 2508C>A), E750V (2514A>T), V751L (2516G>C 2518G>T), L752T (2519T>A 2520T>C 2521A>C), T753S (2522A>T 2524A>T), T760N (2544C>A 2545T>C), D762E (2551T>A), Q764E (2555C>G), P765A (2558C>G), Q768T (2567C>A 2568A>C 2569A>G), T770V (2573A>G 2574C>T), S771D (2576A>G 2577G>A), E772S (2579G>A 2580A>G 2581A>C), A773F (2582G>T 2583C>T 2584T>C), V774T (2585G>A 2586T>C 2587T>A), E775N (2588G>A 2590A>T), A776G (2592C>G 2593T>A), P777A (2594C>G 2596A>T), L778I (2597T>A 2599G>C), I785V (2618A>G 2620T>A), T796K (2652C>A), K798Q (2657A>C 2659G>A), A803S (2672G>T 2674A>T), N805G (2678A>G 2679A>G), M806L (2681A>T 2683G>A), M807L (2684A>C), V808A (2688T>C 2689A>T), T812V (2699A>G 2700C>T), T814R (2705A>C 2706C>G 2707A>C), T821I (2727C>T 2728A>T), K822_V823insG (2731_2732insGGT), D827E (2746T>A), I831W (2756A>T 2757T>G 2758A>G), S838N (2778G>A), N840R (2784A>G 2785T>A), I849V (2810A>G), A859V (2841C>T), L864S (2855C>T 2856T>C), N869T (2871A>C), D877E (2896T>G), I880V (2903A>G 2905A>G), E888D (2929A>T), P892N (2939C>A 2940C>A 2941A>C), L893M (2942C>A), M902V (2969A>G 2971G>A), Y905F (2979A>T), E910D (2995G>T), S911A (2996T>G), F914E (3005T>G 3006T>A 3007T>A), K915N (3010A>C), L916F (3013G>T), A917S (3014G>T 3016T>A), H919R (3021A>G), D930E (3055T>A), E933D (3064A>C), G934D (3066G>A), G934_D935insA (3067_3068insGCA), D935E (3070T>G), F941I (3086T>A), E942D (3091G>T), P943E (3092C>G 3093C>A), S944T (3095T>A 3097A>C), T945C (3098A>T 3099C>G), Q946E (3101C>G), Y947H (3104T>C), K958L (3137A>C 3138A>T 3139A>C), T965S (3158A>T 3160T>A), S966A (3161T>G), A967E (3165C>A 3166T>A), A968T (3167G>A 3169T>A), L969V (3170C>G), Q970R (3174A>G), P971V (3176C>G 3177C>T), Q975E (3188C>G), D983_T999del (3212_3262delGATAGTCAACAAACTGTTGGTCAACAAGACGGCAGTGAGGACAATCAGACA), I1002E (3269A>G 3270T>A 3271T>G), T1004S (3275A>T), I1005_V1006del (3278_3283delATTGTT), V1008I (3287G>A), Q1009E (3290C>G 3292A>G), Q1011E (3296C>G), L1012P (3299T>C 3300T>C), E1013_M1014del (3302_3307delGAGATG), L1016P (3312T>C), V1019E (3321T>A 3322T>A), V1020E (3324T>A 3325T>A), Q1021_I1023del (3327_3335delAGACTATTG), E1024P (3327_3335delAGACTATTG 3336A>C), S1027Q (3344A>C 3345G>A 3346T>G), S1029T (3351G>C), Y1039A (3380T>G 3381A>C), N1042C (3389A>T 3390A>G), A1043V (3393C>T 3394A>T), E1047K (3404G>A 3406A>G), K1050Q (3413A>C), K1051S (3417A>G 3418G>T), V1052A (3420T>C 3421A>T), K1053N (3424A>T), T1055M (3429C>T 3430A>G), V1057I (3434G>A), V1063I (3452G>A 3454T>A), Y1064H (3455T>C), N1081G (3506A>G 3507A>G), V1085K (3518G>A 3519T>A 3520T>G), A1092K (3539G>A 3540C>A 3541T>G), T1093L (3542A>C 3543C>T 3544T>A), K1098T (3558A>C), V1104L (3575G>T 3577T>G), H1113K (3602C>A 3604C>G), V1122L (3629G>C 3631T>A), K1124A (3635A>G 3636A>C), S1133A (3662A>G 3663G>C 3664T>A), Q1140S (3683C>T 3684A>C 3685G>A), H1141Q (3688C>G), E1142D (3691A>C), V1143I (3692G>A 3694T>C), D1157K (3734G>A 3736C>A), I1159L (3740A>C 3742A>T), H1160Q (3745T>G), R1163Q (3752A>C 3753G>A), D1167Q (3764G>C 3766T>G), N1172Q (3779A>C 3781T>G), L1175I (3788T>A 3790A>T), F1178N (3797T>A 3798T>A), N1181A (3806A>G 3807A>C), D1184E (3817C>G), K1185Q (3818A>C 3820A>G), L1186V (3821C>G), S1188M (3827T>A 3828C>T 3829A>G), S1189D (3830A>G 3831G>A 3832C>T), F1190Y (3834T>A), E1192D (3841A>T), M1193N (3843T>A 3844G>C), K1194L (3845A>C 3846A>T), S1195K (3849G>A 3850T>G), E1196P (3851G>C 3852A>C 3853A>T), K1197R (3855A>G 3856G>A), Q1198V (3857C>G 3858A>T 3859A>G), V1199E (3861T>A 3862T>A), E1200A (3864A>C), Q1201P (3867A>C 3868A>T), I1203Q (3872A>C 3873T>A 3874C>A), A1204E (3876C>A 3877T>G), I1206P (3881A>C 3882T>C 3883T>A), K1208N (3889A>C), E1209T (3890G>A 3891A>C 3892G>A), V1211D (3897T>A), K1212_P1213del (3899_3904delAAGCCA), F1214S (3906T>C 3907T>C), I1215K (3909T>A), S1218E (3917A>G 3918G>A 3919T>G), P1220del (3923_3925delCCT), E1223V (3933A>T), R1225K (3939G>A 3940A>G), K1226P (3941A>C 3942A>C 3943A>T), Q1227V (3944C>G 3945A>T 3946A>C), D1229V (3951A>T 3952T>G), K1230_K1231insP (3955_3956insCCA), V1236I (3971G>A), E1237D (3976A>T), E1251N (4016G>A 4018A>T), N1252K (4021C>G), Y1256F (4032A>T), I1257A (4034A>G 4035T>C), N1262K (4051T>G), H1264Y (4055C>T 4057T>C), P1265H (4059C>A 4060A>T), A1268Q (4067G>C 4068C>A 4069C>G), T1269N (4071C>A 4072T>C), L1270M (4073C>A 4075T>G), V1271L (4076G>C), S1272R (4081T>A), D1273G (4083A>G 4084C>T), I1274E (4085A>G 4086T>A 4087T>A), I1276M (4093C>G), T1277S (4094A>T), K1280E (4103A>G), I1286M (4123A>G), V1291I (4136G>A 4138T>C), Q1292T (4139C>A 4140A>C 4141A>T), E1293S (4142G>A 4143A>G 4144G>T), V1295D (4149T>A), L1296I (4151T>A 4153A>C), A1298C (4157G>T 4158C>G), T1303S (4172A>T 4174T>C), A1314S (4205G>T 4207G>A), K1315R (4209A>G), R1318K (4218G>A 4219A>G), T1322V (4229A>G 4230C>T 4231A>T), N1324E (4235A>G 4237T>G), L1334C (4266T>G 4267A>T), N1335A (4268A>G 4269A>C), V1339L (4280G>C 4282A>T), V1345A (4299T>C 4300G>T), I1355V (4328A>G 4330T>A), I1359E (4340A>G 4341T>A 4342T>A), I1360A (4343A>G 4344T>C 4345C>A), S1361P (4346T>C), E1363A (4353A>C 4354G>T), Q1365E (4358C>G), V1391I (4436G>A 4438C>A), V1393M (4442G>A), E1394D (4447A>T), T1395V (4448A>G 4449C>T), K1396R (4452A>G), V1399M (4460G>A 4462T>G), S1400A (4463T>G), V1415I (4508G>A 4510G>C), A1420V (4524C>T 4525T>C), Y1423F (4533A>T), T1429E (4550A>G 4551C>A 4552A>G), T1430P (4553A>C), L1434I (4565C>A), N1436T (4572A>C 4573C>G), T1437K (4575C>A 4576A>G), D1440S (4583G>T 4584A>C), T1444P (4595A>C 4597T>G), L1450I (4613C>A), L1457F (4636A>T), Y1465C (4659A>G), V1471A (4677T>C 4678G>T), T1474V (4685A>G 4686C>T), A1485T (4718G>A 4720G>A), P1497S (4754C>T), I1502V (4769A>G 4771T>A), I1505V (4778A>G 4780C>T), K1512R (4800A>G), S1520R (4823T>C 4824C>G), Q1522E (4829C>G 4831A>G), I1525V (4838A>G 4840A>T), S1534I (4866G>T), Y1537H (4874T>C), T1538_S1539insL (4879_4880insCTG), S1539E (4880A>G 4881G>A 4882T>G), N1540S (4884A>G 4885T>C), T1542V (4889A>G 4890C>T), T1543E (4892A>G 4893C>A 4894A>G), I1551L (4916A>C 4918C>T), T1552S (4919A>T 4921C>A), F1553L (4922T>C), N1555K (4930T>A), T1558S (4938C>G 4939A>T), R1566K (4962G>A), I1577T (4995T>C), V1583L (5012G>C), N1611V (5096A>G 5097A>T 5098T>A), S1612N (5099T>A 5100C>A 5101A>T), Y1619F (5121A>T), N1623S (5133A>G), V1629S (5150G>A 5151T>G), T1638L (5177A>C 5178C>T), P1640E (5183C>G 5184C>A 5185T>G), Y1658F (5238A>T 5239C>T), N1662G (5249A>G 5250A>G), A1677S (5294G>T 5296C>T), T1678S (5298C>G), A1679V (5301C>T 5302A>T), T1682A (5309A>G), I1686L (5321A>C 5323A>T), L1688V (5327T>G 5329G>C), P1692A (5339C>G), D1697E (5356T>G), E1706D (5383A>T), C1718S (5417T>A), S1733T (5463G>C 5464T>C), Y1734H (5465T>C 5467C>T), F1736L (5471T>C 5473T>A), D1742E (5491T>A), C1744A (5495T>G 5496G>C 5497C>A), T1754H (5525A>C 5526C>A), Q1758K (5537C>A 5539G>A), Q1759T (5540C>A 5541A>C 5542G>T), K1763T (5553A>C), E1777D (5596A>T), Q1778N (5597C>A 5599A>T), F1779L (5600T>C), K1781T (5607A>C), Q1784S (5615C>T 5616A>C 5617G>C), T1788V (5627A>G 5628C>T), K1791R (5636A>C 5637A>G 5638A>T), Q1792D (5639C>G 5641A>T), K1795Q (5648A>C), P1803S (5672C>T), Q1813E (5702C>G), E1815K (5708G>A), K1817Q (5714A>C), H1818Q (5719T>A), T1822L (5729A>T 5730C>T 5731T>A), S1825N (5739G>A), K1837T (5775A>C 5776A>T), S1841A (5786T>G), C1847R (5804T>C 5806C>T), L1852H (5819T>C 5820T>A 5821A>C), S1856M (5831T>A 5832C>T 5833C>G), I1863V (5852A>G 5854T>G), N1871T (5877A>C 5878C>A), T1881S (5906A>T 5908T>G), V1888T (5927G>A 5928T>C), C1889Y (5931G>A 5932T>C), D1893E (5944C>A), N1898G (5957A>G 5958A>G 5959T>G), S1905A (5978T>G), F1907Y (5985T>A 5986C>T), N1917T (6015A>C 6016C>T), Y1920L (6024A>T 6025T>A), F1930L (6053T>C 6055T>C), V1931T (6056G>A 6057T>C), D1933S (6062G>T 6063A>C), I1935T (6069T>C 6070C>A), L1944M (6095T>A 6097A>G), Y1947F (6105A>T 6106T>C), K1948T (6108A>C 6109G>A), K1956S (6131A>T 6132A>C 6133A>T), K1973R (6183A>G), T1976S (6191A>T), P1977A (6194C>G 6196C>G), V1994I (6245G>A), N1996Q (6251A>C 6253T>G), N1999T (6261A>C 6262T>C), A2001T (6266G>A 6268C>A), Y2003F (6273A>T 6274T>C), I2010L (6293A>T), E2020D (6325A>T), D2026E (6343T>A), K2029A (6350A>G 6351A>C 6352G>A), S2030V (6353T>G 6354C>T), A2033T (6362G>A 6364G>A), D2043S (6392G>A 6393A>G), L2044Q (6396T>A), K2045Q (6398A>C), V2047T (6404G>A 6405T>C), D2060E (6445C>A), L2062I (6449C>A 6451T>A), N2065D (6458A>G 6460T>C), D2074N (6485G>A 6487C>T), I2075V (6488A>G 6490T>C), A2080S (6503G>T), N2081D (6506A>G), N2082E (6509A>G 6511T>A), S2083G (6512A>G), L2084V (6515T>G 6517A>T), I2086V (6521A>G 6523T>A), E2088Q (6527G>C), V2090L (6533G>T 6535T>A), T2093E (6542A>G 6543C>A 6544A>G), D2101E (6568C>A), S2103T (6572T>A 6574T>A), L2105I (6578C>A), R2115L (6608A>C 6609G>T), V2116A (6612T>C 6613A>C), L2122I (6629C>A), L2127I (6644T>A 6646A>T), V2130I (6653G>A), D2136S (6671G>A 6672A>G), T2137K (6675C>A 6676T>A), A2139L (6680G>T 6681C>T 6682T>G), N2140A (6683A>G 6684A>C), A2142V (6690C>T 6691T>C), N2147G (6704A>G 6705A>G 6706C>A), K2148Q (6707A>C), V2149A (6711T>C 6712T>A), V2150A (6714T>C 6715T>A), S2151I (6717G>T), T2154S (6725A>T 6727T>A), I2156C (6731A>T 6732T>G 6733A>C), V2157A (6735T>C), T2158K (6738C>A 6739A>G), C2160L (6744G>T 6745T>A), L2161A (6746T>G 6747T>C), N2162Q (6749A>C 6751C>A), C2165F (6759G>T), T2166N (6762C>A 6763T>C), F2172V (6779T>G 6781C>G), L2177F (6794C>T 6796A>C), R2184K (6816G>A), K2191R (6837A>G), M2194L (6845A>C 6847G>A), T2202S (6870C>G), G2207A (6885G>C), F2209L (6892T>A), E2212D (6901G>T), S2214G (6905T>G 6906C>G 6907A>C), F2215I (6908T>A), L2218V (6917T>G), N2222K (6931T>A), I2227F (6944A>T 6946A>C), N2228T (6948A>C 6949T>A), I2230A (6953A>G 6954T>C 6955A>T), I2231M (6958T>G), F2233L (6962T>C 6964T>A), V2238I (6977G>A), Y2245C (6999A>G 7000C>T), S2246V (7001T>G 7002C>T), L2250F (7015A>T), M2254L (7025A>T 7027G>A), L2257F (7036A>T), M2259A (7040A>G 7041T>C 7042G>T), T2264N (7056C>A), Y2266V (7061T>G 7062A>T 7063C>T), G2269L (7070G>T 7071G>T 7072C>G), T2274S (7085A>T), I2278T (7098T>C), A2279M (7100G>A 7101C>T 7102A>G), T2280D (7103A>G 7104C>A 7105C>T), Y2281F (7107A>T), T2283E

| | | | | | | | | | |
|---|---|---|---|---|---|---|---|---|---|
| | (7112A>G 7113C>A 7114T>A), I2286F (7121A>T 7123A>T), V2290I (7133G>A), T2300S (7163A>T 7165C>T), S2303A (7172T>G), I2309V (7190A>G 7192T>G), F2314Y (7206T>A 7207T>C), W2316L (7211T>C 7212G>T 7213G>A), A2320I (7223G>A 7224C>T), F2321L (7228T>A), V2324A (7236T>C 7237T>C), F2328V (7247T>G), I2332M (7261T>G), R2336K (7272G>A 7273G>A), V2340L (7283G>T), A2344S (7295G>T 7297T>A), L2349V (7310T>G), S2352G (7319A>G), V2356S (7331G>A 7332T>G 7333A>T), L2367F (7366A>T), N2370S (7374A>G), L2371I (7376C>A), I2377V (7394A>G), V2393I (7442G>A), V2400I (7463G>A 7465T>C), V2401M (7466G>A 7468A>G), N2405T (7479A>C 7480T>C), V2430M (7553G>A 7555T>G), R2431K (7557G>A 7558A>G), K2442R (7589A>C 7590A>G 7591A>T), L2447T (7604C>A 7605T>C 7606A>T), V2453L (7622G>C 7624T>C), A2460T (7643G>A), T2495A (7748A>G 7750A>T), S2500A (7763T>G 7765C>G), I2501L (7766A>C 7768C>T), S2517P (7814T>C 7816T>G), E2550D (7915A>C), S2553A (7922T>G), A2554S (7925G>T 7927A>T), A2584T (8015G>A 8017G>T), A2587S (8024G>T 8026A>C), N2596D (8051A>G 8053T>C), S2600A (8063T>G), N2603S (8073A>G 8074C>T), T2611A (8096A>G), E2617H (8114G>C 8116A>C), A2618S (8117G>A 8118C>G 8119T>C), N2623G (8132A>G 8133A>G), S2625A (8138T>G 8140C>T), N2628G (8147A>G 8148A>G), I2634V (8165A>G 8167T>G), F2641V (8186T>G), S2644T (8195T>A 8197A>C), E2647D (8206A>C), V2652I (8219G>A), Q2660H (8245A>C), I2663L (8252A>T), Y2673F (8283A>T 8284T>C), S2695N (8349G>A), I2709V (8390A>G), A2710S (8393G>T 8395T>A), F2718Y (8418T>A), L2738I (8477T>A), K2741R (8487A>G 8488G>A), V2754I (8525G>A), A2759S (8540G>T), N2767S (8565A>G), N2768T (8568A>C), W2769C (8572G>T), L2770F (8575G>T), Q2772L (8580A>T 8581G>T), L2773M (8582T>A 8584A>G), I2774L (8585A>C), V2776A (8592T>C 8593T>C), V2779L (8600G>T), F2780C (8604T>G), L2781V (8606C>G), F2782L (8609T>C), V2783A (8613T>C), A2785L (8618G>T 8619C>T 8620T>G), I2786V (8621A>G), F2787C (8625T>G 8626C>T), L2789I (8630T>A 8632A>C), I2790V (8633A>G 8635A>T), T2791M (8637C>T 8638A>G), V2795T (8648G>A 8649T>C 8650C>A), M2796L (8651A>T), K2798I (8658A>T 8659A>C), T2800D (8663A>G 8664C>A), D2801G (8667A>G 8668C>T), F2802Y (8670T>A 8671T>C), S2803T (8672T>A), S2804N (8676G>A), D2813Q (8702G>C 8704T>G), G2814D (8706G>A), A2821I (8726G>A 8727C>T 8728A>T), T2825D (8738A>G 8739C>A), D2833G (8763A>G), T2836A (8771A>G), T2846K (8802C>A 8803T>A), A2850S (8813G>A 8814C>G 8815T>C), L2853V (8822T>G 8824G>A), I2854V (8825A>G 8827T>A), V2857I (8834G>A), V2862I (8849G>A 8851G>T), V2865I (8858G>A 8860C>A), I2873V (8882A>G 8884A>G), T2876A (8891A>G), T2877I (8895C>T 8896T>C), T2906S (8982C>G), S2926M (9041T>A 9042C>T 9043T>G), V2938L (9077G>T 9079A>G), V2943I (9092G>A), A2944S (9095G>T), E2946S (9101G>A 9102A>G 9103A>T), S2947E (9104A>G 9105G>A 9106T>G), S2981A (9206T>G), A2994V (9246C>T 9247T>A), V2996I (9251G>A), V2998L (9257G>C), D3009E (9292T>G), Y3010H (9293T>C), S3013A (9302T>G), P3015S (9308C>T), V3024M (9335G>A 9337A>G), L3027I (9344C>A 9346T>A), T3028A (9347A>G), M3030I (9355G>C), I3035V (9368A>G 9370T>G), I3038V (9377A>G 9379T>G), I3043V (9392A>G 9394A>G), I3047V (9404A>G), V3053I (9422G>A 9424A>T), V3056L (9431G>T 9433A>G), L3060A (9443C>G 9444T>C), R3066K (9462G>A 9463G>A), A3070V (9474C>T), S3075N (9489G>A 9490T>C), F3080A (9503T>G 9504T>C), T3082A (9509A>G 9511T>A), V3091I (9536G>A), T3095V (9548A>G 9549C>T), V3097A (9555T>C), I3108F (9587A>T), L3116F (9611C>T 9613T>C), I3126L (9641A>C), M3129F (9650A>T 9652G>T), V3130A (9654T>C 9655T>C), T3133S (9662A>T 9664A>T), L3135I (9668T>A 9670A>T), I3142A (9689A>G 9690T>C 9691T>A), A3143I (9692G>A 9693C>T 9694T>C), I3145V (9698A>G 9700C>A), I3146F (9701A>T 9703T>C), T3150L (9713A>C 9714C>T 9715A>G), F3153C (9723T>G), Y3154H (9725T>C), S3158N (9738G>A 9739T>C), K3162R (9750A>G), R3163K (9753G>A), V3166M (9761G>A 9763C>G), S3171T (9776T>A 9778C>A), D3196E (9853T>G), V3197T (9854G>A 9855T>C 9856G>A), M3221L (9926A>T 9928G>A), S3246A (10001T>G), V3298T (10157G>A 10158T>C), S3309A (10190T>G 10192T>A), N3328S (10248A>G 10249T>C), V3349L (10310G>C 10312A>G), K3351R (10317A>G), A3357S (10334G>T 10336C>T), F3397H (10454T>C 10455T>A 10456C>T), N3443K (10594C>A), V3465L (10658G>T 10660T>A), S3530A (10853T>G 10855A>T), A3548T (10907G>A), L3549I (10910T>A 10912A>T), S3570G (10973A>G), A3571K (10976G>A 10977C>A 10978A>G), V3572F (10979G>T 10981G>C), R3574K (10986G>A), T3575I (10989C>T 10990A>T), I3576V (10991A>G 10993C>T), L3583M (11012T>A), I3587F (11024A>T 11026T>C), V3593I (11042G>A), L3606V (11081T>G 11083G>T), A3615T (11108G>A), M3616L (11111A>C 11113G>T), I3619M (11122T>G), M3621I (11128G>T), S3622A (11129T>G), F3624C (11136T>G), M3627L (11144A>C), F3628L (11147T>C), D3668E (11269T>A), M3669L (11270A>T), V3670A (11274T>C), F3677Y (11295T>A), K3678R (11298A>G), V3689L (11330G>T), G3704A (11376G>C), L3715I (11408T>A 11410G>T), I3737V (11474A>G 11476A>T), V3751I (11516G>A), G3757A (11535G>C), M3761V (11546A>G), C3766Y (11562G>A), I3768L (11567A>T 11569T>G), F3769L (11572C>A), F3789C (11631T>G), T3791C (11636A>T 11637C>G 11638T>C), N3833S (11763A>G), V3847I (11804G>A), K3929R (12051A>G), F3957Y (12135T>A), I4074V (12485A>G 12487A>C), N4078G (12497A>G 12498A>G 12499C>T), T4087N (12525C>A 12526A>C), S4115N (12609G>A 12610T>C), T4174N (12786C>A 12787A>T), T4175S (12788A>T 12790A>G), L4188H (12827T>C 12828T>A 12829A>C), A4276P (13091G>C), K4366R (13362A>G), M4390L (13433A>T), L4391M (13436C>A 13438T>G), Q4397S (13454C>T 13455A>C), S4398T (13457T>A), D4432E (13560T>A), D4454E (13626T>A), D4455G (13628A>G), I4458L (13636A>T 13638T>A), F4469M (13669T>A 13671C>G), L4482V (13708C>G), K4490V (13732A>G 13733A>T 13734A>C), I4498V (13756A>G), A4577S (13993G>T 13995T>A), N4590D (14032A>G), I4615V (14107A>G), T4617V (14113A>G 14114C>T 14115C>A), T4618A (14116A>G 14118G>A), S4621C (14125A>T 14127T>C), V4625I (14137G>A), T4644A (14194A>G), V4649M (14209G>A 14211T>G), T4651A (14215A>G), T4654A (14224A>G), Y4657L (14233T>C 14234A>T 14235C>T), K4673C (14281A>T 14282A>G 14283A>T), V4691I (14335G>A), N5003T (15272A>C 15273C>T), T5035N (15368C>A 15369A>C), S5039N (15380G>A), T5131H (15655A>C 15656C>A 15657A>T), D5132E (15660C>A), N5135D (15667A>G), F5158Y (15737T>A 15738C>T), T5161N (15746C>A 15747T>C), S5164A (15754T>G), S5176A (15790T>G), V5894I (17944G>A 17946A>T), V5939I (18079G>A 18081A>C), T5956I (18131C>T 18132T>A), E6024D (18336A>T), P6053E (18421C>G 18422C>A 18423T>A), D6057E (18435T>A), S6059T (18439T>A), S6062N (18449G>A), N6101G (18565A>G 18566A>G 18567T>A), R6137K (18674G>A), A6145S (18697G>T 18699T>A), H6153N (18721C>A), I6156V (18730A>G 18732T>G), L6184Q (18815T>A 18816G>A), Y6185H (18817T>C), T6218S (18916A>T), I6219V (18919A>G), K6229R (18950A>G), I6230V (18952A>G), A6232S (18958G>T 18960G>T), A6244S (18994G>T), D6270E (19074T>A), S6299H (19159T>C 19160C>A 19161T>C), S6321A (19225T>G 19227C>A), V6362T (19348G>A 19349T>C), L6418Q (19516T>C 19517T>A), V6435I (19567G>A), F6459Y (19640T>A), Q6470H (19674A>C), Q6471A (19675C>G 19676A>C 19677G>C), V6474A (19685T>C), T6482A (19708A>G), V6490I (19732G>A), L6494I (19744T>A 19746G>C), V6521I (19825G>A 19827G>T), D6543E (19893T>A), I6548V (19906A>G), S6555T (19927T>A 19929T>A), T6566S (19961C>G 19962G>T), I6567A (19963A>G 19964T>C), A6569S (19969G>T 19971A>T), P6570S (19972C>T), F6574L (19986T>G), D6580E (20004T>A), Q6604T (20074C>A 20075A>C), V6607K (20083G>A 20084T>A 20085A>G), K6610A (20092A>G 20093A>C), L6614V (20104C>G 20106T>C), A6623S (20131G>T 20133C>A), Y6631F (20156A>T), V6637I (20173G>A), V6638I (20176G>A 20178C>T), N6651D (20215A>G 20217T>C), Q6653E (20221C>G 20223A>G), E6654D (20226A>T), I6663T (20252T>C), E6675Q (20287G>C 20289A>G), S6695G (20347A>G 20349T>A), L6703M (20371C>A), F6710S (20393T>C 20394T>A), K6711Q (20395A>C 20397G>A), E6712D (20400A>T), F6715L (20407T>C), E6716K (20410G>A), V6766I (20560G>A), T6777A (20593A>G 20595A>T), S6799A (20659T>G 20661T>A), D6830E (20754T>A), S6831N (20756G>A), T6833V (20761A>G 20762C>T 20763A>T), L6834I (20764T>A), K6933R (21062A>G), N6936H (21070A>C), I6951L (21115A>C 21117T>G), Q6956K (21130C>A 21132A>G), V6965I (21157G>A 21159G>A), I6967V (21163A>G), A6986S (21220G>T), C7007A (21283T>G 21284G>C), R7014K (21304C>A 21305G>A 21306C>G), V7021T (21325G>A 21326T>C), G7063N (21451G>A 21452G>A), L7070Y (21473T>A 21474A>T), S7074E (21484A>G 21485G>A 21486T>A), I7088V (21526A>G), V7092I (21538G>A) | | | | | | | | |
| leader protein (YP_009725297.1) | 1 | 180 | 100% | 1059.0 | 86.0% | 180 (100%) | 152 (84.4%) | 0/0/0/0 | 0 |
| | P6L (282C>T), F8V (287T>G), V38A (378T>C), Q44E (395C>G), D48N (407G>A), V56L (431G>C 433T>G), R77L (494C>T 495G>T), T78S (498C>G 499T>C), A79T (500G>A 502A>C), P80N (503C>A 504C>A), V84K (515G>A 516T>A 517T>G), M85V (518A>G 520G>C), L92M (539C>A 541C>G), E93D (544A>C), E102I (569G>A 570A>T 571G>A), I114T (606T>C 607A>C), V116I (611G>A 613G>T), K120N (625G>T), A138I (677G>A 678C>T), F143Y (693T>A), Y154I (725T>A 726A>T), F157Y (735T>A), Q158E (737C>G), E159Q (740G>C), S166G (761A>G), V169A (771T>C 772T>A), T170L (773A>C 774C>T), M174T (786T>C 787G>T) | | | | | | | | |
| nsp2 (YP_009725298.1) | 1 | 638 | 100% | 3197.0 | 72.6% | 638 (100%) | 436 (68.3%) | 0/0/0/0 | 0 |
| | Y2V (809T>G 810A>T), E19D (862G>T), L24F (875C>T), A31S (896G>T 898T>A), S32M (899T>A 900C>T 901A>G), F41Y (927T>A 928T>C), D43E (934C>G), T44S (935A>T 937T>G), E53D (964A>C), Y61F (987A>T), E66D (1003A>T), L71H (1016T>C 1017T>A 1018G>C), L79S (1040T>A 1041T>G 1042G>T), N87K (1066T>A), N92K (1081T>G), I100K (1104T>A), I101V (1106A>G), T103V (1112A>G 1113C>T 1114T>C), L113T (1142C>A 1143T>C), D114E (1147T>G), N130Q (1193A>C 1195T>G), Q134N (1205C>A 1207A>T), C136H (1211T>C 1212G>A), D144N (1235G>A), G147D (1245G>A), T149V (1250A>G 1251C>T), G154C (1265G>T), V157L (1274G>C 1276T>G), F163H (1292T>C 1293T>A), T170V (1313A>G 1314C>T), K171I (1317A>T 1318A>T), A174P (1325G>C 1327C>T), Q182T (1349C>A 1350A>C 1351A>T), I188M (1369T>G), Y189P (1370T>C 1371A>C 1372T>A), H194Q (1387C>A), N195D (1388A>G 1390T>C), S196P (1391T>C), V198I (1397G>A 1399A>T), L204V (1415C>G), E206D (1423A>T), E210H (1433G>C 1435A>C), G212N (1439G>A 1440G>A), L213I (1442T>A 1444G>T), K214E (1445A>G), I216R (1451A>C 1452T>G 1453T>A), I224R (1476T>G 1477T>A), A225C (1478G>T 1479C>G 1480C>T), S232A (1499T>G 1501T>C), H237Y (1514C>T), C240R (1523T>C), N250D (1553A>G 1555C>T), C253S (1563G>C 1564T>A), N254G (1565A>G 1566A>G), V258I (1577G>A), V259T (1580G>A 1581T>C), E261D (1588A>C), G262N (1589G>A 1590G>A), S263V (1592T>G 1593C>T 1594C>G), G265T (1598G>A 1599G>C 1600T>C), D268E (1609C>G), N269D (1610A>G 1612C>T), Q275S (1628C>A 1629A>G 1630A>T), K276R (1631A>C 1632A>G 1633A>T), K278R (1637A>C 1638A>G 1639A>T), K288H (1667A>C 1669A>T), I293V (1682A>G 1684C>T), V308I (1727G>A 1729G>T), E309D (1732A>C), V311I (1736G>A 1738G>A), G313S (1742G>A), A318S (1757G>T 1759A>T), Q321T (1766C>A 1767A>C 1768A>C), F329Y (1791T>A), A336P (1811G>C 1813T>C), K337V (1814A>G 1815A>T), E345Q (1838G>C), K347R (1845A>G), I349V (1850A>G 1852A>T), S351T (1857G>C 1858T>A), Y354C (1866A>G), A355G (1869C>G 1870A>T), A357P (1874G>C 1876A>C), E359Q (1880G>C), R362G (1889C>G), V364I (1895G>A 1897A>C), S369A (1910T>G 1912C>G), E373D (1924A>T), T374A (1925A>G 1927T>A), Q376N (1931C>A 1933A>C), N377H (1934A>C 1936T>C), V379I (1940G>A 1942G>T), R380P (1944G>C), V381D (1947T>A), K384R (1956A>G 1957G>A), I387V (1964A>G 1966A>C), Q395E (1988C>G 1990G>A), Y396Q (1991T>C 1993T>G), I401V (2006A>G 2008T>C), M405V (2018A>G 2020G>T), F406Y (2022T>A 2023C>T), A411L (2036G>C 2037C>T 2038T>C), N414S (2046A>G), L415V (2048C>G 2050A>C), V416I (2051G>A), V417I (2054G>A 2056A>T), I421V (2066A>G 2068T>A), V425L (2078G>C), L428Q (2087T>C 2088T>A), T434S (2105A>T), I436L (2111A>C 2113C>T), F437L (2116T>G), V440T (2123G>A 2124T>C), Y441V (2126T>G 2127A>T), K445R (2139A>G 2140A>G), V447I (2144G>A), L448F (2147C>T), D449E (2152T>A), L451I (2156C>A), E453A (2163A>C), F455L (2168T>C), K456S (2172A>G 2173G>T), E457A (2175A>C), R463K (2193G>A 2194A>G), G465A (2199G>C), V469L (2210G>C 2212T>C), I472L (2219A>C), S473I (2222T>A 2223C>T 2224A>T), C475G (2228T>G), A476V (2232C>T), C477F (2235G>T), E478D (2239A>C), G481K (2246G>A 2247G>A 2248T>G), V485Q (2258G>C 2259T>A 2260C>G), T486V (2261A>G 2262C>T 2263C>T), C487A (2264T>G 2265G>C), A488S (2267G>T), K489D (2270A>G 2272G>T), E490N (2273G>A 2275A>C), E493D (2284G>T), S494C (2285A>T), Q496K (2291C>A 2293G>A), T497C (2294A>T 2295C>G 2296A>C), F499I (2300T>A), K500D (2303A>G 2305G>T), L501V (2306C>G), F505A (2318T>G 2319T>C 2320T>A), A507E (2325C>A 2326T>A), L508M (2327T>A), A510I (2333G>A 2334C>T), S512Q (2339T>C 2340C>A 2341T>A), I513V (2342A>G), I514T (2346T>C), G516A (2352G>C), K521R (2366A>C 2367A>G), A522S (2369G>T 2371C>A), T528V (2387A>G 2388C>T 2389A>C), V530I (2393G>A), T531A (2396A>G 2398G>T), H532Q (2401C>A), K539Q (2420A>C), V541I (2426G>A 2428T>A), K542R (2429A>C 2430A>G 2431A>T), S543G (2432T>G 2433C>G), R544K (2436G>A 2437A>G), E546Q (2441G>C 2443A>G), T547L (2444A>C 2445C>T 2446T>G), G548Q (2447G>C 2448G>A 2449C>A), I559V (2480A>G 2482T>A), I560T (2484T>C), E565D (2500A>T), T566S (2501A>T), L567H (2505T>A), P568D (2507C>G 2508C>A), E570V (2514A>T), V571L (2516G>C 2518G>T), L572T (2519T>A 2520T>C 2521A>C), T573S (2522A>T 2524A>T), T580N (2544C>A 2545T>C), D582E (2551T>A), Q584E (2555C>G), P585A (2558C>G), Q588T (2567C>A 2568A>C 2569A>G), T590V (2573A>G 2574C>T), S591D (2576A>G 2577G>A), E592S (2579G>A 2580A>G 2581A>C), A593F (2582G>T 2583C>T 2584T>C), V594T (2585G>A 2586T>C 2587T>A), E595N (2588G>A 2590A>T), A596G (2592C>G 2593T>A), P597A (2594C>G 2596A>T), L598I (2597T>A 2599G>C), I605V (2618A>G 2620T>A), T616K (2652C>A), K618Q (2657A>C 2659G>A), A623S (2672G>T 2674A>T), N625G (2678A>G 2679A>G), M626L (2681A>T 2683G>A), M627L (2684A>C), V628A (2688T>C 2689A>T), T632V (2699A>G 2700C>T), T634R (2705A>C 2706C>G 2707A>C) | | | | | | | | |

| | Begin | End | Coverage | Score | Concordance | Matches | Identities | I/D/M/F* | Stop Codons |
|---|---|---|---|---|---|---|---|---|---|
| nsp3 (YP_009725299.1) | 1 | 1945 | 100% | 10414.0 | 80.5% | 1918 (98.4%) | 1485 (76.2%) | 4/27/0/0 | 0 |
| | T3I (2727C>T 2728A>T), K4_V5insG (2731_2732insGGT), D9E (2746T>A), I13W (2756A>T 2757T>G 2758A>G), S20N (2778G>A), N22R (2784A>G 2785T>A), I31V (2810A>G), A41V (2841C>T), L46S (2855C>T 2856T>C), N51T (2871A>C), D59E (2896T>G), I62V (2903A>G 2905A>G), E70D (2929A>T), P74N (2939C>A 2940C>A 2941A>C), L75M (2942C>A), M84V (2969A>G 2971G>A), Y87F (2979A>T), E92D (2995G>T), S93A (2996T>G), F96E (3005T>G 3006T>A 3007T>A), K97N (3010A>C), L98F (3013G>T), A99S (3014G>T 3016T>A), H101R (3021A>G), D112E (3055T>A), E115D (3064A>C), G116D (3066G>A), G116_D117insA (3067_3068insGCA), D117E (3070T>G), F123I (3086T>A), E124D (3091G>T), P125E (3092C>G 3093C>A), S126T (3095T>A 3097A>C), T127C (3098A>T 3099C>G), Q128E (3101C>G), Y129H (3104T>C), K140L (3137A>C 3138A>T 3139A>C), T147S (3158A>T 3160T>A), S148A (3161T>G), A149E (3165C>A 3166T>A), A150T (3167G>A 3169T>A), L151V (3170C>G), Q152R (3174A>G), P153V (3176C>G 3177C>T), Q157E (3188C>G), D165_T181del (3212_3262delGATAGTCAACAAACTGTTGGTCAACAAGACGGCAGTGAGGACAATCAGACA), I184E (3269A>G 3270T>A 3271T>G), T186S (3275A>T), I187_V188del (3278_3283delATTGTT), V190I (3287G>A), Q191E (3290C>G 3292A>G), Q193E (3296C>G), L194P (3299T>C 3300T>C), E195_M196del (3302_3307delGAGATG), L198P (3312T>C), V201E (3321T>A 3322T>A), V202E (3324T>A 3325T>A), Q203_I205del (3327_3335delAGACTATTG), E206P (3327_3335delAGACTATTG 3336A>C), S209Q (3344A>C 3345G>A 3346T>G), S211T (3351G>C), Y221A (3380T>G 3381A>C), N224C (3389A>T 3390A>G), A225V (3393C>T 3394A>T), E229K (3404G>A 3406A>G), K232Q (3413A>C), K233S (3417A>G 3418G>T), V234A (3420T>C 3421A>T), K235N (3424A>T), T237M (3429C>T 3430A>G), V239I (3434G>A), V245I (3452G>A 3454T>A), Y246H (3455T>C), N263G (3506A>G 3507A>G), V267K (3518G>A 3519T>A 3520T>G), A274K (3539G>A 3540C>A 3541T>G), T275L (3542A>C 3543C>T 3544T>A), K280T (3558A>C), V286L (3575G>T 3577T>G), H295K (3602C>A 3604C>G), V304L (3629G>C 3631T>A), K306A (3635A>G 3636A>C), S315A (3662A>G 3663G>C 3664T>A), Q322S (3683C>T 3684A>C 3685G>A), H323Q (3688C>G), E324D (3691A>C), V325I (3692G>A 3694T>C), D339K (3734G>A 3736C>A), I341L (3740A>C 3742A>T), H342Q (3745T>G), R345Q (3752A>C 3753G>A), D349Q (3764G>C 3766T>G), N354Q (3779A>C 3781T>G), L357I (3788T>A 3790A>T), F360N (3797T>A 3798T>A), N363A (3806A>G 3807A>C), D366E (3817C>G), K367Q (3818A>C 3820A>G), L368V (3821C>G), S370M (3827T>A 3828C>T 3829A>G), S371D (3830A>G 3831G>A 3832C>T), F372Y (3834T>A), E374D (3841A>T), M375N (3843T>A 3844G>C), K376L (3845A>C 3846A>T), S377K (3849G>A 3850T>G), E378P (3851G>C 3852A>C 3853A>T), K379R (3855A>G 3856G>A), Q380V (3857C>G 3858A>T 3859A>G), V381E (3861T>A 3862T>A), E382A (3864A>C), Q383P (3867A>C 3868A>T), I385Q (3872A>C 3873T>A 3874C>A), A386E (3876C>A 3877T>G), I388P (3881A>C 3882T>C 3883T>A), K390N (3889A>C), E391T (3890G>A 3891A>C 3892G>A), V393D (3897T>A), K394_P395del (3899_3904delAAGCCA), F396S (3906T>C 3907T>C), I397K (3909T>A), S400E (3917A>G 3918G>A 3919T>G), P402del (3923_3925delCCT), E405V (3933A>T), R407K (3939G>A 3940A>G), K408P (3941A>C 3942A>C 3943A>T), Q409V (3944C>G 3945A>T 3946A>C), D411V (3951A>T 3952T>G), K412_K413insP (3955_3956insCCA), V418I (3971G>A), E419D (3976A>T), E433N (4016G>A 4018A>T), N434K (4021C>G), Y438F (4032A>T), I439A (4034A>G 4035T>C), N444K (4051T>G), H446Y (4055C>T 4057T>C), P447H (4059C>A 4060A>T), A450Q (4067G>C 4068C>A 4069C>G), T451N (4071C>A 4072T>C), L452M (4073C>A 4075T>G), V453L (4076G>C), S454R (4081T>A), D455G (4083A>G 4084C>T), I456E (4085A>G 4086T>A 4087T>A), I458M (4093C>G), T459S (4094A>T), K462E (4103A>G), I468M (4123A>G), V473I (4136G>A 4138T>C), Q474T (4139C>A 4140A>C 4141A>T), E475S (4142G>A 4143A>G 4144G>T), V477D (4149T>A), L478I (4151T>A 4153A>C), A480C (4157G>T 4158C>G), T485S (4172A>T 4174T>C), A496S (4205G>T 4207G>A), K497R (4209A>G), R500K (4218G>A 4219A>G), T504V (4229A>G 4230C>T 4231A>T), N506E (4235A>G 4237T>G), L516C (4266T>G 4267A>T), N517A (4268A>G 4269A>C), V521L (4280G>C 4282A>T), V527A (4299T>C 4300G>T), I537V (4328A>G 4330T>A), I541E (4340A>G 4341T>A 4342T>A), I542A (4343A>G 4344T>C 4345C>A), S543P (4346T>C), E545A (4353A>C 4354G>T), Q547E (4358C>G), V573I (4436G>A 4438C>A), V575M (4442G>A), E576D (4447A>T), T577V (4448A>G 4449C>T), K578R (4452A>G), V581M (4460G>A 4462T>G), S582A (4463T>G), V597I (4508G>A 4510G>C), A602V (4524C>T 4525T>C), Y605F (4533A>T), T611E (4550A>G 4551C>A 4552A>G), T612P (4553A>C), L616I (4565C>A), N618T (4572A>C 4573C>G), T619K (4575C>A 4576A>G), D622S (4583G>T 4584A>C), T626P (4595A>C 4597T>G), L632I (4613C>A), L639F (4636A>T), Y647C (4659A>G), V653A (4677T>C 4678G>T), T656V (4685A>G 4686C>T), A667T (4718G>A 4720G>A), P679S (4754C>T), I684V (4769A>G 4771T>A), I687V (4778A>G 4780C>T), K694R (4800A>G), S702R (4823T>C 4824C>G), Q704E (4829C>G 4831A>G), I707V (4838A>G 4840A>T), S716I (4866G>T), Y719H (4874T>C), T720_S721insL (4879_4880insCTG), S721E (4880A>G 4881G>A 4882T>G), N722S (4884A>G 4885T>C), T724V (4889A>G 4890C>T), T725E (4892A>G 4893C>A 4894A>G), I733L (4916A>C 4918C>T), T734S (4919A>T 4921C>A), F735L (4922T>C), N737K (4930T>A), T740S (4938C>G 4939A>T), R748K (4962G>A), I759T (4995T>C), V765L (5012G>C), N793V (5096A>G 5097A>T 5098T>A), S794N (5099T>A 5100C>A 5101A>T), Y801F (5121A>T), N805S (5133A>G), V811S (5150G>A 5151T>G), T820L (5177A>C 5178C>T), P822E (5183C>G 5184C>A 5185T>G), Y840F (5238A>T 5239C>T), N844G (5249A>G 5250A>G), A859S (5294G>T 5296C>T), T860S (5298C>G), A861V (5301C>T 5302A>T), T864A (5309A>G), I868L (5321A>C 5323A>T), L870V (5327T>G 5329G>C), P874A (5339C>G), D879E (5356T>G), E888D (5383A>T), C900S (5417T>A), S915T (5463G>C 5464T>C), Y916H (5465T>C 5467C>T), F918L (5471T>C 5473T>A), D924E (5491T>A), C926A (5495T>G 5496G>C 5497C>A), T936H (5525A>C 5526C>A), Q940K (5537C>A 5539G>A), Q941T (5540C>A 5541A>C 5542G>T), K945T (5553A>C), E959D (5596A>T), Q960N (5597C>A 5599A>T), F961L (5600T>C), K963T (5607A>C), Q966S (5615C>T 5616A>C 5617G>C), T970V (5627A>G 5628C>T), K973R (5636A>C 5637A>G 5638A>T), Q974D (5639C>G 5641A>T), K977Q (5648A>C), P985S (5672C>T), Q995E (5702C>G), E997K (5708G>A), K999Q (5714A>C), H1000Q (5719T>A), T1004L (5729A>T 5730C>T 5731T>A), S1007N (5739G>A), K1019T (5775A>C 5776A>T), S1023A (5786T>G), C1029R (5804T>C 5806C>T), L1034H (5819T>C 5820T>A 5821A>C), S1038M (5831T>A 5832C>T 5833C>G), I1045V (5852A>G 5854T>G), N1053T (5877A>C 5878C>A), T1063S (5906A>T 5908T>G), V1070T (5927G>A 5928T>C), C1071Y (5931G>A 5932T>C), D1075E (5944C>A), N1080G (5957A>G 5958A>G 5959T>G), S1087A (5978T>G), F1089Y (5985T>A 5986C>T), N1099T (6015A>C 6016C>T), Y1102L (6024A>T 6025T>A), F1112L (6053T>C 6055T>C), V1113T (6056G>A 6057T>C), D1115S (6062G>T 6063A>C), I1117T (6069T>C 6070C>A), L1126M (6095T>A 6097A>G), Y1129F (6105A>T 6106T>C), K1130T (6108A>C 6109G>A), K1138S (6131A>T 6132A>C 6133A>T), K1155R (6183A>G), T1158S (6191A>T), P1159A (6194C>G 6196C>G), V1176I (6245G>A), N1178Q (6251A>C 6253T>G), N1181T (6261A>C 6262T>C), A1183T (6266G>A 6268C>A), Y1185F (6273A>T 6274T>C), I1192L (6293A>T), E1202D (6325A>T), D1208E (6343T>A), K1211A (6350A>G 6351A>C 6352G>A), S1212V (6353T>G 6354C>T), A1215T (6362G>A 6364G>A), D1225S (6392G>A 6393A>G), L1226Q (6396T>A), K1227Q (6398A>C), V1229T (6404G>A 6405T>C), D1242E (6445C>A), L1244I (6449C>A 6451T>A), N1247D (6458A>G 6460T>C), D1256N (6485G>A 6487C>T), I1257V (6488A>G 6490T>C), A1262S (6503G>T), N1263D (6506A>G), N1264E (6509A>G 6511T>A), S1265G (6512A>G), L1266V (6515T>G 6517A>T), I1268V (6521A>G 6523T>A), E1270Q (6527G>C), V1272L (6533G>T 6535T>A), T1275E (6542A>G 6543C>A 6544A>G), D1283E (6568C>A), S1285T (6572T>A 6574T>A), L1287I (6578C>A), R1297L (6608A>C 6609G>T), V1298A (6612T>C 6613A>C), L1304I (6629C>A), L1309I (6644T>A 6646A>T), V1312I (6653G>A), D1318S (6671G>A 6672A>G), T1319K (6675C>A 6676T>A), A1321L (6680G>T 6681C>T 6682T>G), N1322A (6683A>G 6684A>C), A1324V (6690C>T 6691T>C), N1329G (6704A>G 6705A>G 6706C>A), K1330Q (6707A>C), V1331A (6711T>C 6712T>A), V1332A (6714T>C 6715T>A), S1333I (6717G>T), T1336S (6725A>T 6727T>A), I1338C (6731A>T 6732T>G 6733A>C), V1339A (6735T>C), T1340K (6738C>A 6739A>G), C1342L (6744G>T 6745T>A), L1343A (6746T>G 6747T>C), N1344Q (6749A>C 6751C>A), C1347F (6759G>T), T1348N (6762C>A 6763T>C), F1354V (6779T>G 6781C>G), L1359F (6794C>T 6796A>C), R1366K (6816G>A), K1373R (6837A>G), M1376L (6845A>C 6847G>A), T1384S (6870C>G), G1389A (6885G>C), F1391L (6892T>A), E1394D (6901G>T), S1396G (6905T>G 6906C>G 6907A>C), F1397I (6908T>A), L1400V (6917T>G), N1404K (6931T>A), I1409F (6944A>T 6946A>C), N1410T (6948A>C 6949T>A), I1412A (6953A>G 6954T>C 6955A>T), I1413M (6958T>G), F1415L (6962T>C 6964T>A), V1420I (6977G>A), Y1427C (6999A>G 7000C>T), S1428V (7001T>G 7002C>T), L1432F (7015A>T), M1436L (7025A>T 7027G>A), L1439F (7036A>T), M1441A (7040A>G 7041T>C 7042G>T), T1446N (7056C>A), Y1448V (7061T>G 7062A>T 7063C>T), G1451L (7070G>T 7071G>T 7072C>G), T1456S (7085A>T), I1460T (7098T>C), A1461M (7100G>A 7101C>T 7102A>G), T1462D (7103A>G 7104C>A 7105C>T), Y1463F (7107A>T), T1465E (7112A>G 7113C>A 7114T>A), I1468F (7121A>T 7123A>T), V1472I (7133G>A), T1482S (7163A>T 7165C>T), S1485A (7172T>G), I1491V (7190A>G 7192T>G), F1496Y (7206T>A 7207T>C), W1498L (7211T>C 7212G>T 7213G>A), A1502I (7223G>A 7224C>T), F1503L (7228T>A), V1506A (7236T>C 7237T>C), F1510V (7247T>G), I1514M (7261T>G), R1518K (7272G>A 7273G>A), V1522L (7283G>T), A1526S (7295G>T 7297T>A), L1531V (7310T>G), S1534G (7319A>G), V1538S (7331G>A 7332T>G 7333A>T), L1549F (7366A>T), N1552S (7374A>G), L1553I (7376C>A), I1559V (7394A>G), V1575I (7442G>A), V1582I (7463G>A 7465T>C), V1583M (7466G>A 7468A>G), N1587T (7479A>C 7480T>C), V1612M (7553G>A 7555T>G), R1613K (7557G>A 7558A>G), K1624R (7589A>C 7590A>G 7591A>T), L1629T (7604C>A 7605T>C 7606A>T), V1635L (7622G>C 7624T>C), A1642T (7643G>A), T1677A (7748A>G 7750A>T), S1682A (7763T>G 7765C>G), I1683L (7766A>C 7768C>T), S1699P (7814T>C 7816T>G), E1732D (7915A>C), S1735A (7922T>G), A1736S (7925G>T 7927A>T), A1766T (8015G>A 8017G>T), A1769S (8024G>T 8026A>C), N1778D (8051A>G 8053T>C), S1782A (8063T>G), N1785S (8073A>G 8074C>T), T1793A (8096A>G), E1799H (8114G>C 8116A>C), A1800S (8117G>A 8118C>G 8119T>C), N1805G (8132A>G 8133A>G), S1807A (8138T>G 8140C>T), N1810G (8147A>G 8148A>G), I1816V (8165A>G 8167T>G), F1823V (8186T>G), S1826T (8195T>A 8197A>C), E1829D (8206A>C), V1834I (8219G>A), Q1842H (8245A>C), I1845L (8252A>T), Y1855F (8283A>T 8284T>C), S1877N (8349G>A), I1891V (8390A>G), A1892S (8393G>T 8395T>A), F1900Y (8418T>A), L1920I (8477T>A), K1923R (8487A>G 8488G>A), V1936I (8525G>A), A1941S (8540G>T) | | | | | | | | |
| nsp4 (YP_009725300.1) | 1 | 500 | 100% | 3038.0 | 86.1% | 500 (100%) | 400 (80.0%) | 0/0/0/0 | 0 |
| | N4S (8565A>G), N5T (8568A>C), W6C (8572G>T), L7F (8575G>T), Q9L (8580A>T 8581G>T), L10M (8582T>A 8584A>G), I11L (8585A>C), V13A (8592T>C 8593T>C), V16L (8600G>T), F17C (8604T>G), L18V (8606C>G), F19L (8609T>C), V20A (8613T>C), A22L (8618G>T 8619C>T 8620T>G), I23V (8621A>G), F24C (8625T>G 8626C>T), L26I (8630T>A 8632A>C), I27V (8633A>G 8635A>T), T28M (8637C>T 8638A>G), V32T (8648G>A 8649T>C 8650C>A), M33L (8651A>T), K35I (8658A>T 8659A>C), T37D (8663A>G 8664C>A), D38G (8667A>G 8668C>T), F39Y (8670T>A 8671T>C), S40T (8672T>A), S41N (8676G>A), D50Q (8702G>C 8704T>G), G51D (8706G>A), A58I (8726G>A 8727C>T 8728A>T), T62D (8738A>G 8739C>A), D70G (8763A>G), T73A (8771A>G), T83K (8802C>A 8803T>A), A87S (8813G>A 8814C>G 8815T>C), L90V (8822T>G 8824G>A), I91V (8825A>G 8827T>A), V94I (8834G>A), V99I (8849G>A 8851G>T), V102I (8858G>A 8860C>A), I110V (8882A>G 8884A>G), T113A (8891A>G), T114I (8895C>T 8896T>C), T143S (8982C>G), S163M (9041T>A 9042C>T 9043T>G), V175L (9077G>T 9079A>G), V180I (9092G>A), A181S (9095G>T), E183S (9101G>A 9102A>G 9103A>T), S184E (9104A>G 9105G>A 9106T>G), S218A (9206T>G), A231V (9246C>T 9247T>A), V233I (9251G>A), V235L (9257G>C), D246E (9292T>G), Y247H (9293T>C), S250A (9302T>G), P252S (9308C>T), V261M (9335G>A 9337A>G), L264I (9344C>A 9346T>A), T265A (9347A>G), M267I (9355G>C), I272V (9368A>G 9370T>G), I275V (9377A>G 9379T>G), I280V (9392A>G 9394A>G), I284V (9404A>G), V290I (9422G>A 9424A>T), V293L (9431G>T 9433A>G), L297A (9443C>G 9444T>C), R303K (9462G>A 9463G>A), A307V (9474C>T), S312N (9489G>A 9490T>C), F317A (9503T>G 9504T>C), T319A (9509A>G 9511T>A), V328I (9536G>A), T332V (9548A>G 9549C>T), V334A (9555T>C), I345F (9587A>T), L353F (9611C>T 9613T>C), I363L (9641A>C), M366F (9650A>T 9652G>T), V367A (9654T>C 9655T>C), T370S (9662A>T 9664A>T), L372I (9668T>A 9670A>T), I379A (9689A>G 9690T>C 9691T>A), A380I (9692G>A 9693C>T 9694T>C), I382V (9698A>G 9700C>A), I383F (9701A>T 9703T>C), T387L (9713A>C 9714C>T 9715A>G), F390C (9723T>G), Y391H (9725T>C), S395N (9738G>A 9739T>C), K399R (9750A>G), R400K (9753G>A), V403M (9761G>A 9763C>G), S408T (9776T>A 9778C>A), D433E (9853T>G), V434T (9854G>A 9855T>C 9856G>A), M458L (9926A>T 9928G>A), S483A (10001T>G) | | | | | | | | |
| 3C-like proteinase (YP_009725301.1) | 1 | 306 | 100% | 2172.0 | 97.4% | 306 (100%) | 294 (96.1%) | 0/0/0/0 | 0 |

| | Begin | End | Coverage | Score | Concordance | Matches | Identities | I/D/M/F* | Stop Codons |
|---|---|---|---|---|---|---|---|---|---|
| | V35T (10157G>A 10158T>C), S46A (10190T>G 10192T>A), N65S (10248A>G 10249T>C), V86L (10310G>C 10312A>G), K88R (10317A>G), A94S (10334G>T 10336C>T), F134H (10454T>C 10455T>A 10456C>T), N180K (10594C>A), V202L (10658G>T 10660T>A), S267A (10853T>G 10855A>T), A285T (10907G>A), L286I (10910T>A 10912A>T) | | | | | | | | |
| nsp6 (YP_009725302.1) | 1 | 290 | 100% | 1813.0 | 89.9% | 290 (100%) | 253 (87.2%) | 0/0/0/0 | 0 |
| | S1G (10973A>G), A2K (10976G>A 10977C>A 10978A>G), V3F (10979G>T 10981G>C), R5K (10986G>A), T6I (10989C>T 10990A>T), I7V (10991A>G 10993C>T), L14M (11012T>A), I18F (11024A>T 11026T>C), V24I (11042G>A), L37V (11081T>G 11083G>T), A46T (11108G>A), M47L (11111A>C 11113G>T), I50M (11122T>G), M52I (11128G>T), S53A (11129T>G), F55C (11136T>G), M58L (11144A>C), F59L (11147T>C), D99E (11269T>A), M100L (11270A>T), V101A (11274T>C), F108Y (11295T>A), K109R (11298A>G), V120L (11330G>T), G135A (11376G>C), L146I (11408T>A 11410G>T), I168V (11474A>G 11476A>T), V182I (11516G>A), G188A (11535G>C), M192V (11546A>G), C197Y (11562G>A), I199L (11567A>T 11569T>G), F200L (11572C>A), F220C (11631T>G), T222C (11636A>T 11637C>G 11638T>C), N264S (11763A>G), V278I (11804G>A) | | | | | | | | |
| nsp7 (YP_009725303.1) | 1 | 83 | 100% | 508.0 | 99.4% | 83 (100%) | 82 (98.8%) | 0/0/0/0 | 0 |
| | K70R (12051A>G) | | | | | | | | |
| nsp8 (YP_009725304.1) | 1 | 198 | 100% | 1210.0 | 98.0% | 198 (100%) | 193 (97.5%) | 0/0/0/0 | 0 |
| | F15Y (12135T>A), I132V (12485A>G 12487A>C), N136G (12497A>G 12498A>G 12499C>T), T145N (12525C>A 12526A>C), S173N (12609G>A 12610T>C) | | | | | | | | |
| nsp9 (YP_009725305.1) | 1 | 113 | 100% | 752.0 | 98.4% | 113 (100%) | 110 (97.3%) | 0/0/0/0 | 0 |
| | T34N (12786C>A 12787A>T), T35S (12788A>T 12790A>G), L48H (12827T>C 12828T>A 12829A>C) | | | | | | | | |
| nsp10 (YP_009725306.1) | 1 | 139 | 100% | 1061.0 | 98.7% | 139 (100%) | 135 (97.1%) | 0/0/0/0 | 0 |
| | A23P (13091G>C), K113R (13362A>G), M137L (13433A>T), L138M (13436C>A 13438T>G) | | | | | | | | |
| RNA-dependent RNA polymerase (YP_009725307.1) | 1 | 932 | 100% | 6561.0 | 97.4% | 932 (100%) | 898 (96.4%) | 0/0/0/0 | 0 |
| | Q5S (13454C>T 13455A>C), S6T (13457T>A), D40E (13560T>A), D62E (13626T>A), D63G (13628A>G), I66L (13636A>T 13638T>A), F77M (13669T>A 13671C>G), L90V (13708C>G), K98V (13732A>G 13733A>T 13734A>C), I106V (13756A>G), A185S (13993G>T 13995T>A), N198D (14032A>G), I223V (14107A>G), T225V (14113A>G 14114C>T 14115C>A), T226A (14116A>G 14118G>A), S229C (14125A>T 14127T>C), V233I (14137G>A), T252A (14194A>G), V257M (14209G>A 14211T>G), T259A (14215A>G), T262A (14224A>G), Y265L (14233T>C 14234A>T 14235C>T), K281C (14281A>T 14282A>G 14283A>T), V299I (14335G>A), N611T (15272A>C 15273C>T), T643N (15368C>A 15369A>C), S647N (15380G>A), T739H (15655A>C 15656C>A 15657A>T), D740E (15660C>A), N743D (15667A>G), F766Y (15737T>A 15738C>T), T769N (15746C>A 15747T>C), S772A (15754T>G), S784A (15790T>G) | | | | | | | | |
| helicase (YP_009725308.1) | 1 | 601 | 100% | 4241.0 | 99.9% | 601 (100%) | 600 (99.8%) | 0/0/0/0 | 0 |
| | V570I (17944G>A 17946A>T) | | | | | | | | |
| 3'-to-5' exonuclease (YP_009725309.1) | 1 | 527 | 100% | 3864.0 | 96.8% | 527 (100%) | 501 (95.1%) | 0/0/0/0 | 0 |
| | V14I (18079G>A 18081A>C), T31I (18131C>T 18132T>A), E99D (18336A>T), P128E (18421C>G 18422C>A 18423T>A), D132E (18435T>A), S134T (18439T>A), S137N (18449G>A), N176G (18565A>G 18566A>G 18567T>A), R212K (18674G>A), A220S (18697G>T 18699T>A), H228N (18721C>A), I231V (18730A>G 18732T>G), L259Q (18815T>A 18816G>A), Y260H (18817T>C), T293S (18916A>T), I294V (18919A>G), K304R (18950A>G), I305V (18952A>G), A307S (18958G>T 18960G>T), A319S (18994G>T), D345E (19074T>A), S374H (19159T>C 19160C>A 19161T>C), S396A (19225T>G 19227C>A), V437T (19348G>A 19349T>C), L493Q (19516T>C 19517T>A), V510I (19567G>A) | | | | | | | | |
| endoRNAse (YP_009725310.1) | 1 | 346 | 100% | 2174.0 | 92.0% | 346 (100%) | 307 (88.7%) | 0/0/0/0 | 0 |
| | F7Y (19640T>A), Q18H (19674A>C), Q19A (19675C>G 19676A>C 19677G>C), V22A (19685T>C), T30A (19708A>G), V38I (19732G>A), L42I (19744T>A 19746G>C), V69I (19825G>A 19827G>T), D91E (19893T>A), I96V (19906A>G), S103T (19927T>A 19929T>A), T114S (19961C>G 19962G>T), I115A (19963A>G 19964T>C), A117S (19969G>T 19971A>T), P118S (19972C>T), F122L (19986T>G), D128E (20004T>A), Q152T (20074C>A 20075A>C), V155K (20083G>A 20084T>A 20085A>G), K158A (20092A>G 20093A>C), L162V (20104C>G 20106T>C), A171S (20131G>T 20133C>A), Y179F (20156A>T), V185I (20173G>A), V186I (20176G>A 20178C>T), N199D (20215A>G 20217T>C), Q201E (20221C>G 20223A>G), E202D (20226A>T), I211T (20252T>C), E223Q (20287G>C 20289A>G), S243G (20347A>G 20349T>A), L251M (20371C>A), F258S (20393T>C 20394T>A), K259Q (20395A>C 20397G>A), E260D (20400A>T), F263L (20407T>C), E264K (20410G>A), V314I (20560G>A), T325A (20593A>G 20595A>T) | | | | | | | | |
| 2'-O-ribose methyltransferase (YP_009725311.1) | 1 | 298 | 100% | 1968.0 | 95.3% | 298 (100%) | 278 (93.3%) | 0/0/0/0 | 0 |
| | S1A (20659T>G 20661T>A), D32E (20754T>A), S33N (20756G>A), T35V (20761A>G 20762C>T 20763A>T), L36I (20764T>A), K135R (21062A>G), N138H (21070A>C), I153L (21115A>C 21117T>G), Q158K (21130C>A 21132A>G), V167I (21157G>A 21159G>A), I169V (21163A>G), A188S (21220G>T), C209A (21283T>G 21284G>C), R216K (21304C>A 21305G>A 21306C>G), V223T (21325G>A 21326T>C), G265N (21451G>A 21452G>A), L272Y (21473T>A 21474A>T), S276E (21484A>G 21485G>A 21486T>A), I290V (21526A>G), V294I (21538G>A) | | | | | | | | |
| orf1a polyprotein (YP_009725295.1) | 1 | 4406 | 100% | 25296.0 | 84.3% | 4379 (99.3%) | 3552 (80.5%) | 4/27/0/0 | 1 |

| Begin | End | Coverage | Score | Concordance | Matches | Identities | I/D/M/F* | Stop Codons |
|---|---|---|---|---|---|---|---|---|

P6L (282C>T), F8V (287T>G), V38A (378T>C), Q44E (395C>G), D48N (407G>A), V56L (431G>C 433T>G), R77L (494C>T 495G>T), T78S (498C>G 499T>C), A79T (500G>A 502A>C), P80N (503C>A 504C>A), V84K (515G>A 516T>A 517T>G), M85V (518A>G 520G>C), L92M (539C>A 541C>G), E93D (544A>C), E102I (569G>A 570A>T 571G>A), I114T (606T>C 607A>C), V116I (611G>A 613G>T), K120N (625G>T), A138I (677G>A 678C>T), F143Y (693T>A), Y154I (725T>A 726A>T), F157Y (735T>A), Q158E (737C>G), E159Q (740G>C), S166G (761A>G), V169A (771T>C 772T>A), T170L (773A>C 774C>T), M174T (786T>C 787G>T), Y182V (809T>G 810A>T), E199D (862G>T), L204F (875C>T), A211S (896G>T 898T>A), S212M (899T>A 900C>T 901A>G), F221Y (927T>A 928T>C), D223E (934C>G), T224S (935A>T 937T>G), E233D (964A>C), Y241F (987A>T), E246D (1003A>T), L251H (1016T>C 1017T>A 1018G>C), L259S (1040T>A 1041T>G 1042G>T), N267K (1066T>A), N272K (1081T>G), I280K (1104T>A), I281V (1106A>G), T283V (1112A>G 1113C>T 1114T>C), L293T (1142C>A 1143T>C), D294E (1147T>G), N310Q (1193A>C 1195T>G), Q314N (1205C>A 1207A>T), C316H (1211T>C 1212G>A), D324N (1235G>A), G327D (1245G>A), T329V (1250A>G 1251C>T), G334C (1265G>T), V337L (1274G>C 1276T>G), F343H (1292T>C 1293T>A), T350V (1313A>G 1314C>T), K351I (1317A>T 1318A>T), A354P (1325G>C 1327C>T), Q362T (1349C>A 1350A>C 1351A>T), I368M (1369T>G), Y369P (1370T>C 1371A>C 1372T>A), H374Q (1387C>A), N375D (1388A>G 1390T>C), S376P (1391T>C), V378I (1397G>A 1399A>T), L384V (1415C>G), E386D (1423A>T), E390H (1433G>C 1435A>C), G392N (1439G>A 1440G>A), L393I (1442T>A 1444G>T), K394E (1445A>G), I396R (1451A>C 1452T>G 1453T>A), I404R (1476T>G 1477T>A), A405C (1478G>T 1479C>G 1480C>T), S412A (1499T>G 1501T>C), H417Y (1514C>T), C420R (1523T>C), N430D (1553A>G 1555C>T), C433S (1563G>C 1564T>A), N434G (1565A>G 1566A>G), V438I (1577G>A), V439T (1580G>A 1581T>C), E441D (1588A>C), G442N (1589G>A 1590G>A), S443V (1592T>G 1593C>T 1594C>G), G445T (1598G>A 1599G>C 1600T>C), D448E (1609C>G), N449D (1610A>G 1612C>T), Q455S (1628C>A 1629A>G 1630A>T), K456R (1631A>C 1632A>G 1633A>T), K458R (1637A>C 1638A>G 1639A>T), K468H (1667A>C 1669A>T), I473V (1682A>G 1684C>T), V488I (1727G>A 1729G>T), E489D (1732A>C), V491I (1736G>A 1738G>A), G493S (1742G>A), A498S (1757G>T 1759A>T), Q501T (1766C>A 1767A>C 1768A>C), F509Y (1791T>A), A516P (1811G>C 1813T>C), K517V (1814A>G 1815A>T), E525Q (1838G>C), K527R (1845A>G), I529V (1850A>G 1852A>T), S531T (1857G>C 1858T>A), Y534C (1866A>G), A535G (1869C>G 1870A>T), A537P (1874G>C 1876A>C), E539Q (1880G>C), R542G (1889C>G), V544I (1895G>A 1897A>C), S549A (1910T>G 1912C>G), E553D (1924A>T), T554A (1925A>G 1927T>A), Q556N (1931C>A 1933A>C), N557H (1934A>C 1936T>C), V559I (1940G>A 1942G>T), R560P (1944G>C), V561D (1947T>A), K564R (1956A>G 1957G>A), I567V (1964A>G 1966A>C), Q575E (1988C>G 1990G>A), Y576Q (1991T>C 1993T>G), I581V (2006A>G 2008T>C), M585V (2018A>G 2020G>T), F586Y (2022T>A 2023C>T), A591L (2036G>C 2037C>T 2038T>C), N594S (2046A>G), L595V (2048C>G 2050A>C), V596I (2051G>A), V597I (2054G>A 2056A>T), I601V (2066A>G 2068T>A), V605L (2078G>C), L608Q (2087T>C 2088T>A), T614S (2105A>T), I616L (2111A>C 2113C>T), F617L (2116T>G), V620T (2123G>A 2124T>C), Y621V (2126T>G 2127A>T), K625R (2139A>G 2140A>G), V627I (2144G>A), L628F (2147C>T), D629E (2152T>A), L631I (2156C>A), E633A (2163A>C), F635L (2168T>C), K636S (2172A>G 2173G>T), E637A (2175A>C), R643K (2193G>A 2194A>G), G645A (2199G>C), V649L (2210G>C 2212T>C), I652L (2219A>C), S653I (2222T>A 2223C>T 2224A>T), C655G (2228T>G), A656V (2232C>T), C657F (2235G>T), E658D (2239A>C), G661K (2246G>A 2247G>A 2248T>G), V665Q (2258G>C 2259T>A 2260C>G), T666V (2261A>G 2262C>T 2263C>T), C667A (2264T>G 2265G>C), A668S (2267G>T), K669D (2270A>G 2272G>T), E670N (2273G>A 2275A>C), E673D (2284G>T), S674C (2285A>T), Q676K (2291C>A 2293G>A), T677C (2294A>T 2295C>G 2296A>C), F679I (2300T>A), K680D (2303A>G 2305G>T), L681V (2306C>G), F685A (2318T>G 2319T>C 2320T>A), A687E (2325C>A 2326T>A), L688M (2327T>A), A690I (2333G>A 2334C>T), S692Q (2339T>C 2340C>A 2341T>A), I693V (2342A>G), I694T (2346T>C), G696A (2352G>C), K701R (2366A>C 2367A>G), A702S (2369G>T 2371C>A), T708V (2387A>G 2388C>T 2389A>C), V710I (2393G>A), T711A (2396A>G 2398G>T), H712Q (2401C>A), K719Q (2420A>C), V721I (2426G>A 2428T>A), K722R (2429A>C 2430A>G 2431A>T), S723G (2432T>G 2433C>G), R724K (2436G>A 2437A>G), E726Q (2441G>C 2443A>G), T727L (2444A>C 2445C>T 2446T>G), G728Q (2447G>C 2448G>A 2449C>A), I739V (2480A>G 2482T>A), I740T (2484T>C), E745D (2500A>T), T746S (2501A>T), L747H (2505T>A), P748D (2507C>G 2508C>A), E750V (2514A>T), V751L (2516G>C 2518G>T), L752T (2519T>A 2520T>C 2521A>C), T753S (2522A>T 2524A>T), T760N (2544C>A 2545T>C), D762E (2551T>A), Q764E (2555C>G), P765A (2558C>G), Q768T (2567C>A 2568A>C 2569A>G), T770V (2573A>G 2574C>T), S771D (2576A>G 2577G>A), E772S (2579G>A 2580A>G 2581A>C), A773F (2582G>T 2583C>T 2584T>C), V774T (2585G>A 2586T>C 2587T>A), E775N (2588G>A 2590A>T), A776G (2592C>G 2593T>A), P777A (2594C>G 2596A>T), L778I (2597T>A 2599G>C), I785V (2618A>G 2620T>A), T796K (2652C>A), K798Q (2657A>C 2659G>A), A803S (2672G>T 2674A>T), N805G (2678A>G 2679A>G), M806L (2681A>T 2683G>A), M807L (2684A>C), V808A (2688T>C 2689A>T), T812V (2699A>G 2700C>T), T814R (2705A>C 2706C>G 2707A>C), T821I (2727C>T 2728A>T), K822_V823insG (2731_2732insGGT), D827E (2746T>A), I831W (2756A>T 2757T>G 2758A>G), S838N (2778G>A), N840R (2784A>G 2785T>A), I849V (2810A>G), A859V (2841C>T), L864S (2855C>T 2856T>C), N869T (2871A>C), D877E (2896T>G), I880V (2903A>G 2905A>G), E888D (2929A>T), P892N (2939C>A 2940C>A 2941A>C), L893M (2942C>A), M902V (2969A>G 2971G>A), Y905F (2979A>T), E910D (2995G>T), S911A (2996T>G), F914E (3005T>G 3006T>A 3007T>A), K915N (3010A>C), L916F (3013G>T), A917S (3014G>T 3016T>A), H919R (3021A>G), D930E (3055T>A), E933D (3064A>C), G934D (3066G>A), G934_D935insA (3067_3068insGCA), D935E (3070T>G), F941I (3086T>A), E942D (3091G>T), P943E (3092C>G 3093C>A), S944T (3095T>A 3097A>C), T945C (3098A>T 3099C>G), Q946E (3101C>G), Y947H (3104T>C), K958L (3137A>C 3138A>T 3139A>C), T965S (3158A>T 3160T>A), S966A (3161T>G), A967E (3165C>A 3166T>A), A968T (3167G>A 3169T>A), L969V (3170C>G), Q970R (3174A>G), P971V (3176C>G 3177C>T), Q975E (3188C>G), D983_T999del (3212_3262delGATAGTCAACAAACTGTTGGTCAACAAGACGGCAGTGAGGACAATCAGACA), I1002E (3269A>G 3270T>A 3271T>G), T1004S (3275A>T), I1005_V1006del (3278_3283delATTGTT), V1008I (3287G>A), Q1009E (3290C>G 3292A>G), Q1011E (3296C>G), L1012P (3299T>C 3300T>C), E1013_M1014del (3302_3307delGAGATG), L1016P (3312T>C), V1019E (3321T>A 3322T>A), V1020E (3324T>A 3325T>A), Q1021_I1023del (3327_3335delAGACTATTG), E1024P (3327_3335delAGACTATTG 3336A>C), S1027Q (3344A>C 3345G>A 3346T>G), S1029T (3351G>C), Y1039A (3380T>G 3381A>C), N1042C (3389A>T 3390A>G), A1043V (3393C>T 3394A>T), E1047K (3404G>A 3406A>G), K1050Q (3413A>C), K1051S (3417A>G 3418G>T), V1052A (3420T>C 3421A>T), K1053N (3424A>T), T1055M (3429C>T 3430A>G), V1057I (3434G>A), V1063I (3452G>A 3454T>A), Y1064H (3455T>C), N1081G (3506A>G 3507A>G), V1085K (3518G>A 3519T>A 3520T>G), A1092K (3539G>A 3540C>A 3541T>G), T1093L (3542A>C 3543C>T 3544T>A), K1098T (3558A>C), V1104L (3575G>T 3577T>G), H1113K (3602C>A 3604C>G), V1122L (3629G>C 3631T>A), K1124A (3635A>G 3636A>C), S1133A (3662A>G 3663G>C 3664T>A), Q1140S (3683C>T 3684A>C 3685G>A), H1141Q (3688C>G), E1142D (3691A>C), V1143I (3692G>A 3694T>C), D1157K (3734G>A 3736C>A), I1159L (3740A>C 3742A>T), H1160Q (3745T>G), R1163Q (3752A>C 3753G>A), D1167Q (3764G>C 3766T>G), N1172Q (3779A>C 3781T>G), L1175I (3788T>A 3790A>T), F1178N (3797T>A 3798T>A), N1181A (3806A>G 3807A>C), D1184E (3817C>G), K1185Q (3818A>C 3820A>G), L1186V (3821C>G), S1188M (3827T>A 3828C>T 3829A>G), S1189D (3830A>G 3831G>A 3832C>T), F1190Y (3834T>A), E1192D (3841A>T), M1193N (3843T>A 3844G>C), K1194L (3845A>C 3846A>T), S1195K (3849G>A 3850T>G), E1196P (3851G>C 3852A>C 3853A>T), K1197R (3855A>G 3856G>A), Q1198V (3857C>G 3858A>T 3859A>G), V1199E (3861T>A 3862T>A), E1200A (3864A>C), Q1201P (3867A>C 3868A>T), I1203Q (3872A>C 3873T>A 3874C>A), A1204E (3876C>A 3877T>G), I1206P (3881A>C 3882T>C 3883T>A), K1208N (3889A>C), E1209T (3890G>A 3891A>C 3892G>A), V1211D (3897T>A), K1212_P1213del (3899_3904delAAGCCA), F1214S (3906T>C 3907T>C), I1215K (3909T>A), S1218E (3917A>G 3918G>A 3919T>G), P1220del (3923_3925delCCT), E1223V (3933A>T), R1225K (3939G>A 3940A>G), K1226P (3941A>C 3942A>C 3943A>T), Q1227V (3944C>G 3945A>T 3946A>C), D1229V (3951A>T 3952T>G), K1230_K1231insP (3955_3956insCCA), V1236I (3971G>A), E1237D (3976A>T), E1251N (4016G>A 4018A>T), N1252K (4021C>G), Y1256F (4032A>T), I1257A (4034A>G 4035T>C), N1262K (4051T>G), H1264Y (4055C>T 4057T>C), P1265H (4059C>A 4060A>T), A1268Q (4067G>C 4068C>A 4069C>G), T1269N (4071C>A 4072T>C), L1270M (4073C>A 4075T>G), V1271L (4076G>C), S1272R (4081T>A), D1273G (4083A>G 4084C>T), I1274E (4085A>G 4086T>A 4087T>A), I1276M (4093C>G), T1277S (4094A>T), K1280E (4103A>G), I1286M (4123A>G), V1291I (4136G>A 4138T>C), Q1292T (4139C>A 4140A>C 4141A>T), E1293S (4142G>A 4143A>G 4144G>T), V1295D (4149T>A), L1296I (4151T>A 4153A>C), A1298C (4157G>T 4158C>G), T1303S (4172A>T 4174T>C), A1314S (4205G>T 4207G>A), K1315R (4209A>G), R1318K (4218G>A 4219A>G), T1322V (4229A>G 4230C>T 4231A>T), N1324E (4235A>G 4237T>G), L1334C (4266T>G 4267A>T), N1335A (4268A>G 4269A>C), V1339L (4280G>C 4282A>T), V1345A (4299T>C 4300G>T), I1355V (4328A>G 4330T>A), I1359E (4340A>G 4341T>A 4342T>A), I1360A (4343A>G 4344T>C 4345C>A), S1361P (4346T>C), E1363A (4353A>C 4354G>T), Q1365E (4358C>G), V1391I (4436G>A 4438C>A), V1393M (4442G>A), E1394D (4447A>T), T1395V (4448A>G 4449C>T), K1396R (4452A>G), V1399M (4460G>A 4462T>G), S1400A (4463T>G), V1415I (4508G>A 4510G>C), A1420V (4524C>T 4525T>C), Y1423F (4533A>T), T1429E (4550A>G 4551C>A 4552A>G), T1430P (4553A>C), L1434I (4565C>A), N1436T (4572A>C 4573C>G), T1437K (4575C>A 4576A>G), D1440S (4583G>T 4584A>C), T1444P (4595A>C 4597T>G), L1450I (4613C>A), L1457F (4636A>T), Y1465C (4659A>G), V1471A (4677T>C 4678G>T), T1474V (4685A>G 4686C>T), A1485T (4718G>A 4720G>A), P1497S (4754C>T), I1502V (4769A>G 4771T>A), I1505V (4778A>G 4780C>T), K1512R (4800A>G), S1520R (4823T>C 4824C>G), Q1522E (4829C>G 4831A>G), I1525V (4838A>G 4840A>T), S1534I (4866G>T), Y1537H (4874T>C), T1538_S1539insL (4879_4880insCTG), S1539E (4880A>G 4881G>A 4882T>G), N1540S (4884A>G 4885T>C), T1542V (4889A>G 4890C>T), T1543E (4892A>G 4893C>A 4894A>G), I1551L (4916A>C 4918C>T), T1552S (4919A>T 4921C>A), F1553L (4922T>C), N1555K (4930T>A), T1558S (4938C>G 4939A>T), R1566K (4962G>A), I1577T (4995T>C), V1583L (5012G>C), N1611V (5096A>G 5097A>T 5098T>A), S1612N (5099T>A 5100C>A 5101A>T), Y1619F (5121A>T), N1623S (5133A>G), V1629S (5150G>A 5151T>G), T1638L (5177A>C 5178C>T), P1640E (5183C>G 5184C>A 5185T>G), Y1658F (5238A>T 5239C>T), N1662G (5249A>G 5250A>G), A1677S (5294G>T 5296C>T), T1678S (5298C>G), A1679V (5301C>T 5302A>T), T1682A (5309A>G), I1686L (5321A>C 5323A>T), L1688V (5327T>G 5329G>C), P1692A (5339C>G), D1697E (5356T>G), E1706D (5383A>T), C1718S (5417T>A), S1733T (5463G>C 5464T>C), Y1734H (5465T>C 5467C>T), F1736L (5471T>C 5473T>A), D1742E (5491T>A), C1744A (5495T>G 5496G>C 5497C>A), T1754H (5525A>C 5526C>A), Q1758K (5537C>A 5539G>A), Q1759T (5540C>A 5541A>C 5542G>T), K1763T (5553A>C), E1777D (5596A>T), Q1778N (5597C>A 5599A>T), F1779L (5600T>C), K1781T (5607A>C), Q1784S (5615C>T 5616A>C 5617G>C), T1788V (5627A>G 5628C>T), K1791R (5636A>C 5637A>G 5638A>T), Q1792D (5639C>G 5641A>T), K1795Q (5648A>C), P1803S (5672C>T), Q1813E (5702C>G), E1815K (5708G>A), K1817Q (5714A>C), H1818Q (5719T>A), T1822L (5729A>T 5730C>T 5731T>A), S1825N (5739G>A), K1837T (5775A>C 5776A>T), S1841A (5786T>G), C1847R (5804T>C 5806C>T), L1852H (5819T>C 5820T>A 5821A>C), S1856M (5831T>A 5832C>T 5833C>G), I1863V (5852A>G 5854T>G), N1871T (5877A>C 5878C>A), T1881S (5906A>T 5908T>G), V1888T (5927G>A 5928T>C), C1889Y (5931G>A 5932T>C), D1893E (5944C>A), N1898G (5957A>G 5958A>G 5959T>G), S1905A (5978T>G), F1907Y (5985T>A 5986C>T), N1917T (6015A>C 6016C>T), Y1920L (6024A>T 6025T>A), F1930L (6053T>C 6055T>C), V1931T (6056G>A 6057T>C), D1933S (6062G>T 6063A>C), I1935T (6069T>C 6070C>A), L1944M (6095T>A 6097A>G), Y1947F (6105A>T 6106T>C), K1948T (6108A>C 6109G>A), K1956S (6131A>T 6132A>C 6133A>T), K1973R (6183A>G), T1976S (6191A>T), P1977A (6194C>G 6196C>G), V1994I (6245G>A), N1996Q (6251A>C 6253T>G), N1999T (6261A>C 6262T>C), A2001T (6266G>A 6268C>A), Y2003F (6273A>T 6274T>C), I2010L (6293A>T), E2020D (6325A>T), D2026E (6343T>A), K2029A (6350A>G 6351A>C 6352G>A), S2030V (6353T>G 6354C>T), A2033T (6362G>A 6364G>A), D2043S (6392G>A 6393A>G), L2044Q (6396T>A), K2045Q (6398A>C), V2047T (6404G>A 6405T>C), D2060E (6445C>A), L2062I (6449C>A 6451T>A), N2065D (6458A>G 6460T>C), D2074N (6485G>A 6487C>T), I2075V (6488A>G 6490T>C), A2080S (6503G>T), N2081D (6506A>G), N2082E (6509A>G 6511T>A), S2083G (6512A>G), L2084V (6515T>G 6517A>T), I2086V (6521A>G 6523T>A), E2088Q (6527G>C), V2090L (6533G>T 6535T>A), T2093E (6542A>G 6543C>A 6544A>G), D2101E (6568C>A), S2103T (6572T>A 6574T>A), L2105I (6578C>A), R2115L (6608A>C 6609G>T), V2116A (6612T>C 6613A>C), L2122I (6629C>A), L2127I (6644T>A 6646A>T), V2130I (6653G>A), D2136S (6671G>A 6672A>G), T2137K (6675C>A 6676T>A), A2139L (6680G>T 6681C>T 6682T>G), N2140A (6683A>G 6684A>C), A2142V (6690C>T 6691T>C), N2147G (6704A>G 6705A>G 6706C>A), K2148Q (6707A>C), V2149A (6711T>C 6712T>A), V2150A (6714T>C 6715T>A), S2151I (6717G>T), T2154S (6725A>T 6727T>A), I2156C (6731A>T 6732T>G 6733A>C), V2157A (6735T>C), T2158K (6738C>A 6739A>G), C2160L (6744G>T 6745T>A), L2161A (6746T>G 6747T>C), N2162Q (6749A>C 6751C>A), C2165F (6759G>T), T2166N (6762C>A 6763T>C), F2172V (6779T>G 6781C>G), L2177F (6794C>T 6796A>C), R2184K (6816G>A), K2191R (6837A>G), M2194L (6845A>C 6847G>A), T2202S (6870C>G), G2207A (6885G>C), F2209L (6892T>A), E2212D (6901G>T), S2214G (6905T>G 6906C>G 6907A>C), F2215I (6908T>A), L2218V (6917T>G), N2222K (6931T>A), I2227F (6944A>T 6946A>C), N2228T (6948A>C 6949T>A), I2230A (6953A>G 6954T>C 6955A>T), I2231M (6958T>G), F2233L (6962T>C 6964T>A), V2238I (6977G>A), Y2245C (6999A>G 7000C>T), S2246V (7001T>G 7002C>T), L2250F (7015A>T), M2254L (7025A>T 7027G>A), L2257F (7036A>T), M2259A (7040A>G 7041T>C 7042G>T), T2264N (7056C>A), Y2266V (7061T>G 7062A>T 7063C>T), G2269L (7070G>T 7071G>T 7072C>G), T2274S (7085A>T), I2278T (7098T>C), A2279M (7100G>A 7101C>T 7102A>G), T2280D (7103A>G 7104C>A 7105C>T), Y2281F (7107A>T), T2283E

| | | | | | | | | | |
|---|---|---|---|---|---|---|---|---|---|
| | (7112A>G 7113C>A 7114T>A), I2286F (7121A>T 7123A>T), V2290I (7133G>A), T2300S (7163A>T 7165C>T), S2303A (7172T>G), I2309V (7190A>G 7192T>G), F2314Y (7206T>A 7207T>C), W2316L (7211T>C 7212G>T 7213G>A), A2320I (7223G>A 7224C>T), F2321L (7228T>A), V2324A (7236T>C 7237T>C), F2328V (7247T>G), I2332M (7261T>G), R2336K (7272G>A 7273G>A), V2340L (7283G>T), A2344S (7295G>T 7297T>A), L2349V (7310T>G), S2352G (7319A>G), V2356S (7331G>A 7332T>G 7333A>T), L2367F (7366A>T), N2370S (7374A>G), L2371I (7376C>A), I2377V (7394A>G), V2393I (7442G>A), V2400I (7463G>A 7465T>C), V2401M (7466G>A 7468A>G), N2405T (7479A>C 7480T>C), V2430M (7553G>A 7555T>G), R2431K (7557G>A 7558A>G), K2442R (7589A>C 7590A>G 7591A>T), L2447T (7604C>A 7605T>C 7606A>T), V2453L (7622G>C 7624T>C), A2460T (7643G>A), T2495A (7748A>G 7750A>T), S2500A (7763T>G 7765C>G), I2501L (7766A>C 7768C>T), S2517P (7814T>C 7816T>G), E2550D (7915A>C), S2553A (7922T>G), A2554S (7925G>T 7927A>T), A2584T (8015G>A 8017G>T), A2587S (8024G>T 8026A>C), N2596D (8051A>G 8053T>C), S2600A (8063T>G), N2603S (8073A>G 8074C>T), T2611A (8096A>G), E2617H (8114G>C 8116A>C), A2618S (8117G>A 8118C>G 8119T>C), N2623G (8132A>G 8133A>G), S2625A (8138T>G 8140C>T), N2628G (8147A>G 8148A>G), I2634V (8165A>G 8167T>G), F2641V (8186T>G), S2644T (8195T>A 8197A>C), E2647D (8206A>C), V2652I (8219G>A), Q2660H (8245A>C), I2663L (8252A>T), Y2673F (8283A>T 8284T>C), S2695N (8349G>A), I2709V (8390A>G), A2710S (8393G>T 8395T>A), F2718Y (8418T>A), L2738I (8477T>A), K2741R (8487A>G 8488G>A), V2754I (8525G>A), A2759S (8540G>T), N2767S (8565A>G), N2768T (8568A>C), W2769C (8572G>T), L2770F (8575G>T), Q2772L (8580A>T 8581G>T), L2773M (8582T>A 8584A>G), I2774L (8585A>C), V2776A (8592T>C 8593T>C), V2779L (8600G>T), F2780C (8604T>G), L2781V (8606C>G), F2782L (8609T>C), V2783A (8613T>C), A2785L (8618G>T 8619C>T 8620T>G), I2786V (8621A>G), F2787C (8625T>G 8626C>T), L2789I (8630T>A 8632A>C), I2790V (8633A>G 8635A>T), T2791M (8637C>T 8638A>G), V2795T (8648G>A 8649T>C 8650C>A), M2796L (8651A>T), K2798I (8658A>T 8659A>C), T2800D (8663A>G 8664C>A), D2801G (8667A>G 8668C>T), F2802Y (8670T>A 8671T>C), S2803T (8672T>A), S2804N (8676G>A), D2813Q (8702G>C 8704T>G), G2814D (8706G>A), A2821I (8726G>A 8727C>T 8728A>T), T2825D (8738A>G 8739C>A), D2833G (8763A>G), T2836A (8771A>G), T2846K (8802C>A 8803T>A), A2850S (8813G>A 8814C>G 8815T>C), L2853V (8822T>G 8824G>A), I2854V (8825A>G 8827T>A), V2857I (8834G>A), V2862I (8849G>A 8851G>T), V2865I (8858G>A 8860C>A), I2873V (8882A>G 8884A>G), T2876A (8891A>G), T2877I (8895C>T 8896T>C), T2906S (8982C>G), S2926M (9041T>A 9042C>T 9043T>G), V2938L (9077G>T 9079A>G), V2943I (9092G>A), A2944S (9095G>T), E2946S (9101G>A 9102A>G 9103A>T), S2947E (9104A>G 9105G>A 9106T>G), S2981A (9206T>G), A2994V (9246C>T 9247T>A), V2996I (9251G>A), V2998L (9257G>C), D3009E (9292T>G), Y3010H (9293T>C), S3013A (9302T>G), P3015S (9308C>T), V3024M (9335G>A 9337A>G), L3027I (9344C>A 9346T>A), T3028A (9347A>G), M3030I (9355G>C), I3035V (9368A>G 9370T>G), I3038V (9377A>G 9379T>G), I3043V (9392A>G 9394A>G), I3047V (9404A>G), V3053I (9422G>A 9424A>T), V3056L (9431G>T 9433A>G), L3060A (9443C>G 9444T>C), R3066K (9462G>A 9463G>A), A3070V (9474C>T), S3075N (9489G>A 9490T>C), F3080A (9503T>G 9504T>C), T3082A (9509A>G 9511T>A), V3091I (9536G>A), T3095V (9548A>G 9549C>T), V3097A (9555T>C), I3108F (9587A>T), L3116F (9611C>T 9613T>C), I3126L (9641A>C), M3129F (9650A>T 9652G>T), V3130A (9654T>C 9655T>C), T3133S (9662A>T 9664A>T), L3135I (9668T>A 9670A>T), I3142A (9689A>G 9690T>C 9691T>A), A3143I (9692G>A 9693C>T 9694T>C), I3145V (9698A>G 9700C>A), I3146F (9701A>T 9703T>C), T3150L (9713A>C 9714C>T 9715A>G), F3153C (9723T>G), Y3154H (9725T>C), S3158N (9738G>A 9739T>C), K3162R (9750A>G), R3163K (9753G>A), V3166M (9761G>A 9763C>G), S3171T (9776T>A 9778C>A), D3196E (9853T>G), V3197T (9854G>A 9855T>C 9856G>A), M3221L (9926A>T 9928G>A), S3246A (10001T>G), V3298T (10157G>A 10158T>C), S3309A (10190T>G 10192T>A), N3328S (10248A>G 10249T>C), V3349L (10310G>C 10312A>G), K3351R (10317A>G), A3357S (10334G>T 10336C>T), F3397H (10454T>C 10455T>A 10456C>T), N3443K (10594C>A), V3465L (10658G>T 10660T>A), S3530A (10853T>G 10855A>T), A3548T (10907G>A), L3549I (10910T>A 10912A>T), S3570G (10973A>G), A3571K (10976G>A 10977C>A 10978A>G), V3572F (10979G>T 10981G>C), R3574K (10986G>A), T3575I (10989C>T 10990A>T), I3576V (10991A>G 10993C>T), L3583M (11012T>A), I3587F (11024A>T 11026T>C), V3593I (11042G>A), L3606V (11081T>G 11083G>T), A3615T (11108G>A), M3616L (11111A>C 11113G>T), I3619M (11122T>G), M3621I (11128G>T), S3622A (11129T>G), F3624C (11136T>G), M3627L (11144A>C), F3628L (11147T>C), D3668E (11269T>A), M3669L (11270A>T), V3670A (11274T>C), F3677Y (11295T>A), K3678R (11298A>G), V3689L (11330G>T), G3704A (11376G>C), L3715I (11408T>A 11410G>T), I3737V (11474A>G 11476A>T), V3751I (11516G>A), G3757A (11535G>C), M3761V (11546A>G), C3766Y (11562G>A), I3768L (11567A>T 11569T>G), F3769L (11572C>A), F3789C (11631T>G), T3791C (11636A>T 11637C>G 11638T>C), N3833S (11763A>G), V3847I (11804G>A), K3929R (12051A>G), F3957Y (12135T>A), I4074V (12485A>G 12487A>C), N4078G (12497A>G 12498A>G 12499C>T), T4087N (12525C>A 12526A>C), S4115N (12609G>A 12610T>C), T4174N (12786C>A 12787A>T), T4175S (12788A>T 12790A>G), L4188H (12827T>C 12828T>A 12829A>C), A4276P (13091G>C), K4366R (13362A>G), M4390L (13433A>T), L4391M (13436C>A 13438T>G), Q4397S (13454C>T 13455A>C), S4398T (13457T>A) | | | | | | | | |
| nsp11 (YP_009725312.1) | 1 | 13 | 100% | 71.0 | 86.6% | 13 (100%) | 11 (84.6%) | 0/0/0/0 | 0 |
| | Q5S (13454C>T 13455A>C), S6T (13457T>A) | | | | | | | | |
| surface glycoprotein (YP_009724390.1) | 1 | 1274 | 100% | 7176.0 | 82.1% | 1250 (97.7%) | 974 (76.1%) | 6/24/0/0 | 1 |
| | V3I (21569G>A), V6L (21578G>T 21580T>A), L7F (21583A>T), P9T (21587C>A 21589A>T), V11T (21593G>A 21594T>C 21595C>T), S13G (21599A>G), S13_Q14insSDLD (21601_21602insAGTGACCTTGAC), Q14R (21603A>G), V16T (21608G>A 21609T>C 21610T>C), N17T (21612A>C), L18F (21614C>T), T19D (21617A>G 21618C>A 21619A>T), T20D (21620A>G 21621C>A 21622C>T), R21del (21623_21625delAGA), T22V (21626A>G 21627C>T), L24del (21632_21634delTTA), P25A (21635C>G 21637C>T), A27N (21641G>A 21642C>A 21643A>T), T29_N30insQH (21649_21650insCAACAT), N30T (21651A>C), F32S (21657T>C 21658C>T), T33M (21660C>T 21661A>G), K41E (21683A>G), V42I (21686G>A), S46D (21698T>G 21699C>A 21700A>C), V47T (21701G>A 21702T>C), H49Y (21707C>T), S50L (21711C>T), F59Y (21738T>A), W64G (21752T>G), A67T (21761G>A), H69_T73del (21767_21781delCATGTCTCTGGGACC), G75H (21785G>C 21786G>A), K77_R78del (21790_21795delTAAGAG), D80G (21801A>G 21802T>C), L84I (21812C>A), N87K (21823T>G), V90I (21830G>A), S94A (21842T>G), I100V (21860A>G 21862A>T), I101V (21863A>G 21865A>C), I105V (21875A>G), T108S (21884A>T), L110M (21890T>A 21892A>G), D111N (21893G>A 21895T>C), S112N (21896T>A 21897C>A 21898G>C), T114S (21902A>T 21904C>A), L117V (21911C>G 21913A>G), L118I (21914C>A), V120I (21920G>A), A123S (21929G>T), K129R (21947A>C 21948A>G), V130A (21951T>C 21952C>A), E132N (21956G>A 21958A>C), Q134E (21962C>G), F135L (21967T>G), N137D (21971A>G 21973T>C), D138N (21974G>A 21976T>C), L141F (21985G>T), G142A (21987G>C), Y144S (21993A>C), Y145K (21995T>A 21997C>A), H146P (21999A>C), K147M (22002A>T 22003A>G), N148_S151del (22004_22015delAACAACAAAAGT), W152G (22016T>G 22018G>T), M153T (22020T>C 22021G>A), E154Q (22022G>C 22024A>G), S155T (22026G>C 22027T>A), E156H (22028G>C 22030G>T), F157T (22031T>A 22032T>C 22033C>T), R158M (22035G>T 22036A>G), V159I (22037G>A 22039T>A), Y160F (22041A>T 22042T>C), S161D (22043T>G 22044C>A), S162N (22047G>A), N164F (22052A>T 22053A>T), V171I (22073G>A 22075C>A), Q173D (22079C>G 22081G>T), P174A (22082C>G 22084T>C), L176S (22088C>T 22089T>C 22090T>G), M177L (22091A>C 22093G>T), L179V (22097C>G), E180S (22100G>T 22101A>C), G181E (22104G>A), Q183S (22109C>T 22110A>C 22111G>A), N188H (22124A>C 22126T>C), I197K (22152T>A 22153T>A), Y200F (22161A>T), F201L (22163T>C 22165T>C), K202Y (22166A>T 22168A>T), I203V (22169A>G 22171A>T), S205K (22175T>A 22176C>A 22177T>G), K206G (22178A>G 22179A>G 22180G>C), H207Y (22181C>T 22183C>T), T208Q (22184A>C 22185C>A 22186G>A), N211D (22193A>G), L212V (22196T>G), Q218S (22214C>T 22215A>C 22216G>T), S221N (22223T>A 22224C>A 22225G>C), A222T (22226G>A), E224K (22232G>A), L226I (22238T>A 22240G>T), V227F (22241G>T 22243A>T), D228K (22244G>A 22246T>G), I231L (22253A>C 22255A>T), R237N (22272G>A 22273G>T), Q239R (22277C>A 22278A>G), T240A (22280A>G 22282T>C), L241I (22283T>A 22285A>T), A243T (22289G>A 22291T>A), L244A (22292T>G 22293T>C 22294A>C), H245_Y248del (22295_22306delCATAGAAGTTAT), L249F (22309G>T), T250S (22310A>T 22312T>A), G252A (22317G>C), D253Q (22319G>C 22321T>A), S254D (22322T>G 22323C>A 22324T>C), S255I (22325T>A 22326C>T), S256_G257del (22328_22333delTCAGGT), T259G (22337A>G 22338C>G 22339A>C), A260T (22340G>A 22342T>G), G261S (22343G>T 22344G>C 22345T>A), Y266F (22359A>T), Q271K (22373C>A 22375A>G), R273T (22380G>C 22381G>T), L276M (22388C>A 22390A>G), N280D (22400A>G), A292S (22436G>T 22438A>T), L293Q (22440T>A 22441T>A), D294N (22442G>A 22444C>T), S297A (22451T>G 22453A>T), T299L (22457A>C 22458C>T 22459A>C), T302S (22466A>T 22468G>T), L303V (22469T>G 22471G>T), T307E (22481A>G 22482C>A 22483T>G), V308I (22484G>A 22486A>T), E309D (22489A>C), Q321V (22523C>G 22524A>T 22525A>T), T323S (22529A>T), E324G (22533A>G), S325D (22535T>G 22536C>A), I326V (22538A>G), R346K (22599G>A), A348P (22604G>C 22606A>T), N354E (22622A>G 22624C>G), R357K (22632G>A), A372T (22676G>A), S373F (22680C>T 22681A>T), P384A (22712C>G 22714T>C), T393S (22739A>T 22741T>C), I402V (22766A>G 22768T>C), R403K (22770G>A 22771A>G), E406D (22780A>T), K417V (22811A>G 22812A>T 22813G>T), T430M (22851C>T 22852A>G), I434L (22862A>C 22864A>T), S438T (22874T>A), N439R (22878A>G 22879C>G), L441I (22883C>A), S443A (22889T>G), K444T (22893A>C 22894G>T), V445S (22895G>T 22896T>C 22897T>A), G446T (22898G>A 22899G>C), L452K (22916C>A 22917T>A 22918G>A), L455Y (22926T>A 22927G>T), F456L (22928T>C), K458H (22934A>C 22936G>T), S459G (22937T>G 22938C>G 22939T>C), N460K (22942T>G), K462R (22947A>G 22948A>G), T470N (22971C>A), E471V (22974A>T 22975A>G), I472P (22976A>C 22977T>C 22978C>T), Y473F (22980A>T 22981T>C), Q474S (22982C>T 22983A>C 22984G>C), A475P (22985G>C 22987C>T), G476D (22989G>A), S477G (22991A>G), T478K (22995C>A), N481T (23004A>C 23005T>C), G482P (23006G>C 23007G>C 23008T>A), V483del (23009_23011delGTT), E484P (23012G>C 23013A>C 23014A>T), G485A (23016G>C), F486L (23018T>C), F490W (23031T>G 23032T>G), Q493N (23039C>A 23041A>T), S494D (23042T>G 23043C>A 23044A>T), Q498Y (23054C>T 23056A>C), P499T (23057C>A), N501T (23064A>C), V503I (23069G>A), H519N (23117C>A), K529L (23147A>T 23148A>T 23149G>A), N532D (23156A>G 23158T>C), V534I (23162G>A), K537Q (23171A>C 23173A>G), E554P (23222G>C 23223A>C 23224G>T), N556S (23228A>T 23229A>C 23230C>A), K558R (23235A>G 23236G>A), L560Q (23241T>A 23242G>A), I569V (23267A>G), A570S (23270G>T), T572F (23276A>T 23277C>T 23278T>C), A575S (23285G>T 23287T>C), Q580K (23300C>A 23302G>A), L582S (23306C>T 23307T>C), T588S (23324A>T), S591A (23333T>G), T604A (23372A>G), N606S (23378A>T 23379A>C 23380C>T), Q607E (23381C>G 23383G>A), E619D (23419A>T), P621S (23423C>T), V622T (23426G>A 23427T>C 23428T>A), T632A (23456A>G), V635I (23465G>A 23467T>A), S640N (23480T>A 23481C>A 23482T>C), R646Q (23499G>A 23500T>A), N657D (23531A>G), N658T (23535A>C 23536C>T), Q675H (23587G>T), Q677V (23591C>G 23592A>T 23593G>T), T678S (23594A>T), N679L (23597A>T 23598A>T 23599T>A), S680_R683del (23601_23612delCTCCTCGGCGGG), A684L (23601_23612delCTCCTCGGCGGG 23613C>T), V687T (23621G>A 23622T>C 23623A>T), A688S (23624G>A 23625C>G 23626T>C), S689Q (23627A>C 23628G>A 23629T>A), Q690K (23630C>A), I693V (23639A>G 23641T>G), E702D (23668A>T), N703S (23670A>G), V705I (23675G>A), S711T (23693T>A 23695T>C), T719S (23717A>T 23719T>A), V722I (23726G>A), I726V (23738A>G 23740T>A), L727M (23741C>A 23743A>G), T732A (23756A>G 23758C>T), T739N (23778C>A 23779A>T), S750A (23810A>G 23811G>C 23812C>T), T768S (23864A>T 23866T>A), V772A (23877T>C), K776R (23888A>C 23889A>G 23890A>C), Q779R (23898A>G 23899A>T), I788M (23926T>G), P793T (23939C>A 23941A>T), I794L (23942A>T 23944T>G), D796Y (23948G>T), S810L (23990T>C 23991C>T), S813T (24000G>C 24001C>T), I834M (24064C>G), D839E (24079T>A), A845N (24095G>A 24096C>A), E868D (24166A>T), Q872A (24176C>G 24177A>C 24178A>C), S875A (24185T>G), L878V (24194T>G 24196A>T), A879S (24197G>A 24198C>G 24199G>T), I882A (24206A>G 24207T>C), S884A (24212T>G), L922Q (24326T>C 24327T>A 24328G>A), S929K (24348G>A 24349T>G), G932S (24356G>A 24358C>T), K933Q (24359A>C), D936E (24370C>A), S939T (24377T>A 24379T>A), S940T (24380T>A 24382C>A), A942S (24386G>T), S943T (24390G>C), S1055A (24725T>G), A1070S (24770G>T 24772A>C), K1073R (24780A>G), D1084E (24814T>A), H1088Y (24824C>T), S1097F (24852C>T 24853A>T), H1101S (24863C>T 24864A>C 24865C>T), V1104I (24872G>A 24874A>T), Y1110F (24891A>T), E1111S (24893G>T 24894A>C 24895A>T), V1133I (24959G>A 24961C>T), I1216V (25208A>G), M1233L (25259A>T), C1247A (25301T>G 25302G>C 25303T>A) | | | | | | | | |

| | Begin | End | Coverage | Score | Concordance | Matches | Identities | I/D/M/F* | Stop Codons |
|---|---|---|---|---|---|---|---|---|---|
| ORF3a protein (YP_009724391.1) | 1 | 276 | 100% | 1490.0 | 76.7% | 275 (99.6%) | 200 (72.5%) | 0/1/0/0 | 1 |
| | I7F (25411A>T 25413C>T), I10L (25420A>C), G11R (25423G>A), T12S (25426A>T 25428T>A), V13I (25429G>A 25431A>T), L15A (25435T>G 25436T>C 25437G>A), K16Q (25438A>C), Q17P (25442A>C), G18V (25445G>T 25446T>A), E19K (25447G>A), K21D (25453A>G 25455G>C), D22N (25456G>A), T24S (25462A>T), S26A (25468T>G), D27S (25471G>A 25472A>G), F28T (25474T>A 25475T>C), R30H (25481G>A 25482C>T), I37L (25501A>C), I47V (25531A>G), V48I (25534G>A), L52F (25546C>T), S60T (25570T>A), T64A (25582A>G 25584C>G), K66N (25590A>T), S74Y (25613C>A 25614C>T), V77F (25621G>T 25623T>C), H78Q (25626C>G), V80I (25630G>A), V90I (25660G>A 25662T>C), L101M (25693C>A 25695T>G), P104Q (25703C>A 25704T>A), V112I (25726G>A 25728C>A), S117C (25741A>T 25743T>C), F120A (25750T>G 25751T>C 25752T>A), V121C (25753G>T 25754T>G 25755A>T), L127C (25771C>T 25772T>G), R134K (25792C>A 25793G>A 25794T>A), L147V (25831C>G), N152H (25846A>C), C153N (25849T>A 25850G>A 25851T>C), S165D (25885T>G 25886C>A), S166T (25888T>A), I169V (25897A>G), S171E (25903T>G 25904C>A), T175I (25916C>T 25917A>T), T176S (25918A>T), S177T (25922G>C 25923T>A), I179K (25928T>A 25929T>A), S180L (25930T>C 25931C>T 25932T>C), E181K (25933G>A), H182E (25936C>G 25938T>A), T190S (25960A>T), K192D (25966A>G 25968A>T), W193R (25969T>A), E194H (25972G>C 25974A>C), C200Y (25991G>A), L203V (25999T>G), S205G (26005A>G 26007T>C), S209E (26017T>G 26018C>A), D210V (26021A>T 26022C>T), Y215E (26035T>G 26037C>G), L219I (26047T>A 26049G>T), S220T (26051G>C), V225I (26065G>A), H227N (26071C>A), V228A (26075T>C), Y233F (26090A>T 26091C>T), I236L (26098A>C), D238K (26104G>A 26106T>A), E239D (26109G>C), E241P (26113G>C 26114A>C 26115A>G), E242del (26116_26118delGAA), H243N (26119C>A), V256A (26159T>C), V259A (26168T>C), E261D (26175A>T) | | | | | | | | |
| envelope protein (YP_009724392.1) | 1 | 76 | 100% | 447.0 | 94.3% | 76 (98.7%) | 73 (94.8%) | 1/0/0/0 | 1 |
| | S55T (26407T>A 26409T>G), F56V (26410T>G), S68_R69insE (26448_26449insGAA), R69G (26449A>G) | | | | | | | | |
| membrane glycoprotein (YP_009724393.1) | 1 | 223 | 100% | 1419.0 | 92.9% | 222 (99.6%) | 202 (90.6%) | 0/1/0/0 | 1 |
| | S4del (26531_26533delTTC), K15Q (26565A>C 26567G>A), T30A (26610A>G 26612A>C), C33M (26619T>A 26620G>T 26621T>G), A40S (26640G>T 26642C>T), I52V (26676A>G), I76V (26748A>G 26750C>G), L87I (26781C>A), I97V (26811A>G), H125R (26896A>G 26897T>G), L129V (26907C>G), L134M (26922C>A 26924A>G), L145I (26955C>A), I151M (26975T>G), H155S (26985C>T 26986A>C 26987T>C), A188G (27085C>G 27086A>C), G189T (27087G>A 27088G>C), S197N (27112G>A 27113T>C), S211A (27153T>G), S212G (27156A>G), S214N (27163G>A 27164T>C) | | | | | | | | |
| ORF6 protein (YP_009724394.1) | 1 | 62 | 100% | 300.0 | 75.0% | 62 (100%) | 42 (67.7%) | 0/0/0/0 | 0 |
| | L16I (27247C>A 27249A>T), K23R (27269A>G 27270A>G), V24I (27271G>A), S25A (27274T>G 27276C>T), Y31V (27292T>G 27293A>T 27294C>T), N34S (27302A>G 27303C>T), L35S (27304C>T 27305T>C 27306C>A), I37V (27310A>G 27312T>G), K38R (27314A>G), N39Q (27316A>C 27318T>A), S41F (27323C>T), S43P (27328T>C 27330A>T), E46K (27337G>A), N47K (27342T>G), K48N (27345A>T), Q51E (27352C>G 27354A>G), E54D (27363A>T), Q56E (27367C>G), I60L (27379A>T 27381T>A), *62Y (27387A>T) | | | | | | | | |
| ORF7a protein (YP_009724395.1) | 1 | 122 | 100% | 758.0 | 89.6% | 122 (99.2%) | 105 (85.4%) | 1/0/0/0 | 1 |
| | A8T (27415G>A), T11V (27424A>G 27425C>T), L12F (27427C>T 27429C>T), A13T (27430G>A 27432T>A), T14S (27433A>T), S36P (27499T>C 27501T>A), F59T (27568T>A 27569T>C), Q62H (27579A>C), P68A (27595C>G), V71T (27604G>A 27605T>C 27606A>T), K72R (27607A>C 27608A>G), V74T (27613G>A 27614T>C), Q94_E95insQ (27675_27676insCAA), I100L (27691A>C), I107L (27712A>C), I110L (27721A>T), T111I (27725C>T), L116I (27739C>A 27741C>T) | | | | | | | | |
| ORF7b (YP_009725296.1) | 1 | 44 | 100% | 252.0 | 79.7% | 44 (97.8%) | 36 (80.0%) | 1/0/0/0 | 1 |
| | I2N (27760T>A), S5T (27768T>A 27770A>T), L34I (27855C>A 27857G>C), H37L (27865A>T 27866T>A), N38E (27867A>G 27869T>A), T40P (27873A>C), H42T (27879C>A 27880A>C), H42_A43insK (27881_27882insAAA), A43V (27883C>T) | | | | | | | | |
| ORF8 protein (YP_009724396.1) | 1 | 120 | 99.2% | 117.0 | 15.6% | 104 (82.5%) | 37 (29.4%) | 5/17/1/1 | 0 |
| | F3L (27900T>C), V5I (27906G>A), F6V (27909T>G 27911C>T), G8_I10del (27915_27923delGGAATCATC), T12C (27927A>T 27928C>G), V13I (27930G>A 27932A>T), A14S (27933G>T), A15L (27936G>C 27937C>T), F16C (27940T>G 27941T>C), H17S (27942C>A 27943A>G 27944C>T), Q18C (27945C>T 27946A>G 27947A>C), E19I (27948G>A 27949A>T), S21T (27955G>C), L22V (27957T>G), L22_Q23insV (27959_27960insGTA), S24R (27963T>C 27964C>G 27965A>C), T26A (27969A>G 27971T>A), Q27S (27972C>T 27973A>C 27974A>T), H28N (27975C>A), Q29K (27978C>A), Y31H (27984T>C), V33L (27990G>C), D34E (27995T>A), P38_S43del (28005_28024delCCTATTCACTTCTATTCTAA), K44X (28005_28024delCCTATTCACTTCTATTCTAA), W45R (28026T>A), I47N (28033T>A 28034T>C), R48T (28036G>C 28037A>T), V49R (28038G>A 28039T>G 28040A>G), A51N (28044G>A 28045C>A), R52T (28048G>C 28049A>T), K53Y (28050A>T 28052A>T), A55T (28056G>A 28058A>T), P56A (28059C>G), L57W (28063T>G 28064A>G), I58L (28065A>C), E59_L60del (28068_28073delGAATTG), V62A (28078T>C 28079G>T), D63L (28080G>C 28081A>T 28082T>A), E64_A65del (28083_28088delGAGGCT), S67K (28092T>A 28093C>A 28094T>G), K68V (28095A>G 28096A>T 28097A>T), S69L (28099C>T), I71F (28104A>T), Q72H (28109G>T), Y73R (28110T>A 28111A>G 28112C>A), I74W (28113A>T 28114T>G 28115C>G), D75H (28116G>C 28118T>C), I76T (28120T>C 28121C>T), G77M (28122G>A 28123G>T 28124T>G), N78V (28125A>G 28126A>T), Y79Q (28128T>C 28130T>A), V81_S82del (28134_28139delGTTTCC), L84T (28143T>A 28144T>C), P85_F86insN (28148_28149insAAT), F86V (28149T>G), E92D (28169A>T), K94A (28173A>G 28174A>C 28175A>T), L95G (28176T>G 28177T>G 28178G>T), S97A (28182A>G 28183G>C 28184T>G), V99I (28188G>A), V100A (28192T>C 28193G>T), S103W (28201C>G), F104Y (28204T>A), Y105L (28206T>C 28207A>T), E106H (28209G>C 28211A>T), D107E (28214C>A), F108G (28215T>G 28216T>G), L109H (28218T>C 28219T>A 28220A>C), E110Q (28221G>C 28223G>A), E110_Y111insTAA (28223_28224insACTGCTGCA), Y111F (28225A>T), H112R (28227C>A 28228A>G 28229T>A), R115L (28237G>T), D119N (28248G>A), F120K (28251T>A 28252T>A 28253C>A) | | | | | | | | |
| nucleocapsid phosphoprotein (YP_009724397.2) | 1 | 420 | 100% | 2641.0 | 92.1% | 420 (99.3%) | 382 (90.3%) | 3/0/0/0 | 1 |
| | Q7_N8insS (28294_28295insTCA), N11S (28305A>G), S21T (28334T>A), G25D (28347G>A), S26N (28350G>A), E31G (28365A>G), S33N (28371G>A), S37P (28382T>C), D63E (28462C>A), K65R (28467A>G), S79G (28508A>G), I94V (28553A>G), D103E (28582T>G), G120S (28631G>T 28632G>C), D128E (28657C>A), I131V (28664A>G), A152N (28727G>A 28728C>A), I157T (28743T>C), N192G (28847A>G 28848A>G 28849C>T), S193N (28851G>A), T205N (28887C>A), G212S (28907G>A), N213G (28910A>G 28911A>G 28912T>A), D216E (28921T>A), A217T (28922G>A), M234V (28973A>G 28975G>T), A267Q (29072G>C 29073C>A 29074A>G), E290D (29143A>C), T334H (29273A>C 29274C>A 29275A>T), N345Q (29306A>C 29308T>A), Q349N (29318C>A 29320A>C), A376T (29399G>A), T379A (29408A>G), A381P (29414G>C 29416C>T), Q390P (29442A>C 29443A>C), L400M (29471T>A), K405R (29487A>G), Q409N (29498C>A 29500A>T), S413G (29510A>G 29511_29512insAGCTTC), S413_A414insAS (29511_29512insAGCTTC) | | | | | | | | |
| ORF10 protein (YP_009725255.1) | 1 | 39 | 100% | 212.0 | 78.8% | 39 (100%) | 32 (82.1%) | 0/0/0/0 | 2 |
| | I4V (29567A>G), F9I (29582T>A), Y14H (29597T>C), Y26* (29635C>A), I27T (29637T>C), D31G (29649A>G), V32L (29651G>T) | | | | | | | | |

*: Inserts / Deletes / Misaligned / Frameshifts
